# The evolution and developmental dynamics of histone-based chromatin regulation in Annelida

**DOI:** 10.1101/2024.09.20.614060

**Authors:** Francisco M. Martín-Zamora, Joby Cole, Rory D. Donnellan, Kero Guynes, Allan M. Carrillo-Baltodano, Mark J. Dickman, Paul J. Hurd, José M. Martín-Durán

**Author notes:** Correspondence: José M. Martín-Durán, Francisco M. Martín-Zamora.

## Abstract

Eukaryotic histones protect and package nuclear DNA into nucleosomes. The dynamic addition and removal of posttranslational modifications on these proteins define regulatory regions that play a central role in genome and chromatin biology. However, our understanding of these regulatory mechanisms in animals is largely based on a few model systems, which prevents a general understanding of how histone-based regulation unfolds and promotes phenotypic variation during animal embryogenesis. Here, we apply a comprehensive multi-omics approach to dissect the histone-based regulatory complement in Annelida, one of the largest invertebrate phyla. Annelids exhibit a conserved histone repertoire organised in clusters of dynamically regulated, hyperaccessible chromatin. However, unlike other animals with reduced genomes, the worm *Dimorphilus gyrociliatus* shows a dramatically streamlined histone repertoire, revealing that genome compaction has lineage-specific effects on histone-based regulation. Notably, the annelid *Owenia fusiformis* has two H2A.X variants that co-occur in other animals, whose functional implications are unclear but represent a unique case of widespread parallel evolution of a histone variant in Eukarya. Histone-modifying enzyme complements are largely conserved amongst annelids. Yet, temporal differences in the expression of a reduced set of histone modifiers correlate with distinct ontogenetic traits and variation in the adult landscapes of histone modifications, as revealed by quantitative mass spectrometry in *O. fusiformis* and *Capitella teleta*. Collectively, our unparalleled analysis of histone-based epigenetics within a non-model phylum informs the evolution of histone-based regulation, presenting a framework to explore how this fundamental genome regulatory layer contributes to developmental and morphological diversification in annelids and animals generally.

## Introduction

Genome packaging is a critical phenomenon in cell biology. Different mechanisms, like DNA supercoiling, aid in this process (Gilbert and Allan 2014), yet the packaging in certain viruses (Bryson et al. 2022; Irwin and Richards 2024), bacteria (Hocher et al. 2023), archaea (Pereira et al. 1997), and especially in eukaryotes (Khorasanizadeh 2004; Abouelmagd and Ageely 2013; Nelson et al. 2021), results from the interaction of genomic DNA with histones. Eukaryotic histones are highly basic, DNA-associated proteins that protect and package nuclear, naked DNA to arrange them into higher-order and densely compacted structures, all the way up to the chromosome level. By doing so, they play a central role as regulators of genome architecture and almost all DNA-based biological processes, especially the regulation of gene expression (Khorasanizadeh 2004). Histone-based regulation is, indeed, one of the most versatile and logically intricate mechanisms modulating genome functioning (Lee et al. 2010; Bannister and Kouzarides 2011; Millán-Zambrano et al. 2022) (Fig. 1A). These mechanism include histone dynamics and differential usage of histone variants (Henikoff and Smith 2015; Buschbeck and Hake 2017; Ferrand et al. 2020). Yet, arguably, the most functionally important regulatory mechanism involves the deposition and removal of small covalent posttranslational modifications, such as methyl and acetyl moieties, to key residues primarily located in the N-terminal end, or tail, of histones (Kimura 2013; Skvortsova et al. 2018). Certain histone posttranslational modifications (hPTMs) cause structural perturbations in the nucleosome structure, thereby changing the level of chromatin condensation or accessibility. Others act as binding marks for modules of reader proteins and complexes that recruit chromatin-modifying machinery, leading to different biological readouts (Workman and Kingston 1998; Bannister and Kouzarides 2011; Yun et al. 2011; Tessarz and Kouzarides 2014). In this way, hPTMs define regulatory regions in the genome and their levels of regulatory activity, constituting critical regulators of various processes like proliferation, differentiation, and most importantly, embryogenesis and development (Chi et al. 2010; Audia and Campbell 2016; Jambhekar et al. 2019; Völker-Albert et al. 2020; Millán-Zambrano et al. 2022). Even though the central histone-based regulation mechanisms are largely evolutionarily conserved, their vast regulatory activity also makes them a very promising source of evolutionary variation.

**Figure 1.**
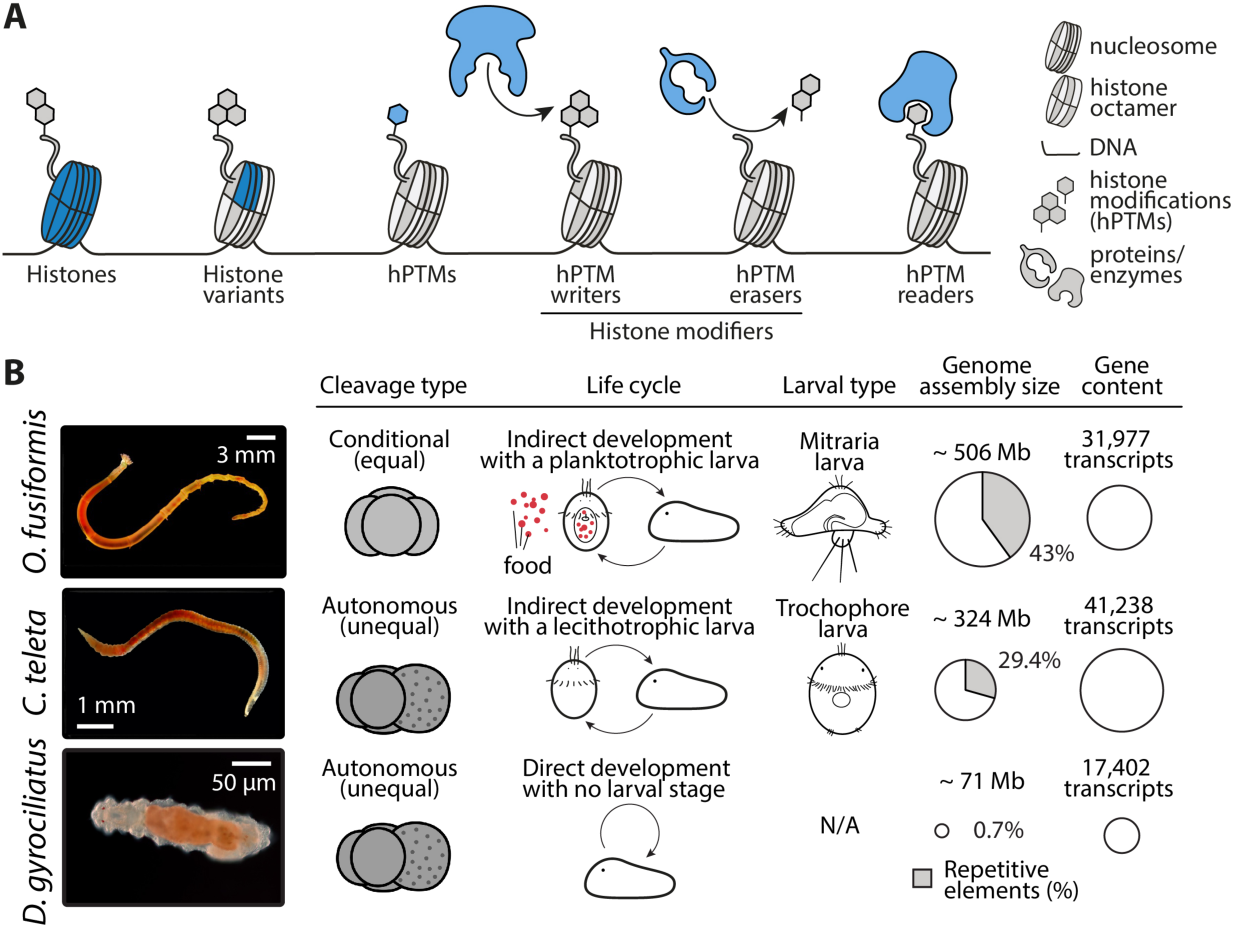
Histone-based regulation of three annelid species. (**A**) Schematic drawing depicting the potential sources of evolutionary variation in the histone-based regulation complement. Drawings are not to scale. (**B**) *O. fusiformis*, *C. teleta*, and *D. gyrociliatus* possess a very conserved early embryogenesis programme yet display key differences. From left to right, cleavage type, life cycle, larval type, genome assembly size and repetitive elements percentage, and gene content are displayed for each species. While *D. gyrociliatus* and *C. teleta* are unequal cleavers and commit cell fates during the first cell divisions, *O. fusiformis* is a conditional cleaver that does not do so until later cleavage stages. *O. fusiformis* possesses the mitraria larva, a feeding, i.e., planktotrophic, larva, while *C. teleta* is an example of a species with a classic non-feeding or lecithotrophic trochophore larva, and *D. gyrociliatus* lacks a larval stage. *D. gyrociliatus* has a very compact genome, almost devoid of repetitive elements, that has reduced its size and gene content conservatively. Images are for adult specimens of each species. Bubble plots are proportional to genome size and gene content. hPTMs: histone posttranslational modifications.

Little is known about the repertoire and role of hPTMs outside traditional animal models for which high-quality reference genomes and methods to interrogate histone regulation have been readily available. In Spiralia, one of the three largest clades of bilaterally symmetrical animals comprising morphologically diverse lineages and nearly half of the animal phyla (Martín-Durán and Marlétaz 2020), functional genomics work has been limited (Martín-Zamora et al. 2023a). Research on histones peaked during the second half of the 20^th^ century. Those early works were primarily limited to biochemical studies on variations in histone sequence, structure and composition across a handful of major groups, namely molluscs (van Helden et al. 1979; Wouters-Tyrou et al. 1982; Sellos 1985; Ausio and Van Holde 1988), annelids (Kmiecik et al. 1983; Sellos et al. 1990; Davis et al. 1992), and nemerteans (Wang and Ausió 2001). Likewise, hPTM-targeting functional genomics studies with ChIP-seq (Johnson et al. 2007; Kasinathan et al. 2014) and more modern techniques (Schmidl et al. 2015; Skene and Henikoff 2017; Kaya-Okur et al. 2019; Meers et al. 2019; Kaya-Okur et al. 2020) have so far been restricted to Platyhelminthes, where efforts have focused on understanding stem cell maintenance and regeneration in the planarian *Schmidtea mediterranea* (Rink 2013; Duncan et al. 2015; Dattani et al. 2018; Mihaylova et al. 2018; Blanco et al. 2020; Li et al. 2021; Verma et al. 2021; Neiro et al. 2022; Pascual-Carreras et al. 2023) and the life cycle of the parasitic helminth *Schistosoma mansoni* to develop potential treatments to schistosomiasis, the disease it causes (Cabezas-Cruz et al. 2014; Liu 2016; Picard et al. 2016; Roquis et al. 2016; Anderson et al. 2017; Cosseau et al. 2017; Roquis et al. 2018; Picard et al. 2019; Ghazy et al. 2022). Therefore, our knowledge of histone-based regulation in non-model systems such as spiralians is quite limited, thus hampering a comprehensive understanding of how changes in genome regulation promotes variation in gene expression programmes, and ultimately phenotypic change, during animal evolution.

Variations in the timing or rate of development, commonly called heterochronies, are a key source of evolutionary variation (McNamara 2012; Keyte and Smith 2014; Buendía-Monreal and Gillmor 2018; Dobreva et al. 2022). In annelids, one of the largest spiralian clades, and likely in bilaterians generally (Degnan and Degnan 2023; Formery and Lowe 2023; Martín-Zamora et al. 2023b), developmental heterochronies correlate with the diversification of embryonic outcomes and life cycles. The annelids *Owenia fusiformis* Delle Chiaje, 1844; *Capitella teleta* Blake, Grassle & Eckelbarger, 2009; and *Dimorphilus gyrociliatus* (O. Schmidt, 1857) share a common early developmental programme—spiral cleavage—but have distinct strategies for cell fate specification, life cycles, and larval types (Fig. 1B) (Seaver et al. 2005; Kerbl et al. 2016a; Carrillo-Baltodano et al. 2021). In these annelids, genes involved in key cellular processes (e.g., autophagy) and enzymatic pathways (e.g., chitin synthesis pathway) are either pre- or post-displaced between larval types, i.e., they have an earlier or later expression onset, respectively (Martín-Zamora et al. 2023b). Moreover, the trunk patterning transcriptional programme, which includes the expression of the *Hox* genes, is progressively brought forward in development—i.e., pre-displaced—as larval traits are lost and indirect development transitions into direct development (Martín-Zamora et al. 2023b). Importantly, these changes in gene expression mirror chromatin accessibility dynamics, as observed in the open chromatin regions of the *Hox* cluster and the transcription factor binding dynamics of HOX DNA-binding motifs (Martín-Zamora et al. 2023b).Therefore, given their fundamental role in regulating gene expression, differences in the dynamics of histone-based regulation—particularly of hPTMs—may be upstream and account for these heterochronies, providing a mechanistic explanation of how changes in genome regulation trigger adaptive variation in developmental programmes.

Here, we begin to tackle the relationship between histone-based regulation and developmental transcriptional programmes by mining and characterising the histone complement and histone modification machinery in the annelids *O. fusiformis*, *C. teleta*, and *D. gyrociliatus* through a large-scale multi-omics profiling at the genomic, transcriptomic, epigenomic, and proteomic levels. We describe a minimal histone complement in the miniature genome of *D. gyrociliatus* and identify divergence in a key histone variant throughout Eukarya that may account for the evolution of early embryonic phenotypes in animals. Despite their differences in histone numbers, the repertoires of histone modifiers are largely complete and conserved between these annelids, although some families exhibit expansions and domain fusions that hint at neofunctionalization events. Concomitant to the heterochronies in other gene regulatory networks and pathways, some histone modifiers are shifted in their developmental time of expression between these annelids, correlating with their life cycle and larval type differences, as well as with the levels of hPTMs in the adults. Altogether, our study is the most comprehensive comparative characterisation of histone-based regulation in a spiralian clade, paving the way for the genome-wide profiling of hPTMs in annelids and the functional assessment of the interplay between histone modifications and development programmes in the phenotypic diversification of this major animal group.

## Results and Discussion

### Annelids exhibit a conserved histone repertoire

To investigate the histone gene repertoire of spiralians and annelids, we first reannotated the H2A, H2B, H3 and H4 histone gene models by combining a protein-to-genome alignment-based approach, transcriptomic developmental time series data, a careful manual curation, and an orthology assignment based on maximum likelihood and Bayesian reconstructions (Fig. 2A; Supplementary Fig. 1, 2). We recovered 81, 90, and 17 core histone genes for *O. fusiformis*, *C. teleta*, and *D. gyrociliatus*, respectively (Table 1). In these species, only six, three, and four genes, respectively, displayed an atypical divergent amino acid sequence, which we regard as unknown histone variants. Out of these genes with unknown orthology, all six in *O. fusiformis* and one in *C. teleta* were deemed pseudogenic due to having either null or negligible low expression levels throughout development (Supplementary Fig. 7E–G, 8C). Unlike in other protostomes, where certain histone variants have been lost, like H2A.X in Nematoda (Lemmens and Tijsterman 2011), or where features and roles from multiple variants get merged into a single gene, like H2A.X and H2A.Z in the H2A.v variant in *Drosophila* (Baldi and Becker 2013), all three analysed annelid taxa have conserved at least one ortholog of each of the core histone proteins—canonical and variant—presumed to be ancestral to Bilateria (Henikoff and Smith 2015; Grau-Bové et al. 2022), i.e., the canonical H2A, H2B, H3, and H4 proteins, and the H2A.X, H2A.Z (or H2Av), macroH2A, H3.3, and cenH3 histone variants, suggesting a potential functional conservation of the histone complement. *Owenia fusiformis* and *C. teleta* possess between 16 and 23 copies per canonical histone (Table 1). However, the compact genome of *D. gyrociliatus* only encodes two genes per canonical histone (Table 1), thus making this fully conserved histone complement one with the lowest (if not the lowest) described copy numbers for canonical histones in a metazoan lineage.

**Figure 2.**
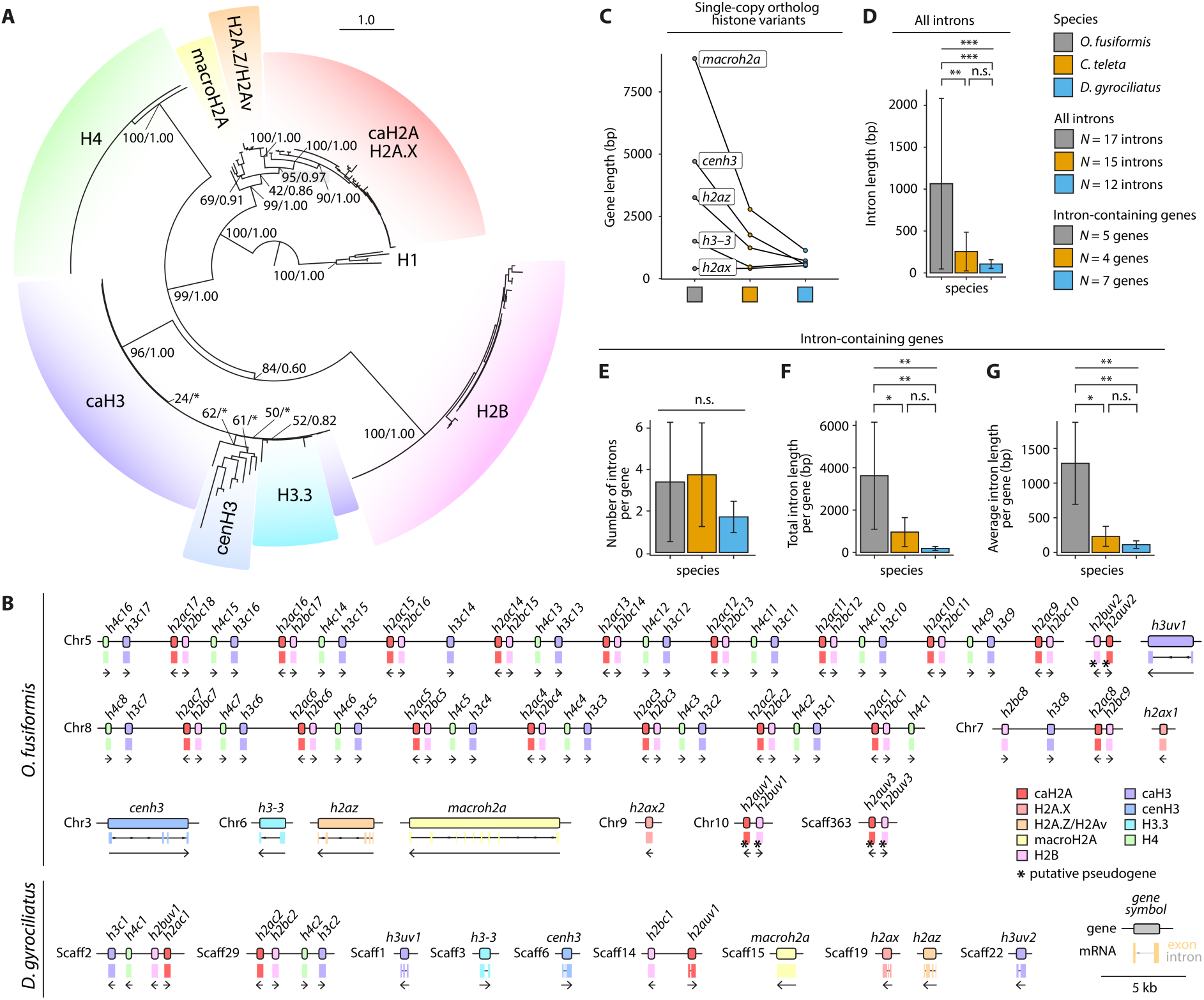
Annelids exhibit a conserved histone repertoire. (**A**) Summary phylogeny of the gene orthology analysis of histone genes in *O. fusiformis*, *C. teleta*, and *D. gyrociliatus*. Depicted tree topology is based on the maximum likelihood reconstruction. Branch support values represent bootstrap values (0–100 values) and posterior probabilities (0–1 values) at key nodes. Coloured boxes highlight the extent of each histone gene or family. Clades supported by maximum likelihood only are flagged with an asterisk (*). The scale bar depicts the number of amino acid changes per site along the branches. caH2A: canonical H2A; caH3: canonical H3. (**B**) Schematic representation to scale of the genomic loci of the histone genes in *O. fusiformis* (in the chromosome-level assembly) and *D. gyrociliatus*. Boxes delimitate gene bodies, with the intron-exon composition shown underneath. Arrows below genes indicate the direction of transcription. Colours correspond to the different histone genes and gene families. Genes flagged with an asterisk (*) represent putative pseudogenes, as inferred from transcriptomic data (see Supplementary Fig. 7E–G, 8C). Chr: chromosome; Scaff: scaffold. (**C**) Gene length of the histone variants with inferable orthology across all three annelid taxa, shown as paired data points. (**D**) Intron length of all introns contained in histone genes. (**E–G**) Gene-wise number of introns (**E**), total intron length (**F**), and average intron length (**G**) in all intron-containing histone genes across all three worm species. Error bars in **D–G** are standard deviations. *P* values were derived using one-way ANOVAs, followed by two-tailed post-hoc Tukey tests for pair-wise comparisons, wherever applicable. *: *P* value < 0.05; **: *P* value < 0.01; *P* value < 0.001; n.s.: not significant.

**Table 1.** Histone gene repertoire in Annelida. Total number of canonical and variant histones encoded in the genomes of *O. fusiformis*, *C. teleta*, and *D. gyrociliatus*, classified by family (H2A, H2B, H3, and H4). For each family, the different canonical and variant histones are shown. Numbers in parentheses denote the number of non-pseudogenic genes.

| Protein | *O. fusiformis* | *C. teleta* | *D. gyrociliatus* |
| --- | --- | --- | --- |
| **H2A family** | 24 (21) | 23 | 6 |
| H2A canonical | 17 | 20 | 2 |
| H2A.X | 2 | 1 | 1 |
| H2A.Z | 1 | 1 | 1 |
| macro-H2A | 1 | 1 | 1 |
| H2A unknown variants | 3 (0) | 0 | 1 |
| **H2B family** | 21 (18) | 19 | 3 |
| H2B canonical | 18 | 18 | 2 |
| H2B unknown variants | 3 (0) | 1 | 1 |
| **H3 family** | 20 | 25 (24) | 6 |
| H3 canonical | 17 | 22 | 2 |
| H3.3 | 1 | 1 | 1 |
| cenH3 | 1 | 1 | 1 |
| H3 unknown variants | 1 | 2 (1) | 2 |
| **H4 family** | 16 | 23 | 2 |
| H4 canonical | 16 | 23 | 2 |
| **Total** | 81 (75) | 90 (89) | 17 |

### Genomic organisation and gene structure of histones

Canonical histones tend to appear in tandemly repeated organised clusters, sometimes even associated with H1 linker histones (Nagel and Grossbach 2000; Buschbeck and Hake 2017; Amatori et al. 2021). To assess whether this also occurs in annelids, we characterised the genomic organisation of the core histone genes in *O. fusiformis* and *D. gyrociliatus*, the two focal species with more contiguous reference genomes (Fig. 2B; Supplementary Fig. 3A). In *O. fusiformis*, canonical core histones are arranged into two large clusters containing 35 and 29 canonical histones in chromosomes 5 and 8, respectively, and a smaller one containing four genes only in chromosome 7. With the exceptions of the chromosome 7 cluster and a single occurrence in the chromosome 5 cluster, core canonical histones are tandemly arrayed as a H4– H3–H2A–H2B repeating unit. In this unit, H4, H3, and H2B are transcribed from one strand, and H2A is transcribed from the opposite one (Fig. 2B, Supplementary Fig. 3A). Concomitant to the extremely reduced histone copy number, the genomic clusters in *D. gyrociliatus* are minimal in size, consisting of a single H3–H4–H2B–H2A repeating unit in scaffolds 2 and 29 (Fig. 2B). This may result from a genomic translocation of the H2A–H2B and H3–H4 units, with the transcripts still being synthesised from the same strands as in *O. fusiformis*. The presence of these H2A–H2B and H3–H4 units makes it plausible that previously described head-to-head gene pairs in *C. teleta* and *Helobdella robusta* (Grau-Bové et al. 2022) are also found in *O. fusiformis* and *D. gyrociliatus* and potentially in other annelids like *Chaetopterus variopedatus* (del Gaudio et al. 1998). Interestingly, *Owenia fusiformis* and *D. gyrociliatus* display different genomic organisations than those reported for the annelids *Platynereis dumerilii* (Sellos et al. 1990; Krawetz et al. 1993) and *Urechis capo* (Davis et al. 1992). Although these are not based on high-quality reference genome analyses and could be reconciled with a single translocation event from the putative *O. fusiformis* state, they might also indicate that different genomic organisations of the histone complement occurred in annelids.

All clustered histone genes lack introns in both species, whereas histone variants are often multi-exonic (Fig. 2B, Supplementary Fig. 3A). Interestingly, differences in the overall gene length between species are not statistically significant (Fig. 2C; Supplementary Fig. 3B, C). However, the overall intron length of histones, as well as the total and average intron length per gene of intron-containing genes, but not the number of introns per gene, are significantly lower in both *C. teleta* and *D. gyrociliatus* compared to *O. fusiformis* (Fig. 2D–G; Supplementary Fig. 3D–F). This reduction of average intron length can even be traced gene by gene in the single-copy histone variant genes *h2ax*, *h2az*, *macroh2a*, *cenh3*, and *h3–3* (Supplementary Fig. 3G–I). Therefore, annelids organise their histone genes in tandemly repeated organised clusters. Still, significant changes have occurred in both the cluster structure and number, as well as the gene structure, throughout annelid diversification.

### Histone genes are in dynamic hyperaccessible chromatin regions

To better understand chromatin regulation around histone genes and their genomic clusters, we leveraged our publicly available ATAC-seq developmental time series for *O. fusiformis* and *C. teleta* (Martín-Zamora et al. 2023b), as well as data for *D. gyrociliatus* female adults (Martín-Durán et al. 2021). Histone genes are amongst those with the highest ATAC-seq enrichment at every developmental point in *O. fusiformis* and *C. teleta*, albeit more starkly in the former (Fig. 3A, B, Supplementary Fig. 4A–D). In these species, open chromatin is most prevalent at the transcription start sites and promoter regions, progressively decreasing along the gene body towards the transcription end sites. During the embryogenesis of *O. fusiformis* and *C. teleta*, chromatin around histone genes becomes more accessible after the blastula stage during gastrulation, reaching its peak openness and remaining at very elevated levels during larval development (Fig. 3A, B, Supplementary Fig. 4A–D). At the end of the developmental time course, however, chromatin around the histone clusters becomes more compact. This above-average enrichment is present in the clusters that contain canonical histones—especially in the large clusters of *O. fusiformis*, where it is very significant—and less so in the loci of the variant histone genes (Fig. 3C, D, Supplementary Fig. 4C, D, 5A–C, 6A, B). In *D. gyrociliatus*, histone genes are also in loci of highly open chromatin (Supplementary Fig. 4E, F). However, their chromatin accessibility landscape differs from those of *O. fusiformis* and *C. teleta*, with chromatin being open almost exclusively at the TSS in defined peaks (Supplementary Fig. 5D, 6C). Therefore, histones are in dynamically regulated, hyperaccessible chromatin regions in annelids, suggesting their expression might be more variable than expected for genes essential for DNA compaction and regulation.

**Figure 3.**
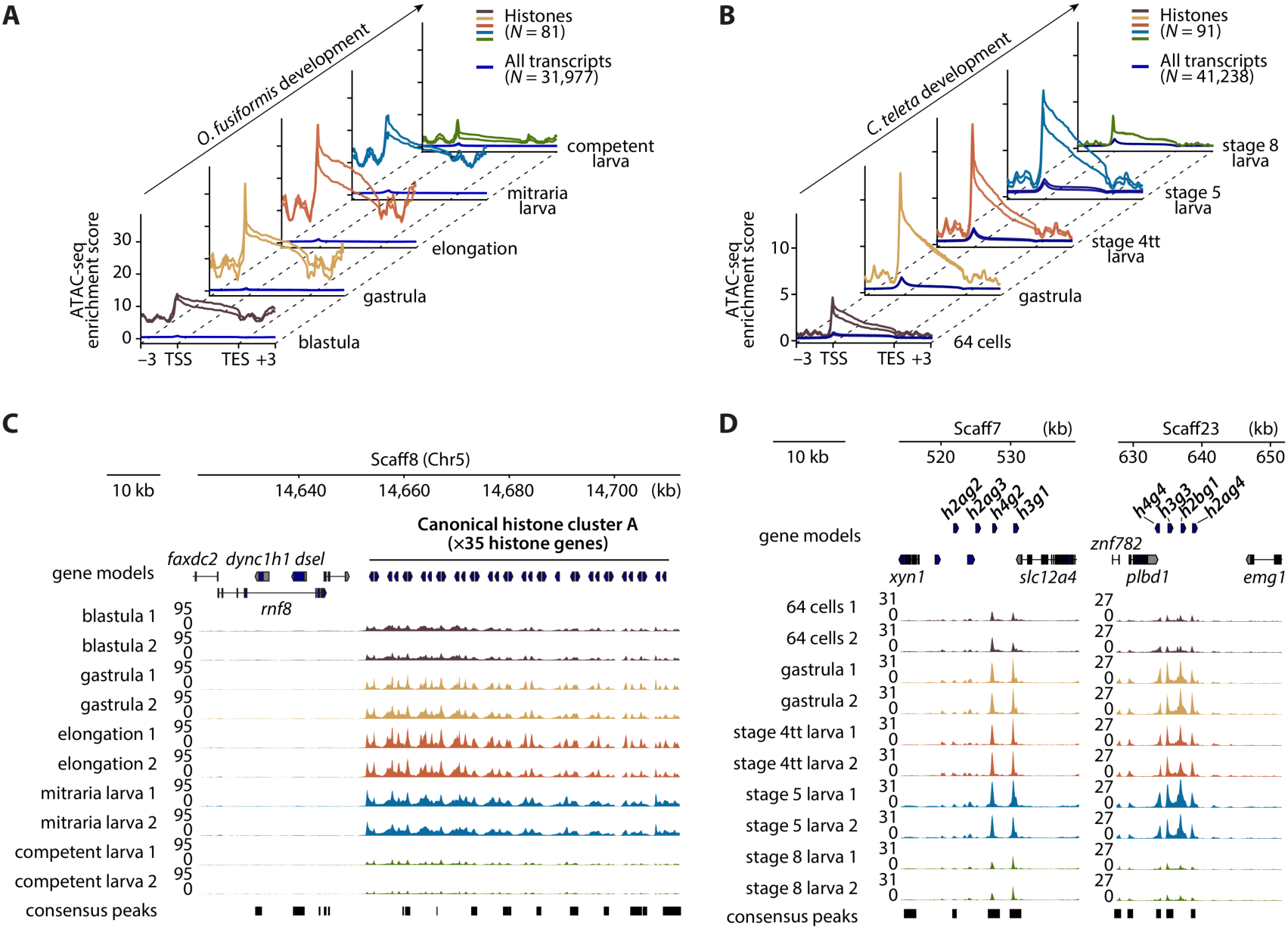
Histone genes are in dynamic hyperaccessible chromatin regions. (**A**, **B**) ATAC-seq enrichment meta-gene profiles of histone genes compared to the whole genome during the embryonic development of *O. fusiformis* (**A**) and *C. teleta* (**B**). Distances are in kilobases (kb). TSS: transcription start site; TES: transcription end site. *dsel*: dermatan sulphate epimerase like; *dync1h1*: dynein cytoplasmic 1 heavy chain 1; *emg1*: EMG1 N1-specific pseudouridine methyltransferase, *faxdc2*: fatty acid hydroxylase domain containing 2; *plbd1*: phospholipase B domain containing 1; *rnf8*: ring finger protein 8; *slc12a4*: solute carrier family 12 member 4; *xyn1*: xylanase 1; *znf782*: zinc finger protein 782. (**C**) ATAC-seq tracks at one of the two large clusters of canonical histones of *O. fusiformis*, located in chromosome 5. (**D**) ATAC-seq tracks at two of the six histone gene clusters of *C. teleta* that harbour at least four histones. Note how in both plots the ATAC-seq signal is so high for histone genes that the peaks called in neighbouring genes are barely visible at the plot scale. Chr: chromosome; Scaff: scaffold.

### Histone gene expression dynamics in Annelida

To investigate histone gene expression dynamics across annelid development, we mapped the developmental RNA-seq time courses for all three species (Burns and Pechenik 2017; Martín-Durán et al. 2021; Martín-Zamora et al. 2023b) against the new gene models containing the curated histone genes. Canonical histone gene expression levels were calculated as the sum of the expression of all the genes encoding the same protein (e.g., all 17 genes for caH2A in *O. fusiformis*). All four canonical histones (caH2A, caH2B, caH3, and caH4) present similar expression patterns within species (Supplementary Fig. 7A–D, 8A). Moreover, in both species with a larval stage, i.e., *O. fusiformis* and *C. teleta*, histones are expressed in the zygote and cleavage stages—particularly in *O. fusiformis*—but reach maximum expression levels after the zygotic genome activation, at the gastrula stage and immediate post-gastrula stages. Thereafter, they progressively decrease until the competent larva and juvenile stages (Supplementary Fig. 7A–C, 8A). This mirrors and correlates with the developmental dynamics of the accessible chromatin regions in which they are located. In the direct developer *D. gyrociliatus*, however, maximum expression of canonical histones occurs at the early embryogenesis time point and plummets immediately after (Supplementary Fig. 7A, D, 8A). More interesting are the expression patterns of histone variants, whose expression dynamics across the development of all three studied annelids are conserved, thereby hinting possible conservation of their roles in development (Supplementary Fig. 7A, E–G, Supplementary Fig. 8B). The *h2az* histone, which is known to maintain the pluripotency state of stem cells in development (Giaimo et al. 2019; Dijkwel and Tremethick 2022), is expressed at high levels in cleavage and early embryogenesis, and it decreases after gastrulation when cell fates are acquired and terminal differentiation into multiple cell lineages starts. Very similar is the pattern of *h3–3*, which incorporates into chromatin to sustain several differentiation programmes across metazoans (Sakai et al. 2009; Bush et al. 2013; Jang et al. 2015). The centromeric *cenh3*, which is critical in kinetochore positioning and assembly during cell division (Quénet and Dalal 2012; Kixmoeller et al. 2020), is also highly expressed at cleavage stages when cells are dividing quickly, but its levels dilute earlier and decrease even before the blastula forms, unlike *h3–3* and *h2az*. The only case in which expression increases dramatically throughout development to reach a maximum in the adult is the *macroh2a* gene, which might be biologically relevant given its known role in transcriptional repression in maintaining the identity of terminally differentiated cells (Buschbeck et al. 2009; Gaspar-Maia et al. 2013). We also found distinct expression patterns of the unknown/unidentified histone variants across all species (Supplementary Fig. 7A, H–J, 8C), particularly that of *h3uv1* in *O. fusiformis* and *h2buv1* in *C. teleta*, which appear to be extremely abundant genes. We systematically explored the conservation of known hPTMs sites in these genes and all other mined histone genes (see Data Availability section), but we still we have no hints of their functional relevance.

### Histone H2A.X variants have evolved in parallel in Metazoa

The histone variant H2A.X, critical in DNA repair (Rogakou et al. 1999; Shroff et al. 2004) and embryonic stem cell development (Andäng et al. 2008; Wu et al. 2014), among other non-canonical functions (Turinetto and Giachino 2015), is strikingly divergent across the studied annelids. While other variants have a single ortholog in all three species, *C. teleta* and *D. gyrociliatus* display a single *h2ax* gene, but *O. fusiformis* encodes two different H2A.X proteins that share an 88.1 % sequence identity (Fig. 2A, B; Supplementary Fig. 1, 2, 3A; Table 1). Both paralogs are maternally deposited in the egg and highly abundant in early embryonic stages, and progressively lose expression after the blastula and gastrula stages up to their minima in the juvenile adult stage (Fig. 4A; Supplementary Fig. 8B). The same expression dynamics are evident for *C. teleta* and *D. gyrociliatus* (Fig. 4A, Supplementary Fig. 8B). Nevertheless, while developmental dynamics of both paralogs in *O. fusiformis* are similar, the expression of *h2ax2* is up to 150-fold greater than that of *h2ax1*, confirming that *h2ax2* is the dominant H2A.X gene (Fig. 4A, Supplementary Fig. 7E). Yet, despite the lower levels of expression, we could confidently prove *h2ax1* is expressed throughout the life cycle of *O. fusiformis* by specifically amplifying both genes from a cDNA pool of mRNA coming from all developmental time points of *O. fusiformis* (Supplementary Fig. 9). Remarkably, some of the differences in protein sequence between *h2ax1* and *h2ax2* lie in critical regions, most strikingly in the C-terminal motif of the protein (Fig. 4B, C; Supplementary Fig. 10A). In mammals and most chordates, Tyr142 is a conserved position that undergoes biologically relevant post-translational modifications (Cook et al. 2009; Xiao et al. 2009). Where *h2ax1* displays this phosphorylatable Tyr142 (from here on also H2A.X-Y), *h2ax2* has a sterically homologous yet unmodifiable Phe142 (thus being called H2A.X-F) (Fig. 4B). Indeed, this residue change has been observed before in the H2A.X-F gene of *Xenopus,* where it is expressed in eggs and early embryos and where it has been hypothesised to facilitate rapid early-embryo cell divisions (Shechter et al. 2009; Wang et al. 2014), but also in some fish (Shechter et al. 2009), a mollusc (González-Romero et al. 2012), and in plants (Shechter et al. 2009; Lorković et al. 2017; Lei and Berger 2020). Indeed, we found that some of the few curated spiralian H2A.X sequences—and also some arthropod and other eukaryote ones— display this change (Fig. 4D; Supplementary Fig. 10B–F). Therefore, the distribution of H2A.X-Y and H2A.X-F variants is more widespread than previously foreseen. Yet, it is unknown whether these originated in the last common eukaryotic ancestor or are the result of parallel evolution.

**Figure 4.**
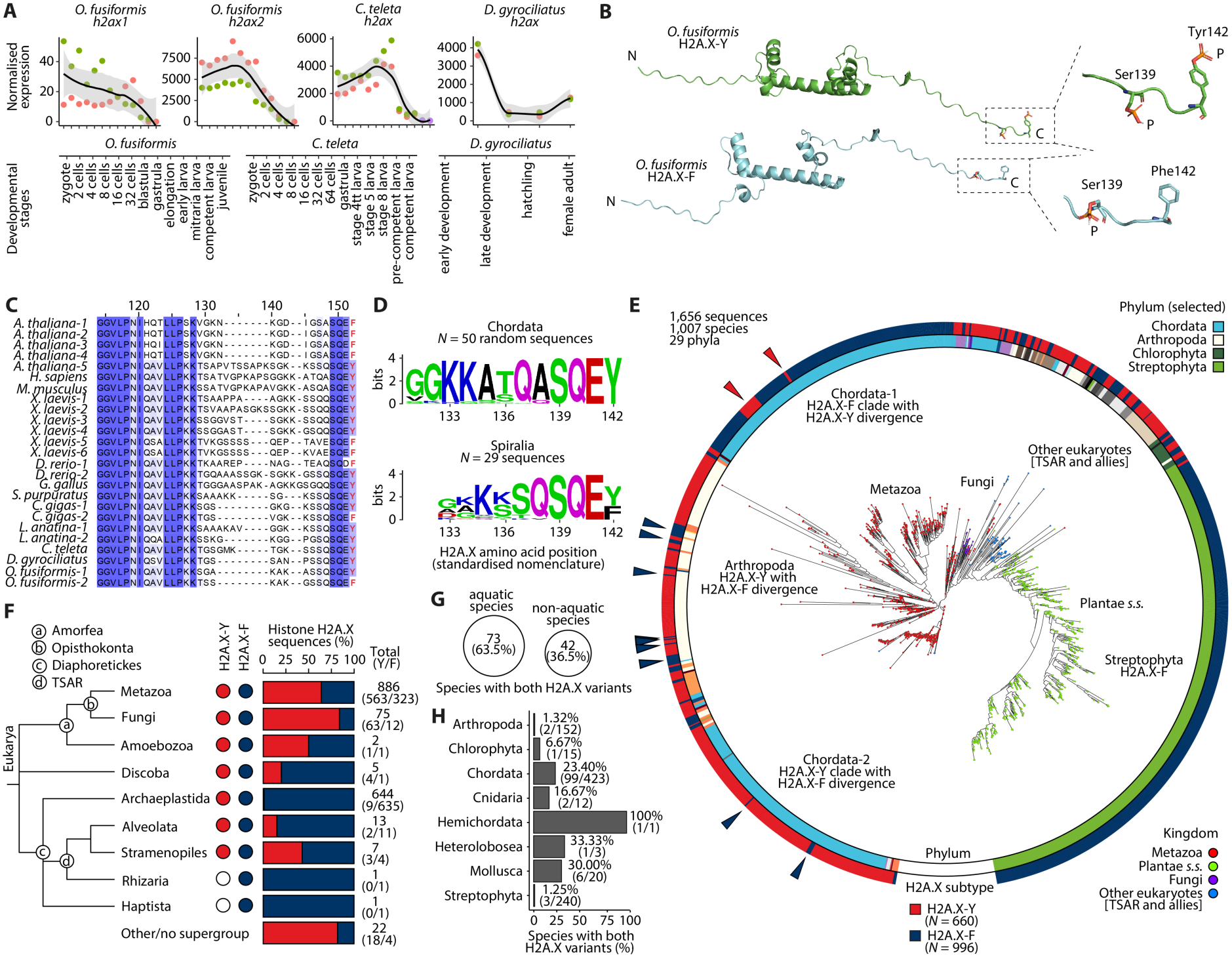
Histone H2A.X variants are a potential source of phenotypic variation in Metazoa. (**A**) Normalised expression levels of *h2ax* variant histone genes for *O. fusiformis* (left, *h2ax1*; and centre left, *h2ax2*), *C. teleta* (centre right), and *D. gyrociliatus* (right). Time points are summarised at the bottom for all three RNA-seq time series. Curves are locally estimated scatterplot smoothing and coloured shaded areas represent the standard error of the mean. (**B**) Renders of AlphaFold3 structural models of the H2A.X-Y and H2A.X-F proteins of *O. fusiformis*. The inset shows a close-up of the C-terminus, where the phosphorylatable Ser139 residues and the Tyr142 and Phe142 residues are depicted as ribbons, highlighting the structural similarities of both amino acids, yet the exclusive phosphorylation of Tyr142. P: phosphate group. (**C**) Multiple sequence alignment (MSA) depicting the C-terminal region of selected H2A.X proteins, highlighting the variation that can be found in the otherwise conserved Y142 residue (position 152 in the MSA, highlighted in red) in key lineages, particularly in *O. fusiformis* (bottom). *A. thaliana*: *Arabidopsis thaliana*; *H. sapiens*: *Homo sapiens*; *M. musculus*: *Mus musculus*; *X. laevis*: *Xenopus laevis*; *D. rerio*: *Danio rerio*; *G. gallus*: *Gallus gallus*; *S. purpuratus*: *Strongylocentrotus purpuratus*; *C. gigas*: *Crassostrea gigas*; *L. anatina*: *Lingula anatina*. The blue gradient represents the sequence identity for each position in the alignment. For a full-length MSA see Supplementary Fig. 10A. (**D**) Sequence logos of the C-terminal region (positions 131–142) of 50 random curated chordate H2A.X sequences (top) and the 29 curated spiralian H2A.X sequences (bottom) obtained from the HistoneDB 2.0 database (Draizen et al. 2016). (**E**) Maximum likelihood evolutionary reconstruction of PHI-BLAST-retrieved H2A.X-Y and H2A.X-F variants. The phylum of sequences is shown in the inner circle, and the H2A.X subtype/variant is shown in the outer circle, both in a colour-coded scale. Coloured arrows point to examples of Y-to-F (blue) and F-to-Y (red) conversions within H2A.X-F or H2A.X-Y clades. *s.s.*: *sensu stricto*; TSAR: Telonemia, Stramenopiles, Alveolata, and Rhizaria. For a fully labelled tree, see Supplementary Fig. 11A. (**F**) Eukaryotic topology as in (Burki et al. 2020) showing the presence/absence, number, and percentage of the different H2A.X variants in the main major eukaryotic lineages with available data. Circled letters denote key eukaryotic supergroups. (**G**) Bubble plot proportional to the number and percentage of aquatic and non-aquatic species in the set of species encoding both a H2A.X-Y and a H2A.X-F variant in their genomes. (**H**) Bar plot showing the number and percentage of species harbouring both H2A.X variants in their genomes in the eight phyla that comprised these species.

To disentangle the evolutionary history of H2A.X, we performed PHI-BLAST searches for H2A.X-Y and H2A.X-F orthologs across Eukarya combined with phylogenetic reconstruction. We recovered 1,656 H2A.X sequences (660 H2A.X-Y and 996 H2A.X-F), mostly belonging to the highly sequenced Metazoa and Plantae clades, across 29 different eukaryotic phyla (Supplementary Fig. 11B–G). Few clear trends could be elucidated beyond almost all plants displaying an H2A.X-F variant, with the few plant H2A.X-Y variants (nine out of 635) being almost entirely restricted to chlorophytes. In arthropods, the ancestral and most common scenario seems to be an H2A.X-Y from which divergent H2A.X-F proteins evolved. Within Chordata, on the other hand, two distinct clades correspond to each of the two H2A.X variants. In each clade, some conversions to the opposite scenario (Y-to-F and F-to-Y transitions) occur, yet generally, there is strong conservation of the amino acid in position 142. This robustly suggests that in chordates these are different genes with distinct evolutionary origins (Fig. 4E; Supplementary Fig. 11A) but that other evolutionary dynamics are also present in animal and other eukaryote lineages (Fig. 4F). Previous research suggested possessing both a H2A.X-Y and a H2A.X-F variant could be an adaptation for quickly developing aquatic species, yet 36.5 % of the species with both variants do not display aquatic lifestyles (Fig. 4G). Here we show, however, that species with this two-H2A.X variant setting are almost exclusively found within Metazoa and are more common than previously thought, as shown in species analysed from the Chordata (23.40 %), Cnidaria (16.67 %), and Mollusca (30 %) (Fig. 4H), likely reflecting H2A.X-Y and H2A.X-F animal variants to be two distinct genes. Altogether, these results demonstrate that histone variants can display intricate phylogenetic histories, highlighting the need for further functional comparative analyses of the genome regulatory implications of the two H2A.X variants in eukaryotes and during animal embryogenesis.

### Histone modifiers show a complex evolution throughout annelid diversification

Besides the histone complement, diversity in the repertoire of histone-modifying enzymes (HME) is the other major potential source of evolutionary variation within histone-based regulation. To address this, we investigated the writers and erasers of histone acetylation and methylation: histone deacetylases (HDAC), histone demethylases (HDM), histone methyltransferases (HMT) of both the lysine-specific (KMT) and arginine-specific (PRMT) types, and histone acetyltransferases (HAT), of both type A and type B. We used a mutual best BLAST hit approach to find candidate sequences in the annelid genomes, which were then subjected to orthology assignment using both maximum likelihood and Bayesian reconstructions in six different phylogenetic analyses (Fig. 5; Supplementary Fig. 12–23). We identified a total of 78 clades corresponding to 77 different genes, of which only three (3.9 %) were not supported by phylogenetic methods, and only five (6.5 %) had bootstrap values or posterior probabilities below 70 or 0.7, respectively, thus making our gene identification highly robust and reliable.

**Figure 5.**
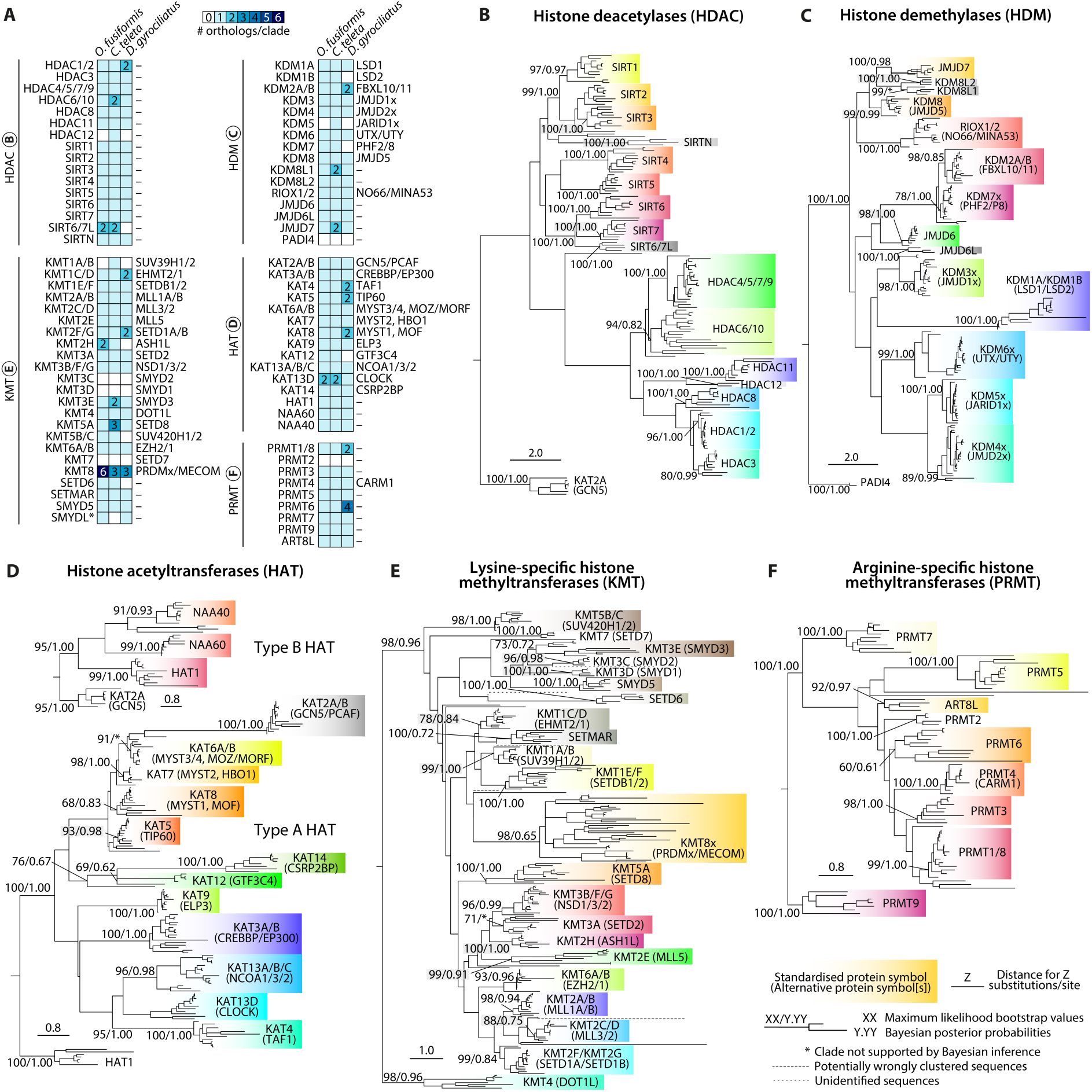
Histone modifiers show a complex evolution throughout annelid diversification. (**A**) Summary heatmaps of the ortholog number in our annelid taxa of the histone modifiers involved in histone methylation and acetylation: HDAC (top left), HDM (top right), HAT (centre right), KMT (bottom left), and PRMT (bottom right). Standardised protein symbols are shown to the right of each summary heatmap, and alternative protein symbols are shown to the left. Some protein symbols are custom for annelid or lineage-specific clades, as described in the text (e.g., SIRT6/7L). Orthologs to more than one gene in mammals are called a single one, separated by strokes (e.g., HDAC1/2). (**B–F**) Summary phylogenies of the gene orthology assignment of HDAC (**B**), HDM (**D**), type A HAT (**D**, bottom), type B HAT (**D**, top), KMT (**E**), and PRMT (**F**) genes, in *O. fusiformis*, *C. teleta*, and *D. gyrociliatus*. Depicted tree topology is based on the maximum likelihood reconstruction. Branch support values represent bootstrap values (0–100 values) and posterior probabilities (0–1 values) at key nodes. Clades supported by maximum likelihood only are flagged with an asterisk (*). Coloured boxes highlight the extent of each histone gene or family. Potentially wrongly clustered sequences and unidentified proteins are shown in long and short-dashed lines, respectively. The scale bar depicts the number of amino acid changes per site along the branches.

### Histone deacetylases

Annelids display at least a *hdac1/2*, *hdac3*, *hdac4/5/7/9*, *hdac8*, and *hdac11* ortholog, as well as one ortholog of each of the known NAD-dependent sirtuins, that is, *sirt1*, *sirt2*, *sirt3*, *sirt4*, *sirt5*, *sirt6*, and *sirt7* (Fig. 5A, B; Supplementary Fig. 12, 13). There is an additional well-assigned sirtuin clade, which we have here termed *sirtuin novel* (*sirtn*), with a single ortholog in all three species, as well as a clade with high similarity to both *sirt6* and *sirt7* genes (*sirt6/7- like*, or *sirt6/7l*), with two copies present in both *O. fusiformis* and *C. teleta*, but not in *D. gyrociliatus*. The latter presents a duplication of the *hdac1/2* gene, whereas the genome of *C. teleta* encodes for two different *hdac6/10* genes. Intriguingly, *C. teleta* also has an additional HDAC gene that clusters independently, and for which the most similar gene is the poorly characterised *hdac12* gene in zebrafish (Fig. 5A, B; Supplementary Fig. 12, 13).

### Histone demethylases

In terms of histone demethylases, annelids encode orthologs of *kdm1a* (*lsd1*), *kdm1b* (*lsd2*), *kdm2a/b* (*fbxl10/11*), *kdm3* (*jmjd1x*), *kdm4* (*jmjd2x*), *kdm5* (*jarid1x*), *kdm6* (*utx/uty*), *kdm7* (*phf2/8*), *kdm8* (*jmjd5*), *riox1/2* (*no66/mina53*), *jmjd6*, and *jmjd7* (Fig. 5A, C; Supplementary Fig. 14, 15). All three species encode a single ortholog of these genes, except for *kdm5*, which is only present in *C. teleta*, the secondary losses of *kdm1b* (*lsd2*) and *kdm7* (*phf2/8*) in *D. gyrociliatus*, and a lineage-specific duplication of *kdm2a/b* (*fbxl10/11*), also in the direct developer *D. gyrociliatus*. We identified a clade with high similarity to *jmjd6*, which we have denoted as *jmjd6-like* (*jmjd6l*), that is nevertheless distinct from *jmjd6* and is present as a single gene in each of the annelids’ genomes. Two other clades phylogenetically closer to *jmjd7* yet automatically assigned as *kdm8* genes were identified. The analysed genomes contain one copy of each, except in *C. teleta*, which harbours two genes within the *kdm8-like 1* (*kdm8l1*) clade. Similarly to all other invertebrates, the protein arginine deiminase *padi4* is absent in our annelids (Fig. 5A, C; Supplementary Fig. 14, 15).

### Histone acetyltransferases

A complete repertoire of nuclear/type A and cytoplasmic/type B occurs in all three annelids (Fig. 5A, D; Supplementary Fig. 16–19). The only exception is *kat12* (*gtf3c4*), which despite being part of a general transcription factor (TFIIIC90) of RNA Pol III (Male et al. 2015; Seifert-Davila et al. 2023), seems to be absent in *D. gyrociliatus*. At least one ortholog was found for all three lineages for *kat2a/b* (*gcn5/pcaf*), *kat3a/b* (*crebbp/ep300*), *kat4* (*taf1*), *kat5* (*tip60*), *kat6a/b* (*myst3/4*, *moz/morf*), *kat7* (*myst2, hbo1*), *kat8* (*myst1*, *mof*), *kat9* (*elp3*), *kat13a/b/c* (*ncoa1/3/2*), *kat13d* (*clock*), *kat14* (*csrp2bp*), *hat1*, *naa60*, and *naa40*. Unexpectedly, the compact genome of *D. gyrociliatus* showed the largest expansions, with *kat4* (*taf1*), *kat5* (*tip60*), and *kat8* (*myst1*, *mof*) duplicated in a lineage-specific manner. But we could also find a likely ancestral duplication common to *O. fusiformis* and *C. teleta* of the *kat13d* (*clock*) gene (Fig. 5A, D; Supplementary Fig. 16–19).

### Lysine-specific histone methyltransferases

The KMT family showed the largest variations between annelids and model organisms. Annelids do not have an ortholog of the chordate-specific *kmt3c* (*smyd2*) and *kmt3d* (*smyd1*) genes (Calpena et al. 2015), or *kmt7* (*setd7*). They do display orthologs for *kmt1c/d* (*ehmt2/1*), *kmt1e/f* (*setdb1/2*), *kmt2a/b* (*mll1a/b*), *kmt2c/d* (*mll3/2*), *kmt2e* (*mll5*), *kmt2f/g* (*setd1a/b*), *kmt2h* (*ash1l*), *kmt3a* (*setd2*), *kmt3b/f/g* (*nsd1/3/2*), *kmt3e* (*smyd3*), *kmt4* (*dot1l*), *kmt5a* (*setd8*), *kmt5b/c* (*suv420h1/2*), *kmt6a/b* (*ezh2/1*), *setd6*, *setmar*, and *smyd5* (Fig. 5A, E; Supplementary Fig. 20, 21). Strikingly, we could only assign a *kmt1a/b* (*suv39h1/2*) ortholog in the case of *C. teleta*. *Dimorphilus gyrociliatus* has suffered losses of *kmt2h* (*ash1l*), *kmt5b/c* (*suv420h1/2*), and *setd6*, but carries two different genes of the *kmt1c/d* (*ehmt2/1*) and *kmt2f/g* (*setd1a/b*) classes. *Owenia fusiformis* has a duplication of the *kmt2h* (*ash1l*) gene, whereas *C. teleta* has one for *kmt3e* (*smyd3*). Moreover, we uncovered an expansion of *kmt5a* (*setd8*) orthologs exclusive of *C. teleta*, which has three different paralogs of this gene. Two sequences of the SET and MYND domain-containing family (SMYD)—one belonging to *O. fusiformis* and another one to *D. gyrociliatus*—did not cluster with any identified KMT clade and were classified broadly as *smyd-like* (*smydl*) genes. Within the *kmt8* (*prdmx/mecom*) clade of genes, both *D. gyrociliatus* and *C. teleta* contained three different genes that we could not subclassify but that may represent different genes. Surprisingly, *O. fusiformis* displayed a large expansion in this family, with up to six different genes that could not be resolved further (Fig. 5A, E; Supplementary Fig. 20, 21).

### Arginine-specific histone methyltransferases

Lastly, we also studied the complement of PRMT enzymes (Fig. 5A, F; Supplementary Fig. 22, 23). Excluding *prmt2*, which could not be found in any polychaete, all others are present, namely, *prmt1/8*, *prmt3*, *prmt4*, *prmt5*, *prmt6*, *prmt7*, and *prmt9*. A further PRMT gene was identified in all three species that only clustered with the fruit fly *Art8* gene and that we deemed *art8-like* (*art8l*). The expansions of *prmt1/8* and *prmt6* in the genome of *D. gyrociliatus* are thus the only changes within the PRMT family in the studied annelids.

### PRMT6 expansions led to domain fusions and likely catalytically dead enzymes

Considering that *D. gyrociliatus* is a case of extreme genome compaction, frequently associated with gene losses, it is striking that it appeared to possess four different *prmt6* genes (hereon referred to as *prmt6-a* through *prmt6-d*) that subcluster independently of the rest of the sequences of the other species within the *prmt6* clade (Fig. 5A, F; Supplementary Fig. 22, 23). PRMT6 enzymes catalyse type II arginine methylation reactions and yield monomethyl arginine (Rme1/MMA) and asymmetric dimethyl arginine (Rme2a/ADMA) (Fig. 6A) in both histones and non-histone proteins and have been shown to regulate cell proliferation and senescence (Blanc and Richard 2017; Lorton and Shechter 2019). Both the PRMT6-A and PRMT6-C proteins are significantly longer—with a full length of 871 and 1090 amino acids long, respectively—than what is expected of a PRMT6 ortholog (340–380 residues) (Fig. 6B). We analysed their domain and region composition and predicted their protein structure (Fig. 6B–D, Supplementary Fig. 24, 25) and uncovered that while *O. fusiformis* and *C. teleta* have a very conserved protein structure and domain length, this is only true for *prmt6-b* in *D. gyrociliatus*. Meanwhile, both *prmt6-c* and *prmt6-d* have a shorter *S-*adenosyl methionine (SAM)-dependent methyltransferase class I domain, and *prmt6-a* and *prmt6-c* contain additional regions and domains in the N-terminal region of the protein, namely three consecutive galactose/rhamnose-binding lectin domains and a progesterone-induced blocking factor 1 family region, respectively (Fig. 6B, D, Supplementary Fig. 25A). Transcriptomic validation of the fusion genes showed that the one present in *prmt6-c* is, however, likely a false positive and a result of a wrongful annotation. Unlike in *prmt6-a*, where both presupposed fused parts of the gene show continuous transcription at similar levels, the *prm6-c* gene showed large discrepancies in read density across the model and displayed a large number of unexplained antisense reads (Supplementary Fig. 25B, C). No PRMT6 ortholog contains the functionally critical tyrosine dyad (Y47 and Y51) in the PRMT characteristic motif (Fig. 6C, Supplementary Fig. 24). There are also a very high number of amino acid changes in highly conserved positions known to interact with either the SAM cofactor (e.g., R66, E141, and S169) and/or with the arginine residue (e.g., E164 and E317) (Fig. 6C, Supplementary Fig. 24). In fact, structural changes affecting the characteristic PRMT domain are evident, most notably in the lack of tertiary structures of the alpha helices where the PRMT characteristic motif and the tyrosine dyad are located (Fig. 6D, Supplementary Fig. 25A). These sequence and structure alterations are likely disrupting the potentially PRMT6 enzymatic function, thus perhaps relaxing selection pressures and allowing for various gene expansions and divergence events to occur. Altogether, our data indicate that in an annelid with a highly compact genome and a reduced and simplified histone complement, like *D. gyrociliatus*, the plasticity in histone-based regulation required during embryogenesis most likely resides in the conservation and, in some cases, diversification of the HME repertoire.

**Figure 6.**
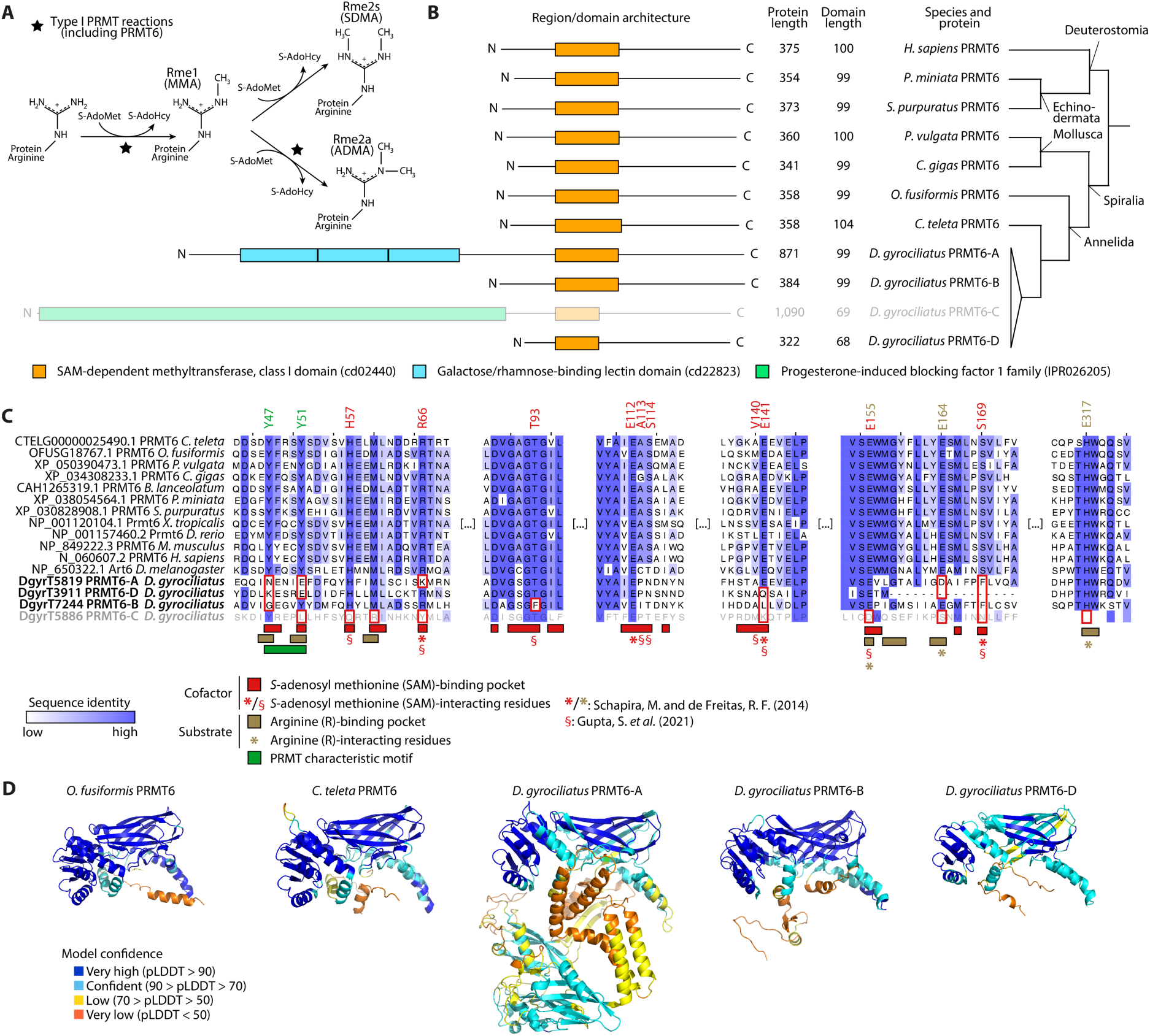
PRMT6 expansions in *D. gyrociliatus* led to domain fusions and likely catalytically dead enzymes. (**A**) Arginine methylation reactions catalysed by PRMT proteins. Reactions catalysed by type I PRMT proteins like PRMT6 are marked with a star. S-AdoMet: *S*-adenosyl methionine/SAM; S-AdoHcy: *S*-adenosyl homocysteine/SAH; Rme1/MMA: monomethyl arginine; Rme2s/SDMA: symmetric dimethyl arginine; Rme2a/ADMA: asymmetric dimethyl arginine. (**B**) Domain/region architecture of selected PRMT6 proteins. PRMT6-B and PRMT6-D in *D. gyrociliatus* harbour reduced SAM-dependent methyltransferase, class I domains. PRMT6-A and PRMT6-C in *D. gyrociliatus* contain additional domains/regions in the N-terminal region not expected for a PRMT6 ortholog. PRMT6-C may be an annotation artefact and is therefore greyed out. Phylogenetic relationships of the species displayed here are shown to the right of the schematics. *P. miniata*: *Patiria miniata*; *P. vulgata*: *Patella vulgata*. (**C**) MSA of representative PRMT6 sequences, trimmed to the substrate and cofactor binding regions. At the bottom of the MSA, all four putative PRMT6 orthologs from *D. gyrociliatus* are highlighted in bold. PRMT6-C is greyed out here as well to show its likelihood as an annotation artefact. Key protein regions and residues are highlighted under the MSA. Amino acids with a specified position (as per the human PRMT6 nomenclature) are SAM and arginine-interacting residues. Residues inside red boxes denote conserved positions in key regions or key interacting residues with no conservation in one or more of the orthologs of *D. gyrociliatus*. *X. tropicalis*: *Xenopus tropicalis*. *: SAM-interacting and arginine-interacting residues determined via homology to PRMT4 (CARM1), as in (Schapira and Freitas 2014); §: SAM-interacting residues determined directly in PRMT6, as in (Gupta et al. 2021). For the full-length MSA trimmed to the R43– Y359 positions (as per the human PRMT6 nomenclature), see Supplementary Fig. 24. (**D**) Renders of the AlphaFold3 structural predictions of the annelid PRMT6 orthologs whose gene models were validated using transcriptomics data. From left to right: *O. fusiformis* PRMT6, *C. teleta* PRMT6, and *D. gyrociliatus* PRMT6-A, PRMT6-B, and PRMT6-D. Renders are roughly aligned to the SAM-dependent methyltransferase, class I domain (cd02440). Render colour depicts the model’s confidence in the prediction.

### Life cycle correlates with heterochronies of histone modifiers

Histone-modifying enzymes (HMEs) constitute a highly plastic source of regulative variation that underpins changes in gene expression patterns (Bannister and Kouzarides 2011; Zhou et al. 2011; Butler et al. 2012; Zhao and Shilatifard 2019; Morgan and Shilatifard 2020). To investigate how temporal shifts in the deployment of orthologous developmental programmes could correlate with large changes in HME expression, we leveraged the entire developmental RNA-seq time series, from the zygote to competent larva/adult stages, for all three annelid species. All expressed transcripts were clustered using an unbiased soft *k*-means approach into an optimal number of 15 clusters (*O. fusiformis* and *C. teleta*) or eight clusters (*D. gyrociliatus*) (see Methods, Supplementary Fig. 26A–C). Next, we assigned the clusters to their corresponding developmental stages according to their maxima and classified them as cleavage or post-cleavage clusters (*O. fusiformis* and *C. teleta*) or early development and late development clusters (*D. gyrociliatus*). We further subdivided post-cleavage clusters in the indirect developers into pre-larval and post-larval clusters (Supplementary Fig. 26D–F). We profiled the expression dynamics of all five superfamilies of histone modifiers described above across the development of the three species. Given the broad array of biological readouts different modifiers from the same superfamily can have, as expected no clear superfamily-specific trends were common to all three taxa (Supplementary Fig. 27, 28). We then decided to focus on gene-wise expression dynamics. To do this, we compared the timing of deployment of every histone modifier with single-copy orthology either between both indirect-developing species or common to all three species (Fig. 7; Supplementary Fig. 29, 30). When comparing *O. fusiformis* and *C. teleta*, we unravelled how there are more histone modifiers under heterochronic shifts between cleavage and post-cleavage clusters (29 genes, 44.6 % of the total) than sharing the same expression pattern (27 genes, 41.5%). This is mostly due to 21 enzymes (32.3 %) being expressed early during cleavage stages in *O. fusiformis* but late in post-cleavage time points in *C. teleta*; yet the opposite is also true for eight of the enzymes (12.3 %) (Fig. 7A, top). Heterochronic genes belong to all five superfamilies of modifiers. In the first group of genes (group I), or those delayed in the lecithotrophic larva, we can find, among others, the H3K27ac acetyltransferase *kat3a/b* (*crebbp/ep300*) and one of the circadian clock gene orthologs *kat13d-a* (*clock-a*); four different sirtuin genes, namely *sirt3*, *sirt4*, *sirt5*, and *sirt6*; both lysine-specific demethylases *kdm1a* (*lsd1*) and *kdm1b* (*lsd2*); the H3K79-specific *kmt4* (*dot1l*) and H3K27-specific *kat6a/b* (*ezh2/1*); as well as *prmt4* and *prmt5*. Displaced towards a late expression in the planktotrophic larva (group II), examples include the ortholog to the steroid receptor co-activators *kat13a/b/c* (*ncoa1/3/2*)*, hdac11* and *sirt1*, the bifunctional histone demethylase/ribosome hydroxylase *riox1/2* (*no66/mina53*), the inactive *kmt2e* (*mll5*), and the previously discussed *prmt6* gene (Fig. 7A, top, D; Supplementary Fig. 29). Contrary to our null expectation of finding heterochronic shifts between pre-larval and post-larval expression, all genes with a post-cleavage expression were consistently deployed at the same life cycle time point—either before or after larval formation—between *C. teleta* and *O. fusiformis* (Fig. 7A, bottom), suggesting that histone-based regulatory mechanisms in later developmental stages might be broadly conserved between indirect developing annelids.

**Figure 7.**
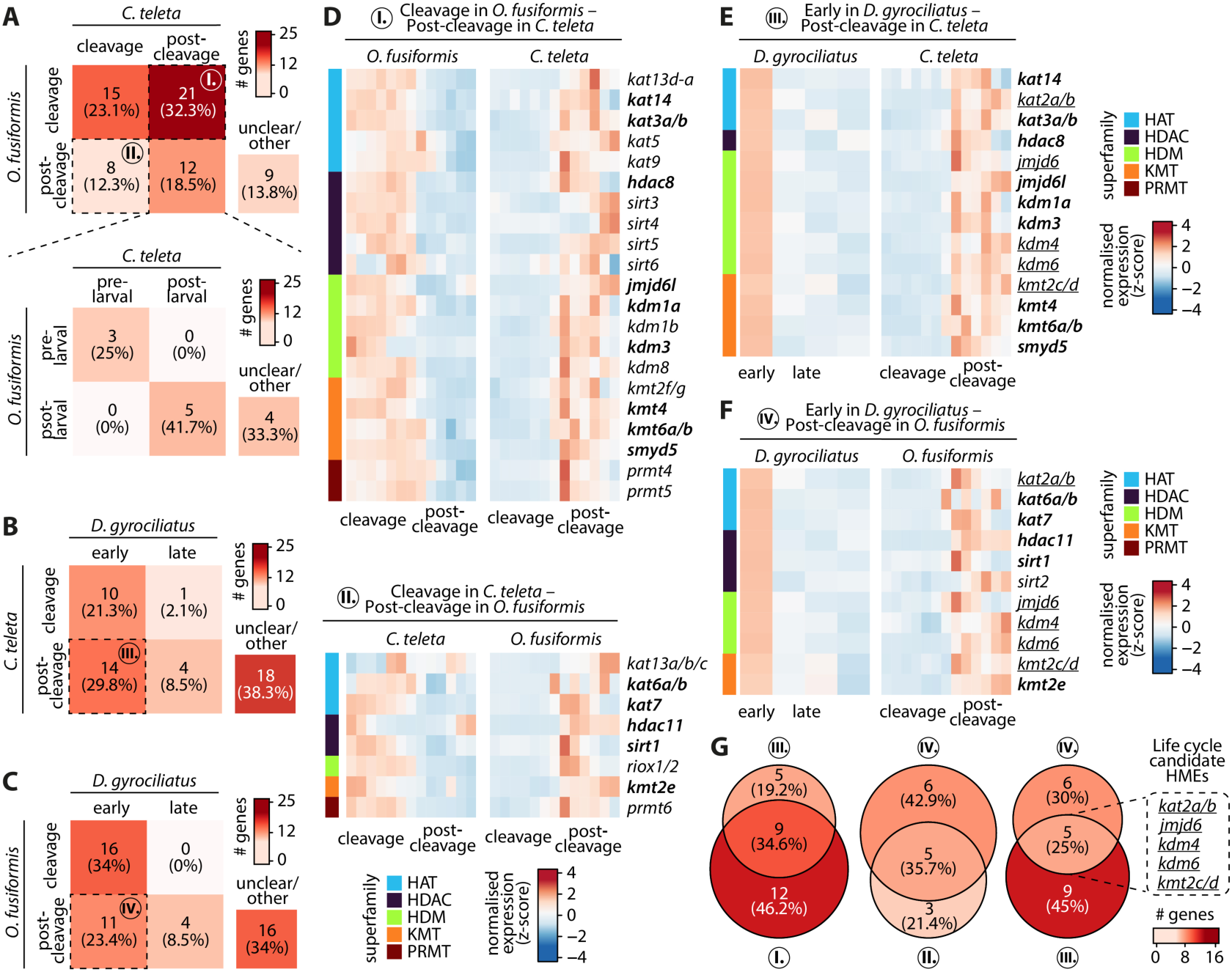
Life cycle correlates with heterochronies of histone modifiers. (**A–C**) Pair-wise comparative gene expression analyses of HMEs across species by classifying clusters of temporally co-regulated genes into early/pre-cleavage and late/post-cleavage clusters and pre- and post-larval clusters. The colour scale denotes the number of genes. Species compared are *O. fusiformis* and *C. teleta* (**A**), both using the cleavage/post-cleavage classification (**A**, top) and the pre- and post-larval one (**A**, bottom); *C. teleta* and *D. gyrociliatus* (**B**); and *O. fusiformis* and *D. gyrociliatus* (**C**). Unclear/other includes comparisons of genes clustered in an unclassified transitional cluster in at least one of the two species (see Supplementary Fig. 26). Dotted lines highlight the gene sets (I–IV) of genes under heterochronic shifts. (**D–F**) Heatmaps of normalised expression dynamics of the gene sets of genes under heterochronic shifts. Colour scale denotes normalised gene expression in a z-score scale. Species compared are *O. fusiformis* (left) and *C. teleta* (right) (**D**, top, gene set I); *C. teleta* (left) and *O. fusiformis* (right) (**D**, bottom, gene set II); *D. gyrociliatus* (left) and *C. teleta* (right) (**E**, gene set III); and *D. gyrociliatus* (left) and *O. fusiformis* (right) (**F**, gene set IV). (**G**) Venn diagrams of the intersections of heterochronic gene sets. The colour scale denotes the number of genes. Intersections between gene sets I and III (left) and gene sets II and IV (centre) represent species-specific heterochronies related to larval development in *O. fusiformis* and *C. teleta*, respectively, which correspond to gene symbols in bold in **D–F**. Intersection between gene sets III and IV (right) represent consistent heterochronies related to the life cycle, corresponding to gene underlined gene symbols in **E** and **F**.

We thus focused on the comparison between direct and indirect developers, i.e., between *D. gyrociliatus* and either *C. teleta* or *O. fusiformis*. As observed with transcription factors (Martín-Zamora et al. 2023b), a significant number of HMEs are pre-displaced from late/post-cleavage expression in *C. teleta* (group III, 14 genes, 29.8 %) and *O. fusiformis* (group IV, 11 genes, 23.4 %) to early development in *D. gyrociliatus* (Fig. 7B, C). Genes involved in these shifts are varied, with all superfamilies except the PRMT enzymes being represented in both groups of heterochronic genes (Fig. 7E, F; Supplementary Fig. 30). A critical advantage of these pair-wise comparisons is that we can intersect them and find those changes that are consistent and specific to direct and indirect developers (Fig. 7G). By comparing groups I. and III., those where genes are shifted from early/cleavage expression in either *O. fusiformis* or *D. gyrociliatus* to late/post-cleavage expression in *C. teleta*, we can robustly describe modifiers specific to the development of the lecithotrophic larva of *C. teleta* (Fig. 7D, top, E, G, left). These are *kat3a/b* (*crebbp/ep300*), *kat14* (*csrp2bp*), *hdac8*, *jmjd6l*, *kdm1a* (*lsd1*), *kdm3* (*jmjd1x*), *kmt4* (*dot1l*), *kmt6a/b* (*ezh2/1*), and *smyd5*. The equivalent can be produced for *O. fusiformis* by comparing groups II and IV, which yields the modifiers specific to the development of the planktotrophic larva of *O. fusiformis* (Fig. 7D, bottom, F, G, centre), namely *kat6a/b* (*myst3/4*, *moz/morf*), *kat7* (*myst2*, *hbo1*), *hdac11*, *sirt1*, and *kmt2e* (*mll5*). Lastly, we compared groups III and IV, i.e., those genes that are consistently pre-displaced to early expression in *D. gyrociliatus* from late/post-cleavage expression in both indirect developers (Fig. 7E, F, G, right), which included *kat2a/b* (*gcn5/pcaf*), three different demethylases, *jmjd6*, *kdm4* (*jmjd2x*), and *kdm6* (*utx/uty*), and *kmt2c/d* (*ehmt2/1*). Therefore, a few orthologous HMEs exhibit different temporal expression dynamics between the studied annelids. These might represent key epigenetic regulators of specific developmental programmes that differ between direct and indirect annelids, thus emerging as a tractable gene set for functional manipulations in the future.

### Adult annelids harbour distinct histone modification enrichments

Finally, given the differences in the HME repertoires between species and their expression timing across development, we investigated whether adults may consequently display different levels of key histone modifications. To assess this, we used quantitative mass spectrometry-based detection of hPTMs in adult specimens of *O. fusiformis* and *C. teleta* and focused on the methylation and acetylation of residues in H3 and H4 histones (Fig. 8A, B; Supplementary Fig. 31A–C). The three biological replicates with two technical replicates strongly clustered by species (Fig. 8C and Supplementary Fig. 31D). Significant differences in key hPTMs were immediately obvious in several peptides. Most notable were the large differences in H3K56 methylation and H3K79 methylation. H3K56 is mostly unmethylated (H3K56me0) or di- and trimethylated (H3K56me2 and H3K56me3) in *C. teleta* but more abundant in its monomethylated form (H3K56me1) in *O. fusiformis* (Fig. 8D, left). Notably, these differences are consistent with the differences in the presence/absence of HMEs between species and their developmental heterochronies. H3K56me3 is known to be introduced in humans by the KMT1A (SUV39H1) and KMT1B (SUV39H2) enzymes. The almost exclusive presence of H3K56me3 in *C. teleta* matches our predictions, given that the annelid KMT1A/B (SUV39H1/2) ortholog is present in *C. teleta* but absent in *O. fusiformis*. On the contrary, the ortholog to the human H3K56me1-depositing enzyme KMT1D (EHMT2), which is KMT1C/D (EHMT2/1) in annelids, is present in *O. fusiformis*, likely explaining the accumulation of H3K56me1 in this species. These different hPTM abundances may have deep biological consequences, as it has been described that while H3K56me1 is involved in regulating DNA replication (Yu et al. 2012), H3K56me3 is a heterochromatic mark that largely overlaps with the constitutive heterochromatin mark H3K9me3 (Jack et al. 2013; Colmenares et al. 2017). We also found large differences in unmodified and monomethylated H3K79, with the former being the dominant form in *C. teleta* and the latter in *O. fusiformis* (Fig. 8D, right). Some well-studied and conserved methylation hPTMs like H3K4me2 and H3K4me3—among many others—appeared to be more enriched in one species from EpiProfile analyses, though statistical quantifications showed no significant differences between species (Fig. 8; Supplementary Fig. 31E). Other methyl marks like H3K36me1/2/3 and H3K27me2/3 were significantly more abundant in one of the annelid lineages. We also found this to be true for two acetylations, namely H3K23ac and H3K27ac, both significantly more abundant in *C. teleta* (Fig. 8F; Supplementary Fig. 31F–H), which could be the result of the heterochronic shift between early/cleavage expression in *O. fusiformis* and late/post-cleavage expression in *C. teleta* of five different HAT genes: *kat5* (*tip60*), *kat9* (*elp3*), *kat13d-a* (*clock-a*), *kat14* (*csrp2bp*), but most interestingly, *kat3a/b* (*crebbp/ep300*). H3K9ac, however, escapes this pattern and appears more abundant in *O. fusiformis* (Supplementary Fig. 31F). Likewise, this may be associated with chromatin and transcription differences in the annelids, as histone acetylations like H3K23ac and H3K27ac are known to be tightly associated with increased transcription across species (Creyghton et al. 2010; Martin et al. 2021; Zhang et al. 2023). Altogether, our data describe a core repertoire of annelid histone methylation and acetylation of histones H3 and H4, revealing some key differences in the adult levels in the abundance of key hPTM, likely to be the result of differences in the repertoire and expression dynamics of HMEs.

**Figure 8.**
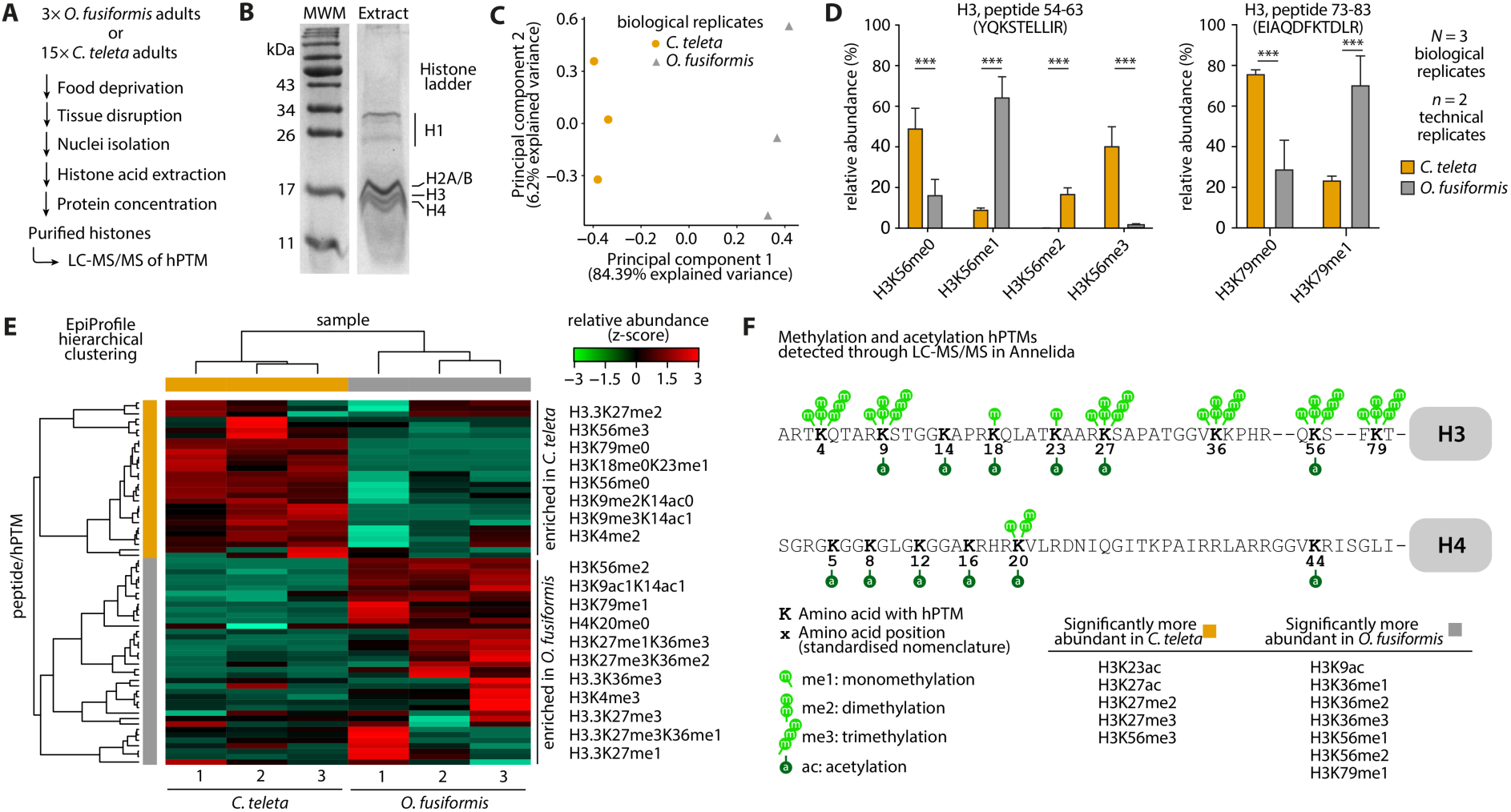
Adult annelids harbour distinct histone modification enrichments. (**A**) Histone acid extraction protocol schematic. (**B**) Sample SDS-PAGE analysis of acid-extracted histones from 15 adult specimens of *C. teleta*. See uncropped gel in Supplementary Fig. 31B. (**C**) Principal component analysis of the histone H3 and H4 hPTM profiles derived from LC-MS/MS experiments, averaged by biological replicate, for all analysed samples of *O. fusiformis* and *C. teleta*. (**D**) Relative abundance bar plots of the H3 54–63 peptide (YQKSTELLIR) based on H3K56 methylation status (left), and the H3 73–83 peptide (EIAQDFKTDLR) based on H3K79 methylation status (right), in *O. fusiformis* and *C. teleta*. Error bars represent standard deviation. *P* values were derived from two-tailed Student’s *t*-tests. ***: *P* value < 0.001. (**E**) Hierarchically clustered heatmap of the relative abundance of all detected histone H3 and H4 peptide/hPTMs. The numbers below samples are for biological replicates. In orange, peptides/hPTMs enriched in *C. teleta* samples; in grey, those enriched in *O. fusiformis*. The list of peptides/hPTMs to the right of the heatmap only includes representative examples. (**F**) Annelid acetylation and methylation hPTM landscape in histones H3 and H4. hPTMs that are statistically significantly more abundant in either species are listed here, based on quantifications shown in **D** and Supplementary Fig. 31E–H.

## Conclusions

In this study, we profiled the histone, HME, and hPTM repertoires to trace the evolution and developmental dynamics of histone-based regulation in three annelid lineages with contrasting genomic features and ontogenetic trajectories (Fig. 1B). Through a multi-omics approach combining comparative genomics, public bulk RNA-seq and ATAC-seq datasets, and the generation of mass spectrometry proteomic data, we identified multiple layers of histone-based regulation common and specific to these annelids that, in some cases, correlate with their life cycles and developmental modes, raising testable and tractable hypotheses to further explore how this genome regulatory mechanism underpins animal embryonic and life cycle evolution.

The in-depth characterisation of histones, histone modifiers, and histone modifications revealed largely stable and conserved repertoires in our three focal annelids, vastly differing from the large differences we recently described in the DNA methylation levels and associated machinery for these same species (Guynes et al. 2024). The genomic organisation of histone genes of *O. fusiformis* and *D. gyrociliatus* determined in this study revealed that H3 and H4 genes are contiguous and always transcribed from the same strand. Yet, when compared with other annelids and molluscs (del Gaudio et al. 1998; Eirín-López et al. 2002; Albig et al. 2003), at least two or more chromosomal rearrangements are required to explain the genomic organization differences, suggesting that either Mollusca or Annelida (or both) have experienced multiple changes in the histone loci from their last common ancestor. Remarkably, *D. gyrociliatus* displays the lowest characterised histone copy number in a free-living animal but a largely complete (and expanded in some instances) HME repertoire. While we cannot exclude the latter to be explained by the multitude of non-histone targets from HMEs, paradoxically, though, this setting is opposite to *Oikopleura dioica*, another animal lineage with an extremely compact genome that has, however, experienced rapid innovation in the histone repertoire (Moosmann et al. 2011). Therefore, genome reduction impacts histone evolution in different ways, putting *D. gyrociliatus* forth as an ideal animal to study the functional impact of hPTMs in animals, as it would simplify the more complex functional investigations implemented in other organisms with more redundant histone repertoires, such as *D. melanogaster* (Günesdogan et al. 2010; McKay et al. 2015) and, most recently, mice (Sankar et al. 2022).

While previous work recapitulating molecular heterochrony invoked several epigenetic regulators, most notably microRNAs (miRNAs) (Slack and Ruvkun 1997; Moss 2007; Zou et al. 2013; Ouchi et al. 2014; Armakola and Ruvkun 2019), less is known about the role of the histone-based regulation machinery in this fundamental developmental and evolutionary process. Through comparative gene expression analyses of HMEs, we discovered that the expression dynamics of HMEs are not always conserved between annelids. Instead, many of them—in some cases more than 50 % of the studied genes—are pre- or post-displaced between larval phenotypes and between species possessing or lacking a larval stage. Very interestingly, HDMs were the most popular category of HMEs consistently shifted between late expression in both indirect developers and early expression in *D. gyrociliatus*, the same heterochrony type previously described for the trunk patterning programme (Martín-Zamora et al. 2023b). These included *kdm4* (*jmjd2x*), *kdm6* (*utx/uty*) and *jmjd6*, whose known roles in mammals (Greenfield et al. 1998; Whetstine et al. 2006; Chang et al. 2007; Shpargel et al. 2012; Liu et al. 2013) might indicate a potential removal of methylation marks of both permissive (H3K36me3) and repressive nature (H3K27me3 and H3K9me3) earlier in direct development to accelerate the expression and subduing of pre-displaced transcriptional programmes. Altogether, our work sets the foundations for the study of histone-based regulation in annelids and other non-model systems, defining a set of tractable HMEs and hPTMs, whose detailed study, from spatial expression and regulatory dynamics to, most importantly, loss-of-function studies, will help understand how variation in chromatin regulation contribute to the fascinating phenotypic evolution found in Spiralia and animals generally.

## Materials and Methods

### Adult culture

Sexually mature *Owenia fusiformis* Delle Chiaje, 1844 adults were obtained from subtidal waters near the Station Biologique de Roscoff (Centre National de la Recherche Scientifique (CNRS)-Sorbonne University, Roscoff, France) and cultured in-house as described before (Carrillo-Baltodano et al. 2021). *Capitella teleta* Blake, Grassle & Eckelbarger, 2009 and *Dimorphilus gyrociliatus* (O. Schmidt, 1857) were cultured following previously described protocols (Seaver et al. 2005; Kerbl et al. 2016b).

### Histone genes mining

Human core histone protein sequences, i.e., H2A, H2B, H3, and H4 proteins, were retrieved from NCBI following the gene definitions provided in the HUGO Gene Nomenclature Committee (HGNC). Curated histone variants sequences from model organisms corresponding to the H2A.X, H2A.Z/H2Av, H3.3, cenH3/CENP-A, and macroH2A subfamilies were downloaded from the NCBI Histone Variants Database 2.0 (HistoneDB 2.0) (https://www.ncbi.nlm.nih.gov/research/histonedb) (Draizen et al. 2016) (Supplementary Table 1). The protein2genome model of the exonerate tool (EMBL-EBI) was employed to align all fetched sequences to either the unmasked or softmasked scaffold-level genome assemblies of *O. fusiformis* (Martín-Zamora et al. 2023b), *C. teleta* (Simakov et al. 2013), and *D. gyrociliatus* (Martín-Durán et al. 2021). The resulting gff files containing putative orthologs were formatted to gff3 format using custom code. The AGAT v.0.6.0. suite of scripts (Dainat et al. 2021) and the gffutils v.0.8.4 package (https://github.com/daler/gffutils) were used to merge transcripts with overlapping codifying sequences, remove identical transcripts, and keep the longest isoform of gene models containing monoexonic transcripts only. Resulting putative sequences were aligned by histone family using MAFFT v.7. (Katoh et al. 2019) with the G-INS-i iterative refinement method, BLOSUM62 as a scoring matrix and a 1.53 gap penalty. Multiple sequence alignments (MSAs) were used as support, together with visual inspections of the corresponding genomic loci in IGV v.2.8.13. (Robinson et al. 2011), to manually curate histone sequences. Transcripts around or in poorly sequenced regions of the genomes and those with non-canonical splicing sites were discarded. Sequences with frameshifts or in-frame termination codons were also discarded. The sequence of multiexonic transcripts was inferred from full-length RNA-seq transcripts from previously published data for all three species (Burns and Pechenik 2017; Martín-Durán et al. 2021; Martín-Zamora et al. 2023b) found through a tblastn search (Camacho et al. 2009) of partial alignments. Only a single transcript per gene and locus was kept, as alternative splicing leading to different protein sequences is rare in histone genes (Bönisch et al. 2012; Martire and Banaszynski 2020) and could not be inferred with certainty from a genome alignment-based approach. Gene models were then lifted off from the scaffold-level assembly of *O. fusiformis* (GenBank: GCA_903813345.1) to the updated chromosome-level assembly (GenBank: GCA_903813345.2) using Liftoff v.1.6.1 (Shumate and Salzberg 2021). Curated protein sequences were then extracted using gffread v.0.12.1 (Pertea and Pertea 2020). Alignments and the corresponding alignment files in .fasta and .nexus formats, updated filtered gene models, transcript and protein files, and genome annotation reports for all three annelid taxa are available in the GitHub repository (see Data Availability section).

### Histone genes orthology assignment

Curated core histone and histone variant sequences from model organisms were retrieved from the NCBI (Supplementary Table 2) and aligned with the curated annelid histones. Sequences were aligned using MAFFT v.7. (Katoh et al. 2019) as described above. To validate a preliminary sequence-based orthology assignment, a phylogenetic analysis was performed on the resulting MSA with a rtREV amino acid substitution matrix (Dimmic et al. 2002) to account for transition rates. We allowed for a proportion of invariable sites (+I) and used the discrete gamma model (Yang 1994) with four rate categories (G4) to describe site evolution rates, together with an optimisation of amino acid frequencies using maximum likelihood (ML) in IQ-TREE v.2.0.3 (Nguyen et al. 2015). 1,000 ultrafast bootstraps (BS) (Hoang et al. 2018) were used to extract branch support values, but bootstrapping convergence was reached earlier. Further support in the form of posterior probabilities was obtained from Bayesian reconstruction in MrBayes v.3.2.7a (Ronquist and Huelsenbeck 2003), using the same rtREV+F+I+G4 model. Two runs with four chains of Markov chain Monte Carlo (MCMC) analyses were run for 50,000,000 generations. Tree visualisation and editing were done using FigTree v.1.4.4 (http://tree.bio.ed.ac.uk/software/figtree/). The resulting histone gene orthology assignment is summarised in Supplementary Table 3 and described in detail for each species in Supplementary Tables 4–6. Alignments and the corresponding alignment files in .fasta and .nexus formats are available in the GitHub repository (see Data Availability section).

### Histone gene structure analysis

Histone gene structure was analysed using the GenomicFeatures v.1.46.5 and GenomicRanges v.1.46.1 packages (Lawrence et al. 2013). Gene length, aggregated intron length, total and average intron length/gene, and total and average intron number/gene were analysed. For gene structure comparisons between the three species of the histone variants with known role and single-copy orthology, i.e., H2A.Z, H2A.X, macroH2A, cenH3, and H3.3, repeated measures one-way ANOVAs were performed, followed by two-tailed paired Student’s *t*-test, when applicable. For *O. fusiformis*, we considered the *h2ax2* gene (H2A.X-F variant) the H2A.X ortholog, as it is the paralog with the highest expression levels in every developmental stage. For all other comparisons, we employed one-way ANOVAs, followed by two-tailed post-hoc Tukey tests for pair-wise comparisons, when applicable. All *P* values derived from pair-wise comparisons were adjusted using the stringent Bonferroni method for multiple testing correction (Supplementary Table 7).

### Chromatin accessibility profiling

Chromatin accessibility dynamics were analysed using publicly available coverage .bw files corresponding to ATAC-seq experiments for all three species (Martín-Durán et al. 2021; Martín-Zamora et al. 2023b) (Supplementary Table 8). Metagene enrichment analysis was computed using computeMatrix and plotHeatmap commands in deepTools v.3.4.3 (Ramírez et al. 2016). Visualization of peak tracks and gene structures was conducted using pyGenomeTracks v.2.1 (Lopez-Delisle et al. 2020) and deepTools v.3.4.3 (Ramírez et al. 2016).

### Gene expression profiling

Gene expression dynamics were re-profiled using publicly available developmental transcriptomes for all three species (Burns and Pechenik 2017; Martín-Durán et al. 2021; Martín-Zamora et al. 2023b) (Supplementary Table 9) to include the previously incomplete histone gene models, from zygote to juvenile stages for *O. fusiformis*, from zygote to competent larva stages for *C. teleta*, and from early embryogenesis to female adult for *D. gyrociliatus*, as described before. Trimmomatic v.0.3980 (Bolger et al. 2014) was used to remove sequencing adaptors. Clean reads were pseudo-aligned to the updated filtered gene models using kallisto v.0.46.2114 (Bray et al. 2016), and DESeq2 v.1.30.1 (Love et al. 2014) was used to normalise counts between samples within each species (Supplementary Tables 10–15). Unidentified variant genes with DESeq2 normalised expression values below ten for all developmental time points and biological replicates were flagged as putative pseudogenes (Supplementary Tables 4–6). Gene expression matrices in TPM and DESeq2 normalised values for all three species are available in the GitHub repository (see Data availability section).

### Amplification of H2A.X variants in O. fusiformis

To validate the RNA-seq data and further confirm the expression of the *h2ax1* (*h2ax-y*) and *h2ax2* (*h2ax-f*) genes, we used four different combinations of two forward (Fw) and two reverse (Rv) primers (Supplementary Table 16) per gene to selectively amplify the transcript from a cDNA pool obtained from a mixture of developmental stages from *O. fusiformis* ranging from the zygote to the juvenile adult. For each gene, 1 µl cDNA was mixed with 1 µl of 10 µM Fw primer (1 or 2), 1 µl of 10 µM Rv primer (1 or 2), 0.5 µl of 10 mM dNTPs mix, 2.5 µl of 10× ThermoPol reaction buffer, and 0.15 µl of 5,000 U·ml^-1^ Taq DNA polymerase, and molecular biology-grade water for a final reaction volume of 25 µl. Samples were amplified by PCR in a thermocycler with a heated lid under the following conditions: initial denaturation at 95 °C for 2 min; 30 cycles of 95 °C for 15 s, 57 °C for 15 s, and 68 °C for 1 min; final extension at 68 °C for 5 min and hold at 12 °C. Length of resulting amplicons (Supplementary Table 17) was analysed via agarose electrophoresis.

### Evolutionary analysis of histone H2A.X variants

An MSA with representative H2A.X sequences across Metazoa (Supplementary Table 18) was obtained using MAFFT v.7. (Katoh et al. 2019) as described above. To mine histone H2A.X variants across Eukarya, we performed a protein PHI-BLAST (Zhang et al. 1998) against the non-redundant protein sequences (nr) public database using the H2A.X-Y (search 1) and the H2A.X-F (search 2) orthologs from *O. fusiformis*, using SQ[DE]F and SQ[DE]Y as PHI patterns, for search 1, and search 2, respectively, and a maximum of 1,000 target sequences. This ensured the presence of the SQ[DE][FY] carboxyterminal (C-terminal) motif of H2A.X and simplified downstream analysis. Non-eukaryotic sequences were discarded (four for search 1, one for search 2) (Supplementary Tables 19, 20). Taxonomical information for sequences was obtained using the ETE 3 library (Huerta-Cepas et al. 2016: 3), and the eukaryotic supergroup was assigned manually based on the assigned phylum and according to current eukaryotic phylogeny (Burki et al. 2020) (Supplementary Tables 21–24). Sequences were aligned using MAFFT v.7. as described above. ML phylogenetic inference was performed in IQ-TREE v.2.0.3110 (Nguyen et al. 2015) with automatic model selection in ModelFinder Plus. The optimal model was JTTDCMut+R10 (Kosiol and Goldman 2005; Soubrier et al. 2012). Branch support values were extracted from 1,000 ultrafast BS (Hoang et al. 2018). GraPhlAn v.1.1.3 (Asnicar et al. 2015) was used to generate circular representations of the trees. To generate sequence logos for the C-terminal motif, H2A.X variants scored against pre-built Hidden Markov Models (HMM) were retrieved from the NCBI Histone Variants Database 2.0 (HistoneDB 2.0) (Draizen et al. 2016) for the Chordata, Streptophyta, Mollusca, Platyhelminthes, and Brachiopoda clades (these last three belonging to the Spiralia clade), aligned using MAFFT v.7. as described above. Alignments were trimmed to the last 12 positions of the alignment, corresponding to the positions 131 to 142 of the histone standardised nomenclature for H2A.X, using Jalview v.2.11.2.6 (Waterhouse et al. 2009). Trimmed alignments were turned into information content sequence logos using WebLogo v.2.8.2. (Crooks et al. 2004). Alignments and the corresponding alignment files in .fasta and .aln formats are available in the GitHub repository (see Data Availability section).

### Histone modifier and reader genes mining and orthology assignment

To mine histone modifier orthologs, we split the search into five different analyses corresponding to the five families of interest: histone deacetylases (HDAC), histone acetyltransferases (HAT), type B HAT, histone demethylases (HDM), arginine-specific histone methyltransferases (PRMT), and lysine-specific histone methyltransferases (KMT) (Supplementary Table 25). Gene models corresponding to seven model organisms, namely *Homo sapiens*, *Mus musculus*, *Danio rerio*, *Xenopus tropicalis*, *Caenorhabditis elegans*, *Drosophila melanogaster*, and *Saccharomyces cerevisiae* were downloaded from NCBI and used to create BLAST local nucleotide databases (Camacho et al. 2009), alongside with the ones corresponding to the three annelid taxa of interest (Supplementary Table 26). Histone modifier protein sequences from these seven model organisms (Supplementary Table 27) were used as queries to find annelid orthologs following the mutual best BLAST hit approach (*e* value ≤ 10^-5^) (Supplementary Tables 28–32), obtaining 67, 97, 120, 40, and 155 unique annelid ortholog candidates for the HDAC, HAT, HDM, PRMT, and KMT families, respectively. HAT type A (nuclear) and type B (cytoplasmic) sequences were split into two groups. Appropriate outgroup sequences were chosen for each of the six alignments (Supplementary Table 33). MAFFT v.7 (Katoh et al. 2019) in the L-INS-I iterative refinement method and default scoring parameters were used to generate all six distinct multiple sequence alignments. For orthology assignment, six phylogenetic analyses were performed on selected candidate sequences, which included the longest isoform for each species-gene combination, given that it contained a properly aligned fragment within the family-specific common domain (see below) and that it was not located in a genomic locus of poor sequencing quality. Sequences were trimmed using Jalview v.2.11.2.6 (Waterhouse et al. 2009) to the family-specific domain(s), i.e. the histone deacetylase (DAC; HDAC), *N*-acetyltransferase (NAT; HAT type A and HAT type B), Jumonji C (JmjC; HDM), *S*-adenosylmethionine methyltransferase protein arginine methyltransferase-type (SAM MTase PRMT-type; PRMT), and the Su(var)3-9, Enhancer-of-zeste and Trithorax (SET; KMT) domains using the domain boundaries defined by ProSITE domain annotation for human HDAC11 (UniProt: Q96DB2; HDAC), KAT2A (UniProt: Q92830; HAT type A), NAA40 (UniProt: Q86UY6; HAT type B), KDM2A (UniProt: Q9Y2K7; HDM), PRMT1 (UniProt: Q99873; PRMT), and SUV39H1 (UniProt: O43463; KMT) proteins, respectively. Trimmed alignments were used for ML phylogenetic inference with automatic model selection using ModelFinder Plus in IQ-TREE v.2.0.3110 (Nguyen et al. 2015). The optimal models were LG+R7 (HDAC) (Le and Gascuel 2008; Soubrier et al. 2012), Q.insect+R5 (HAT type A) (Minh et al. 2021), LG+G4 (HAT type B) (Yang 1994), LG+R6 (HDM), LG+F+R5 (PRMT), and LG+R6 (KMT) (Kosiol and Goldman 2005). Bayesian inference in MrBayes v.3.2.7a (Ronquist and Huelsenbeck 2003) was also performed with an LG replacement matrix. Branch support values were extracted from 1,000 ultrafast BS (Hoang et al. 2018), and posterior probabilities were estimated from two runs with four chains of MCMC analyses run for 50,000,000 generations (80,000,000 for KMT) (Supplementary Table 34). All trees were composed and edited in FigTree v.1.4.4 (http://tree.bio.ed.ac.uk/software/figtree/). The resulting histone modifier orthology assignment is summarised in Supplementary Table 35 and described in detail for each species in Supplementary Table 36. Alignments and the corresponding alignment files in .fasta and .nexus formats are available in the GitHub repository (see Data Availability section).

### PRMT6 sequence and architecture analysis

Candidate PRMT6 sequences were manually selected from a BLAST search of *O. fusiformis*’ PRMT6 protein sequence against the non-redundant (nr) protein database. All ten selected metazoan sequences, together with the annelid PRMT6 sequences (Supplementary Table 37), were aligned using MAFFT v.7 (Katoh et al. 2019) in the L-INS-I iterative refinement method and default scoring parameters. Alignment was trimmed to the conserved region of the protein between residues R43 and Y359, using Jalview v.2.11.2.6 (Waterhouse et al. 2009), using the nomenclature of human PRMT6 (UniProt: Q96LA8). Residues contained in the PRMT characteristic motif, the SAM-binding pocket, the arginine-binding pocket, as well as the SAM-interacting and arginine-interacting residues were derived from previous work (Schapira and Freitas 2014; Gupta et al. 2021). Domains and regions with functional relevance were obtained for 11 sequences using InterProScan (Blum et al. 2021). Transcriptomic validation of key genes was performed via direct visualisation of .bam files in Seqmonk v.1.48.1. Alignment and the corresponding alignment files in .fasta and .aln format are available in the GitHub repository (see Data Availability section).

### Protein structure prediction

3D structural models of *O. fusiformis* H2A.X-F and H2A.X-Y proteins, as well as of *O. fusiformis* PRMT6, *C. teleta* PRMT6, and *D. gyrociliatus* PRMT6-A, PRMT6-B, PRMT6-C and PRMT6-D, were created using the AlphaFold Server (https://alphafoldserver.com/about) implementation of AlphaFold 3 (Abramson et al. 2024). Resulting predictions were rendered in PyMol v.3.0.3 (https://pymol.org/), where the relevant phosphate groups were added to Ser139 and Tyr142 of H2A.X-F and H2A.X-Y, where relevant. Protein structure predictions are available in the GitHub repository (see Data Availability section).

### Gene clustering and comparative gene expression analyses

Full transcriptomes were clustered according to their normalised DESeq2 expression dynamics using soft *k-*means clustering (or soft clustering) in the mfuzz v.2.52 package (Futschik and Carlisle 2005) (Supplementary Tables 38–40). Transcripts that were not expressed at any developmental stage were discarded (154 out of 31,979 for *O. fusiformis*, 1,242 out of 41,238 for *C. teleta*, and 200 out of 17,387 for *D. gyrociliatus*). The Calinski-Harabasz index (Caliński and Harabasz 1974) was computed to determine an optimal number of 15 (*O. fusiformis* and *C. teleta*) and eight temporally co-regulated gene clusters (*D. gyrociliatus*) using the NbClust v.3.0.1 package (Charrad et al. 2014). For interspecies comparisons of histone modifiers expression dynamics (Supplementary Tables 41–43), clusters were classed as early/cleavage (*O. fusiformis*: 1–5; *C. teleta*: 1–5; *D. gyrociliatus*: 1–2) or late/post-cleavage (*O. fusiformis*: 7–15; *C. teleta*: 7–15; *D. gyrociliatus*: 4–6). Post-cleavage clusters in *O. fusiformis* and *C. teleta* were further subclassified as pre-larval (*O. fusiformis*: 7–9; *C. teleta*: 7–8) and post-larval (*O. fusiformis*: 11–15; *C. teleta*: 11–15). This way, four different quadrants were rendered for each species pairwise comparisons: early_species A_–early_species B_, early_species A_– late_species B_, late_species A_–early_species B_, and late_species A_–late_species B_. Clusters with peak expression in the female adult of *D. gyrociliatus* (7 and 8) were discarded for these purposes. To construct the gene sets of genes under heterochronic shifts, we only considered the histone modifiers with a single copy ortholog in both *O. fusiformis* and *C. teleta* (for comparisons between *O. fusiformis* and *C. teleta*, Supplementary Table 44) or in all three species (for comparisons between *D. gyrociliatus* and *O. fusiformis* or *C. teleta*, Supplementary Table 45).

### Histones acid extraction

Histones were acid-extracted following an adaptation of a previous protocol (Dickman et al. 2013). 3× *O. fusiformis* adults and 15× *C. teleta* adults were food-deprived for 24 h in artificial seawater (ASW), washed in 1× PBS and spun down for 1.5 min at 3,000 × *g*. Animals were homogenised with a pellet pestle motor in 200 µl Tissue Extraction Buffer (TEB) (0.5 % Triton X-100, 5 mM sodium butyrate, 2 mM phenylmethylsulphonyl fluoride (PMSF), 0.02 % sodium azide, supplemented with 1× cOmplete™ EDTA-free Protease Inhibitor Cocktail Tablets (Roche)). Homogenates were layered over 2.5 ml of a 1.8 M sucrose solution in TEB and centrifuged at 49,000× *g* and 4 °C for 1 h in a rate-zonal centrifugation. Supernatants were removed, and the pellets were resuspended in 0.5 ml TEB, transferred to a clean tube, and pelleted at 21,000 × *g* and 4 °C for 2 min. Histones were extracted from the nuclear pellets in 1.2 ml 0.5 M HCl for 48 h at 4 °C. Crude extracts were centrifuged at 6,500 × *g* and 4 °C for 10 min to remove insoluble debris. 4× cycles of ultrafiltration in Amicon Ultra-0.5 Centrifugal Filter Units with Ultracel-10 regenerated cellulose membranes (10 kDa nominal molecular weight limit, NWML) (Millipore) using mQ water as exchange buffer were performed following the manufacturer’s recommendations, to remove acid and concentrate histones to 20–25 µl. Aliquots were obtained at various steps to assess protein size and purity via sodium dodecyl sulphate-polyacrylamide gel electrophoresis (SDS-PAGE). For *O. fusiformis*, an additional cleaning step was introduced before the ultrafiltration to remove high molecular weight co-extracted proteins using Amicon Ultra-0.5 Centrifugal Filter Units with Ultracel-30 regenerated cellulose membranes (30 kDa, NWML) (Millipore).

### Histone derivatisation and preparation

Histones were prepared for mass spectrometry as previously described (Cole et al. 2021). Briefly, histones underwent chemical derivatisation by adding 10 μl of 100 mM ammonium bicarbonate pH 8.0 and 4 μl of ammonium hydroxide to 10 μg of histone sample then adding 10 μl of propionic anhydride in isopropanol and ammonium hydroxide were then used to maintain a pH higher than 8.0. After a 15-min incubation at 37 °C, samples were dried down in a vacuum centrifuge (Concentrator plus, Eppendorf), and the whole process was repeated. Samples were then re-suspended in 40 μl of 100 mM ammonium bicarbonate and digested with trypsin overnight. Digestion was stopped by adding glacial acetic acid and freezing at −80 °C for 5 min. Samples were vacuum-centrifuged and then underwent two more rounds of proprionylation. Lastly, we used a Hypersep™ Hypercarb™ tip to desalt the samples following the manufacturer’s protocol (Thermo Fisher Scientific) (Minshull et al. 2016).

### Liquid-chromatography tandem-mass spectrometry (LC-MS/MS)

Histone samples were in 0.1 % trifluoroacetic acid (TFA) and analysed on an Ultimate 3000 online nano-LC system with a PepMap300 C18 trapping column (Thermo Fisher Scientific) coupled to a Q Exactive HF Orbitrap (Thermo Fisher Scientific). Peptides were eluted onto a 50 cm × 75 μm Easy-spray PepMap C18 analytical column at 35 °C at a flow rate of 300 nl·min^-1^ using a gradient of 3 % to 25 % over 55 min, and then 25 % to 60 % until 81 min. Loading solvent was 0.1 % TFA and 3 % acetonitrile (ACN), solvent A comprised 0.1 % formic acid (FA) and 3 % ACN, and solvent B was 0.1 % FA and 80 % ACN . Samples were run in data-independent acquisition mode. Histone posttranslational modifications (hPTM) were identified, and their relative abundance was quantified in EpiProfile 2.0, with manual verification of key hPTMs in Skyline, as previously described (Cole et al. 2019) (Supplementary Tables 46, 47).

## Data availability

Supplementary Figures and Supplementary Tables are available in the Supplementary Information section of this manuscript. Accession codes and unique identifiers to previously publicly available datasets we used for this study are listed in Supplementary Table 1 (sequence identifiers used in histone genes mining), Supplementary Table 2 (sequence identifiers used in histone genes orthology assignment), Supplementary Table 8 (ATAC-seq datasets used in chromatin accessibility profiling), Supplementary Table 9 (RNA-seq datasets used in gene expression profiling), Supplementary Table 18 (sequence identifiers used in evolutionary analysis of histone H2A.X variants), Supplementary Table 26 (genome files used in histone modifier genes mining and orthology assignment), Supplementary Table 27 (sequence identifiers used in histone modifier genes mining and orthology assignment) and Supplementary Table 37 (sequence identifiers used in PRMT6 sequence and architecture analysis). The histone variants database, that is, the NCBI Histone Variants Database 2.0 (HistoneDB 2.0), can be found at https://www.ncbi.nlm.nih.gov/research/histonedb/. The AlphaFold Server can be accessed at https://alphafoldserver.com/. Updated filtered gene models, transcripts and protein files, genome annotation reports, alignment files, gene expression files, protein structure predictions, and any other relevant additional files are all publicly available in the GitHub repository associated with this study: https://github.com/ChemaMD/HistonesAnnelida.

## Code availability

All custom code used is available in the GitHub repository associated with this study: https://github.com/ChemaMD/HistonesAnnelida.

## Acknowledgements

We thank Ben Hadley and Natasha Sandikyan for their help with the initial characterisation of histone modifiers, as well as the technical support staff at the Department of Biology at Queen Mary University of London. This study employed computational resources from the High-performance computing facility Apocrita at Queen Mary University of London. This work was funded by grants to José M. Martín-Durán from the European Union Horizon 2020 Framework Programme under the European Research Council (ERC Starting Grant agreement number 801669), to Francisco M. Martín-Zamora (ASSEMBLE Plus Grant agreement number 730984), and to Paul J. Hurd from the Biotechnology and Biological Sciences Research Council (BBSRC Grant agreement numbers BB/L023164/1 and BB/V009311/1).

## Author contributions

José M. Martín-Durán, Paul J. Hurd, and Francisco M. Martín-Zamora conceived and designed the study. Joby Cole and Mark Dickman performed LC-MS/MS analyses. Kero Guynes, Allan M. Carillo-Baltodano, and Rory D. Donnellan contributed to animal culture and *in vitro* fertilisations. Francisco M. Martín-Zamora performed all other lab work and analyses described here and drafted the manuscript. All authors critically read and commented on the manuscript.

