## Supplementary Information for "The evolution and developmental dynamics of histone-based chromatin regulation in Annelida"

**This PDF includes:**

- 31 Supplementary Figures
- 47 Supplementary Tables legends

**Index of Supplementary Figures:**


- **Supplementary Figure 1.** Maximum likelihood phylogeny of histone genes.
- **Supplementary Figure 2.** Bayesian phylogeny of histone genes.
- **Supplementary Figure 3.** Gene structure and genomic organisation of histones.
- **Supplementary Figure 4.** ATAC-seq enrichment in histone genes across annelid development.
- **Supplementary Figure 5.** Histone gene clusters are located in dynamic hyperaccessible chromatin regions.
- **Supplementary Figure 6.** Open chromatin regions are abundant in the loci of histone variants.
- **Supplementary Figure 7.** Expression levels of canonical histones and unknown histone variants.
- **Supplementary Figure 8.** Histone expression dynamics in the development of Annelida.
- **Supplementary Figure 9.** Amplification of H2A.X variants in *O. fusiformis*.
- **Supplementary Figure 10.** The C-terminal region of the H2A.X variant across Metazoa.
- **Supplementary Figure 11.** Evolutionary analysis of H2A.X variants across Eukarya.
- **Supplementary Figure 12.** Maximum likelihood phylogeny of histone deacetylases.
- **Supplementary Figure 13.** Bayesian phylogeny of histone deacetylases.
- **Supplementary Figure 14.** Maximum likelihood phylogeny of histone demethylases.
- **Supplementary Figure 15.** Bayesian phylogeny of histone demethylases.
- **Supplementary Figure 16.** Maximum likelihood phylogeny of type A histone acetyltransferases.
- **Supplementary Figure 17.** Bayesian phylogeny of type A histone acetyltransferases.
- **Supplementary Figure 18.** Maximum likelihood phylogeny of type B histone acetyltransferases.
- **Supplementary Figure 19.** Bayesian phylogeny of type B histone acetyltransferases.
- **Supplementary Figure 20.** Maximum likelihood phylogeny of lysine-specific histone methyltransferases.
- **Supplementary Figure 21.** Bayesian phylogeny of lysine-specific histone methyltransferases.
- **Supplementary Figure 22.** Maximum likelihood phylogeny of arginine-specific methyltransferases.
- **Supplementary Figure 23.** Bayesian phylogeny of arginine-specific methyltransferases.
- **Supplementary Figure 24.** Sequence diversity in the PRMT6 expansions of *D. gyrociliatus*.
- **Supplementary Figure 25.** Domain fusions in the *D. gyrociliatus prmt6-c* gene are likely an artefact.

- **Supplementary Figure 26.** Transcripts clustering according to full RNA-seq time series.
- **Supplementary Figure 27.** Family-wise histone modifier expression dynamics in Annelida.
- **Supplementary Figure 28.** Histone modifiers expression dynamics in Annelida.
- **Supplementary Figure 29.** Gene expression levels of heterochronic histone modifiers correlated with larval type.
- **Supplementary Figure 30.** Gene expression levels of heterochronic histone modifiers correlated with life cycle.
- **Supplementary Figure 31.** LC-MS/MS hPTM quantification in acid-extracted histones from adult annelids.

**Index of Supplementary Tables:**


- **Supplementary Table 1.** Curated histone sequences used for histone genes mining.
- **Supplementary Table 2.** Curated histone sequences used for histone genes orthology assignment.
- **Supplementary Table 3.** Summary of histone ortholog number by histone family and gene.
- **Supplementary Table 4.** Histone gene repertoire in *O. fusiformis*,
- **Supplementary Table 5.** Histone gene repertoire in *C. teleta*.
- **Supplementary Table 6.** Histone gene repertoire in *D. gyrociliatus*.
- **Supplementary Table 7.** Statistics from comparison analysis of histone gene structure.
- **Supplementary Table 8.** ATAC-seq datasets used for chromatin accessibility profiling.
- **Supplementary Table 9.** RNA-seq datasets used for gene expression profiling.
- **Supplementary Table 10.** Stage-specific TPM gene expression matrix for *O. fusiformis*.
- **Supplementary Table 11.** Stage-specific DESeq2 gene expression matrix for *O. fusiformis*.
- **Supplementary Table 12.** Stage-specific TPM gene expression matrix for *C. teleta*.
- **Supplementary Table 13.** Stage-specific DESeq2 gene expression matrix for *C. teleta*.
- **Supplementary Table 14.** Stage-specific TPM gene expression matrix for *D. gyrociliatus*.
- **Supplementary Table 15.** Stage-specific DESeq2 gene expression matrix for *D. gyrociliatus*.
- **Supplementary Table 16.** Primers used for amplification of *h2ax1* (*h2ax-y*) and *h2ax2* (*h2ax-f*) orthologs in *O. fusiformis*.
- **Supplementary Table 17.** Expected amplicon sizes from amplification of *h2ax1* (*h2ax-y*) and *h2ax2* (*h2ax-f*) orthologs in *O. fusiformis*.
- **Supplementary Table 18.** Representative animal H2A.X sequences used to generate a H2A.X MSA.
- **Supplementary Table 19.** List of H2A.X-Y hits recovered from PHI-BLAST search 1.
- **Supplementary Table 20.** List of H2A.X-F hits recovered from PHI-BLAST search 2.
- **Supplementary Table 21.** Taxonomic analysis and classification of species with PHI-BLAST hits.
- **Supplementary Table 22.** Quantification of H2A.X-Y and H2A.X-F variants per kingdom, phylum, and eukaryotic supergroup.
- **Supplementary Table 23.** Number of H2A.X-Y and H2A.X-F variants per hit species.
- **Supplementary Table 24.** List of species encoding both H2A.X variants in their genomes, including their taxonomical information and mode of development.

- **Supplementary Table 25.** List of histone modifier genes studied in our annelid lineages. Standardised gene symbol may sometimes not coincide with the approved gene symbol.
- **Supplementary Table 26.** Genome annotation and gene models files used for histone modifier and reader genes mining.
- **Supplementary Table 27.** Curated histone modifier sequences used for histone modifier genes mining.
- **Supplementary Table 28.** Summary statistics of mutual best hit analysis of HDAC candidates.
- **Supplementary Table 29.** Summary statistics of mutual best hit analysis of HDM candidates.
- **Supplementary Table 30.** Summary statistics of mutual best hit analysis of HAT candidates.
- **Supplementary Table 31.** Summary statistics of mutual best hit analysis of KMT candidates.
- **Supplementary Table 32.** Summary statistics of mutual best hit analysis of PRMT candidates.
- **Supplementary Table 33.** Curated histone modifier and reader sequences used for histone modifier and reader genes orthology assignment.
- **Supplementary Table 34.** Branch support values from maximum likelihood bootstrap values and Bayesian posterior probabilities for histone modifier gene clades.
- **Supplementary Table 35.** Summary of histone modifier number by histone modifier family and clade.
- **Supplementary Table 36.** Analysed histone modifier gene repertoire in all three annelid taxa.
- **Supplementary Table 37.** Representative animal PRMT6 sequences used to generate a PRMT6 MSA and for domain and region architecture prediction.
- **Supplementary Table 38.** Genome annotation based on RNA-seq clusters for *O. fusiformis*.
- **Supplementary Table 39.** Genome annotation based on RNA-seq clusters for *C. teleta*.
- **Supplementary Table 40.** Genome annotation based on RNA-seq clusters for *D. gyrociliatus*.
- **Supplementary Table 41.** RNA-seq cluster assignment, RNA-seq cluster classification, gene annotation, and DESeq2 gene expression of histone modifiers in *O. fusiformis*.
- **Supplementary Table 42.** RNA-seq cluster assignment, RNA-seq cluster classification, gene annotation, and DESeq2 gene expression of histone modifiers in *C. teleta*.
- **Supplementary Table 43.** RNA-seq cluster assignment, RNA-seq cluster classification, gene annotation, and DESeq2 gene expression of histone modifiers in *D. gyrociliatus*.

- **Supplementary Table 44.** Histone modifiers with single-copy orthology between *O. fusiformis* and *C. teleta* used for comparative gene expression analyses.
- **Supplementary Table 45.** Histone modifiers with single-copy orthology between *O. fusiformis*, *C. teleta*, and *D. gyrociliatus*, used for comparative gene expression analyses.
- **Supplementary Table 46.** Sample-specific intensity of peptide/hPTM peaks from LC-MS/MS.
- **Supplementary Table 47.** List of hPTMs detected through LC-MS/MS.

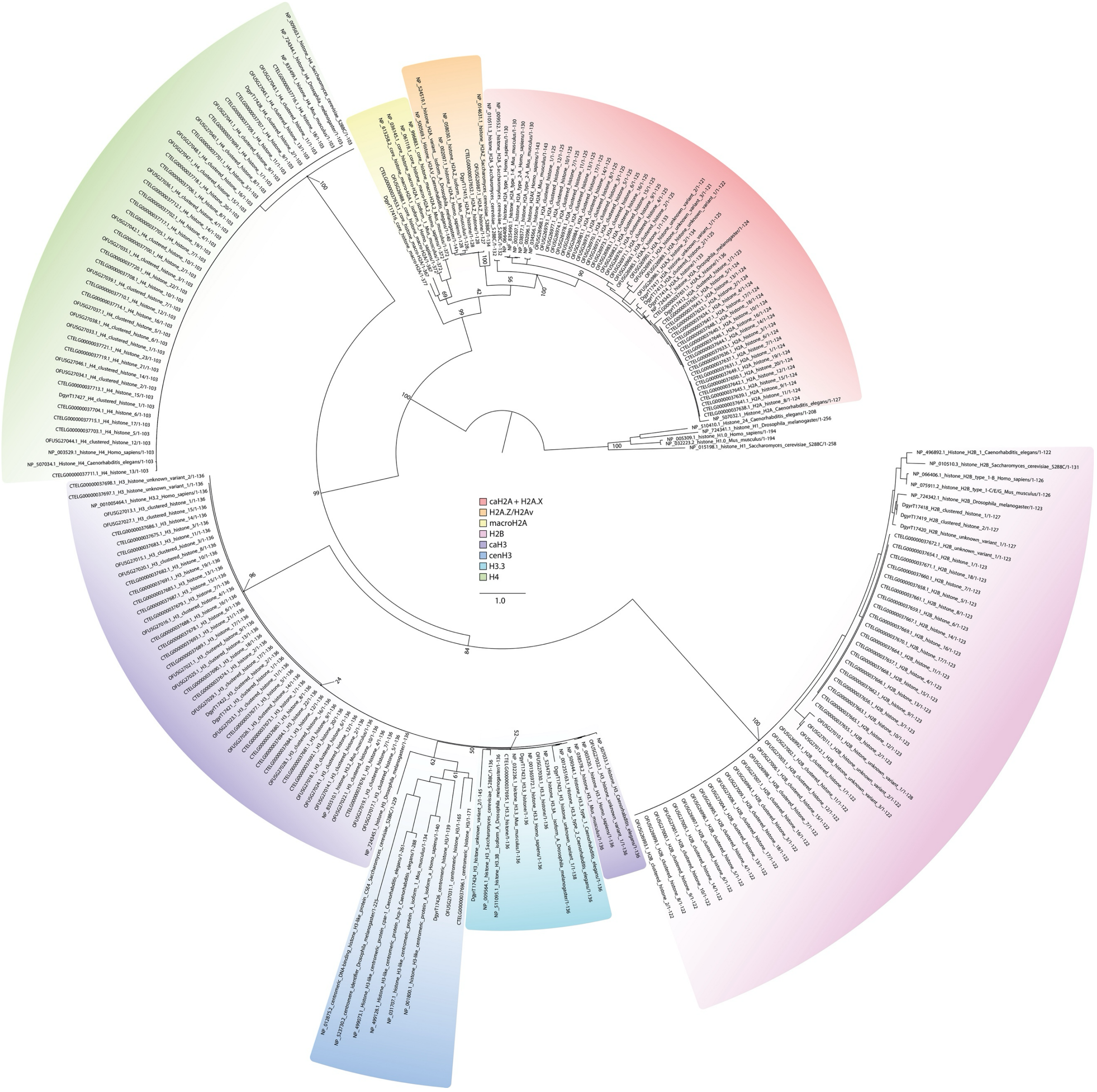


**Supplementary Figure 1 | Maximum likelihood phylogeny of histone genes.**

Maximum likelihood phylogeny for gene orthology analysis of histone genes in *O. fusiformis*, *C. teleta*, and *D. gyrociliatus*. Branch support values represent bootstrap values (0–100 values) at key nodes. Coloured boxes highlight the extent of each histone gene or family. Scale bar depicts the number of amino acid changes per site along the branches. caH2A: canonical H2A; caH3: canonical H3.

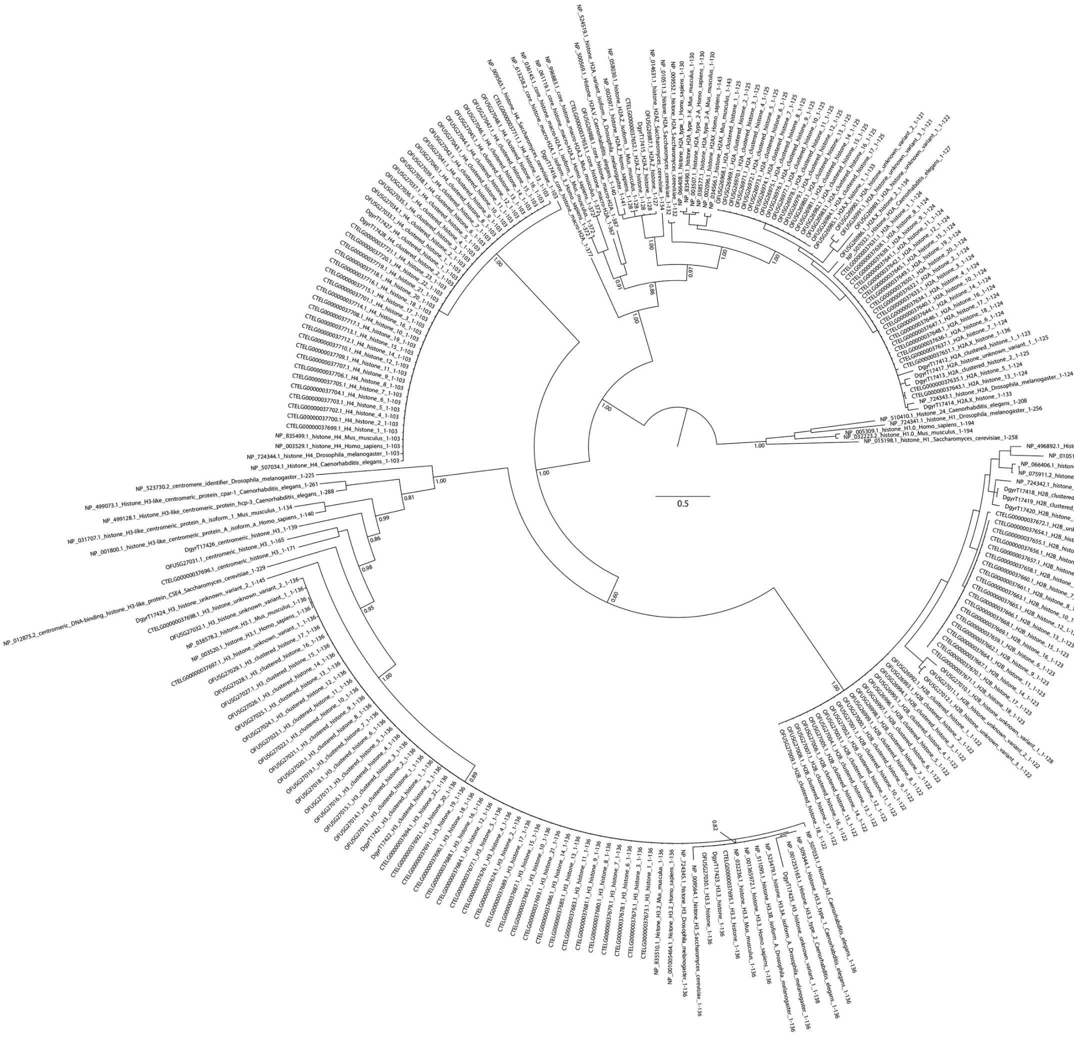

**Supplementary Figure 2 | Bayesian phylogeny of histone genes.**
Bayesian phylogeny for gene orthology analysis of histone genes in *O. fusiformis*, *C. teleta*, and *D. gyrociliatus*. Branch support values represent posterior probabilities (0–1 values) at key nodes. Scale bar depicts the number of amino acid changes per site along the branches. caH2A: canonical H2A; caH3: canonical H3.

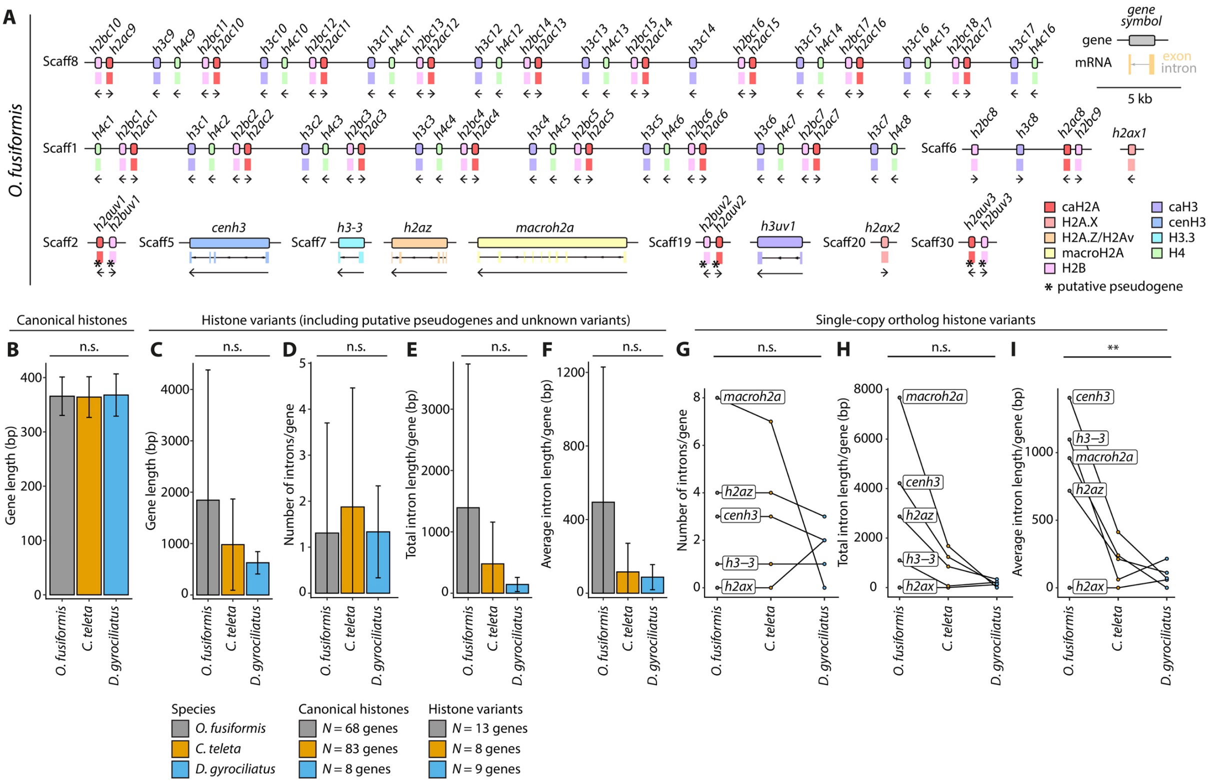

**Supplementary Figure 3 | Gene structure and genomic organisation of histones.**
(**A**) Schematic representation to scale of the genomic loci of the histone genes in *O. fusiformis* in the scaffold-level assembly. Boxes delimitate gene bodies, with the intron-exon composition shown underneath. Arrows below genes indicate direction of transcription. Colours correspond to the different histone genes and gene families. Genes flagged with an asterisk (*) represent putative pseudogenes, as inferred from transcriptomic data (see Supplementary Fig. 7E–G, 8C). Scaff: scaffold. (**B–C**) Gene length of canonical histones (**B**) and all histone variants (**C**) across all three annelid taxa. (**D–I**) Gene-wise number of introns (**D**: all histone variants; **G**: histone variants with inferable orthology), total intron length (**e**: all histone variants; **h**: histone variants with inferable orthology), and average intron length (**F**: all histone variants; **I**: histone variants with inferable orthology), across all three annelid lineages. Data in **G–I** are shown as paired data points. Error bars in **B–F** are standard deviations. *P* values were derived from one-way ANOVAs in **B–F**, or from repeated measures one-way ANOVAs in **G–I**. **: *P* value < 0.01; n.s.: not significant.

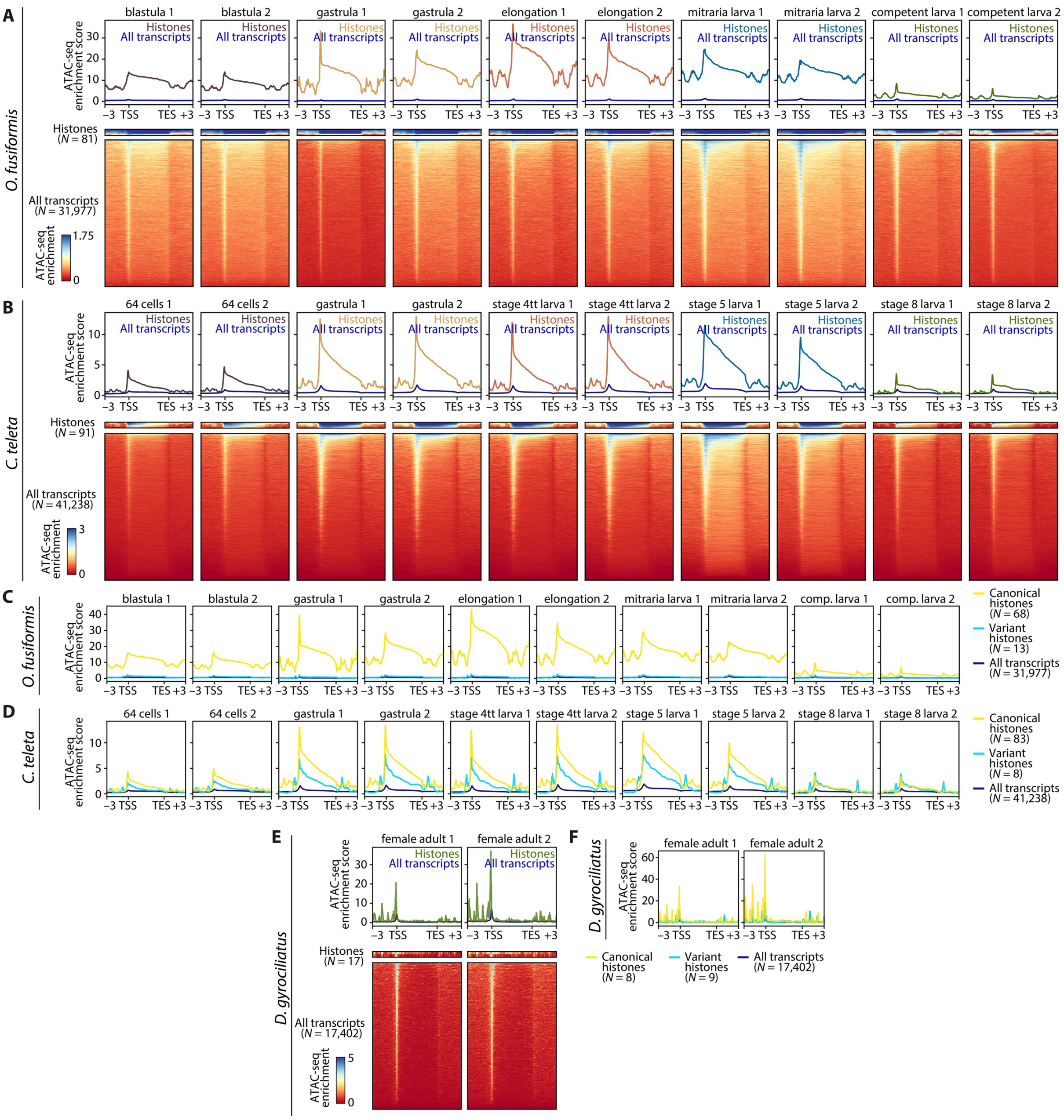


**Supplementary Figure 4 | ATAC-seq enrichment in histone genes across annelid development.**
(**A, B**) ATAC-seq enrichment meta-gene profiles (top) and heatmaps (bottom) of histone genes compared to the whole genome, during the embryonic development of *O. fusiformis* (**A**) and *C. teleta* (**B**). Distances are in kilobases (kb). TSS: transcription start site; TES: transcription end site. (**C, D**) ATAC-seq enrichment meta-gene profiles of canonical (yellow) and variant histones (light blue) compared to the whole genome (dark blue) during the embryonic development of *O. fusiformis* (**C**) and *C. teleta* (**D**) show a predominant enrichment around canonical histones. Colour scale and ATAC-seq enrichment score is shared in each figure panel. (**E**) ATAC-seq enrichment meta-gene profiles (top) and heatmaps (bottom) as in **A** and **B** for the female adult of *D. gyrociliatus*. (**F**) ATAC-seq enrichment meta-gene profiles of canonical (yellow) and variant histones (light blue) compared to the whole genome (dark blue) as in **C** and **D** for the female adult of *D. gyrociliatus*.

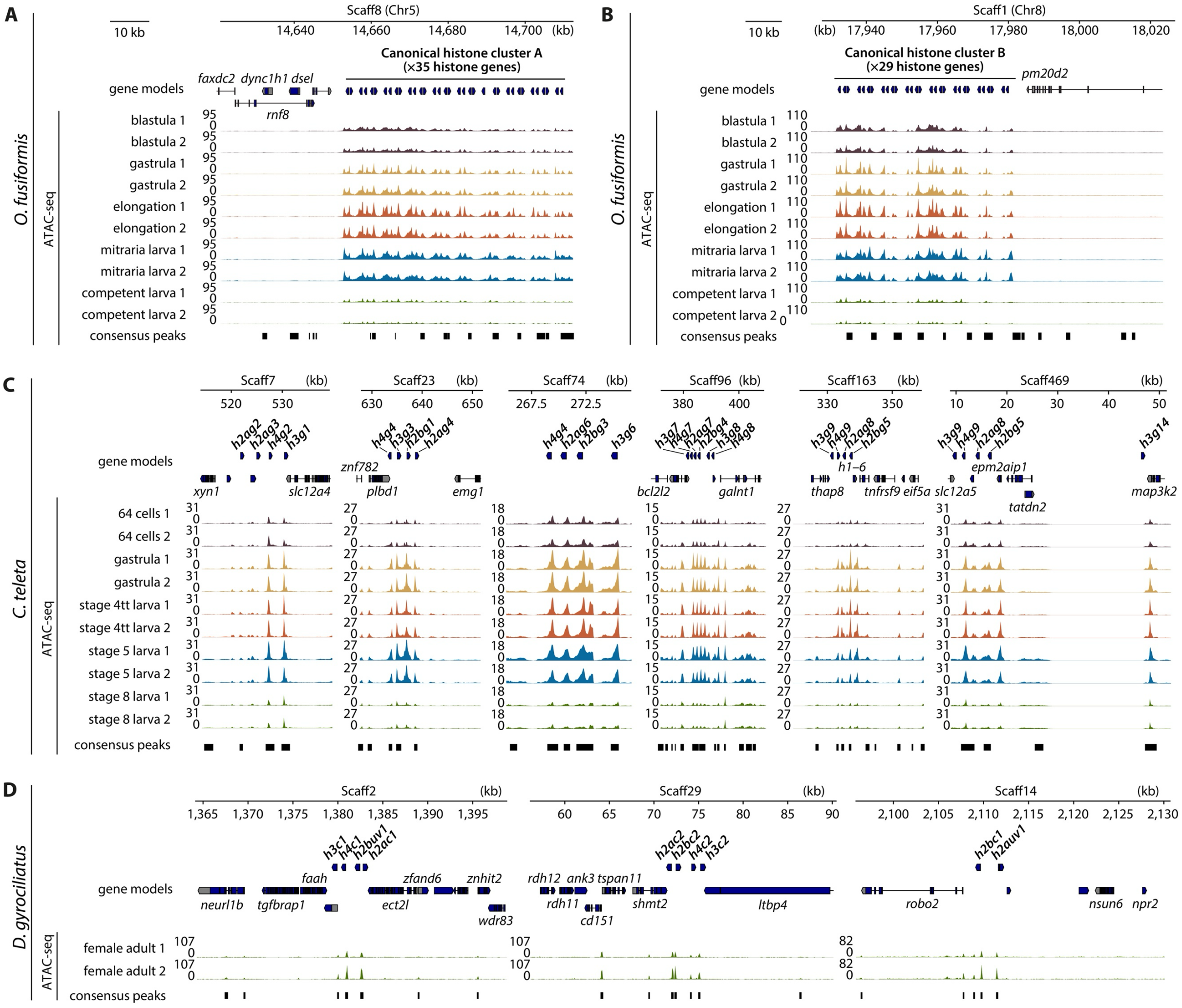


**Supplementary Figure 5 | Histone gene clusters are located in dynamic hyperaccessible chromatin regions.** (**A, B**) ATAC-seq tracks at the large clusters of canonical histones of *O. fusiformis* in chromosome 5 (**A**) and chromosome 8 (**B**). (**C**) ATAC-seq tracks at the six histone gene clusters of *C. teleta* which harbour at least four histones. (**D**) ATAC-seq tracks at the three histone gene clusters of *D. gyrociliatus*. Chr: chromosome; Scaff: scaffold. Note how in all plots the ATAC-seq signal is so high for histone genes that the peaks called in neighbouring genes are not visible at the plot scale. *ank3*: ankyrin 3; *bcl2l2*: bcl2-like 2, *cd151*: CD151 molecule, *dsel*: dermatan sulphate epimerase like; *dync1h1*: dynein cytoplasmic 1 heavy chain 1; *ect2l:* epithelial cell transforming 2 like; *eif5a*: eukaryotic translation initiation factor 5A, *epm2aip1*: EPM2A interacting protein 1, *faah*: fatty acid amide hydrolase; *faxdc2*: fatty acid hydroxylase domain containing 2; *galnt1*: polypeptide *N*-acetylgalactosaminyltransferase 1; *h1–6*: H1.6 linker histone, cluster member; *ltbp4*: latent transforming growth factor beta binding protein 4; *map3k2*: mitogen-activated protein kinase kinase kinase 2; *emg1*: EMG1 N1-specific pseudouridine methyltransferase; *neurl1b*: neutralized E3 ubiquitin protein ligase 1B; *npr2*: natriuretic peptide receptor 2; *nsun6*: NOP2/Sun RNA methyltransferase 6; *plbd1*: phospholipase B domain containing 1; *pm20d2*: peptidase M20 domain containing 2; *rdh12*: retinol dehydrogenase 12; *rdh11*: retinol dehydrogenase 11; *rnf8*: ring finger protein 8; *robo2*: roundabout guidance receptor 2; *shmt2*: serine hydroxymethyltransferase 2; *slc12a4*: solute carrier family 12 member 4; *slc12a5*: solute carrier family 12 member 5; *tatdn2*: TatD DNase domain containing 2; *tgfbrap1*: transforming growth factor beta receptor associated protein 1; *thap8*: THAP domain containing 8; *tnfrsf9*: TNF receptor superfamily member 9; *tspan11*: tetraspanin 11, *wdr83*: WD repeat domain 83, *xyn1*: xylanase 1; *zfand6*: zinc finger AN1-type containing 6; *znf782*: zinc finger protein 782; *znhit2*: zinc finger HIT-type containing 2.

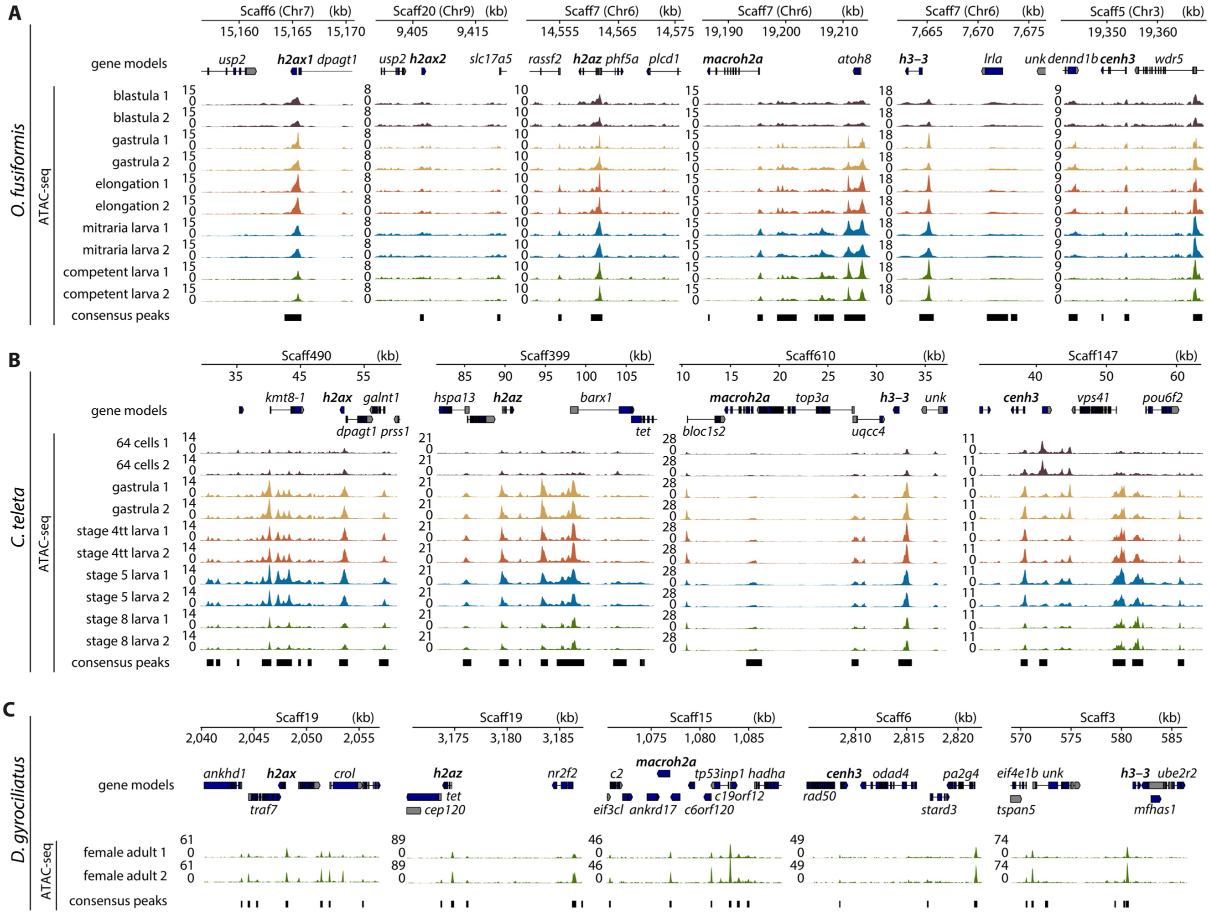


**Supplementary Figure 6 | Open chromatin regions are abundant in the loci of histone variants.** (**A–C**) ATAC-seq tracks at the loci of the histone variants with inferable orthology in *O. fusiformis* (**A**), *C. teleta* (**B**), and *D. gyrociliatus* (**C**). From left to right, *h2ax* (*h2ax1* and *h2ax2* for *O. fusiformis*), *h2az*, *macroh2a*, *h3–3*, and *cenh3*. Unlike what we observed for canonical histones and their clusters (see Supplementary Fig. 5), peaks in or near neighbouring genes to histone variants can be easily observed in most cases, highlighting that the chromatin accessibility levels are lower for variants than for canonical histones for all species. Chr: chromosome; Scaff: scaffold. *ankhd1*: ankyrin repeat and KH domain containing 1; *ankrd17*: ankyrin repeat domain 17; *atoh8*: atonal bHLH transcription factor 8; *barx1*: BARX homeobox 1; *bloc1s2*: biogenesis of lysosomal organelles complex 1 subunit 2; *c2*: complement C2; *c6orf120*: chromosome 6 open reading frame 120; *c19orf12*: chromosome 9 open reading frame 12; *cep120*: centrosomal protein 120; *crol*: crooked legs; *dennd1b*: DENN domain containing 1B; *dpagt1*: dolichyl-phosphate *N*-acetylglucosaminephosphotransferase 1; *eif3cl*: eukaryotic translation initiation factor 3 subunit C like; *eif4e1b*: eukaryotic translation initiation factor 4E family member 1B; *galnt1*: polypeptide *N*-acetylgalactosaminyltransferase 1; *hadha*: hydroxyacyl-CoA dehydrogenase trifunctional multienzyme complex subunit alpha; *hspa13*: heat shock protein family A (Hsp70) member 13; *kmt8-1*: lysine N-methyltransferase 8-1; *lrla*: latrophilin receptor-like protein A; *mfhas1*: multifunctional ROCO family signalling regulator 1; *nr2f2*: nuclear receptor subfamily 2 group F member 2; *odad4*: outer dynein arm docking complex subunit 4; *pa2g4*: proliferation-associated 2G4; *phf5a*: PHD finger protein 5A; *plcd1*: phospholipase C delta 1; *pou6f2*: POU class 6 homeobox 2; *prss1*: serine protease 1; *rad50*: RAD50 double strand break repair protein; *rassf2*: Ras association domain family member 2; *slc17a5*: solute carrier family 17 member 5; *stard3*: StAR related lipid transfer domain containing 3; *tet*: Tet methylcytosine dioxygenase; *top3a*: DNA topoisomerase III alpha; *tp53inp1*: tumor protein p53 inducible nuclear protein 1; *traf7*: TNF receptor associated factor 7; *tspan5*: tetraspanin 5; *ube2r2*: ubiquitin conjugating enzyme E2 R2; *unk*: Unk zinc finger; *uqcc4*: ubiquinol-cytochrome c reductase complex assembly factor 4; *usp2*: ubiquitin specific peptidase 2; *vps41*: VPS41 subunit of HOPS complex; *wdr5*: WD repeat domain 5.

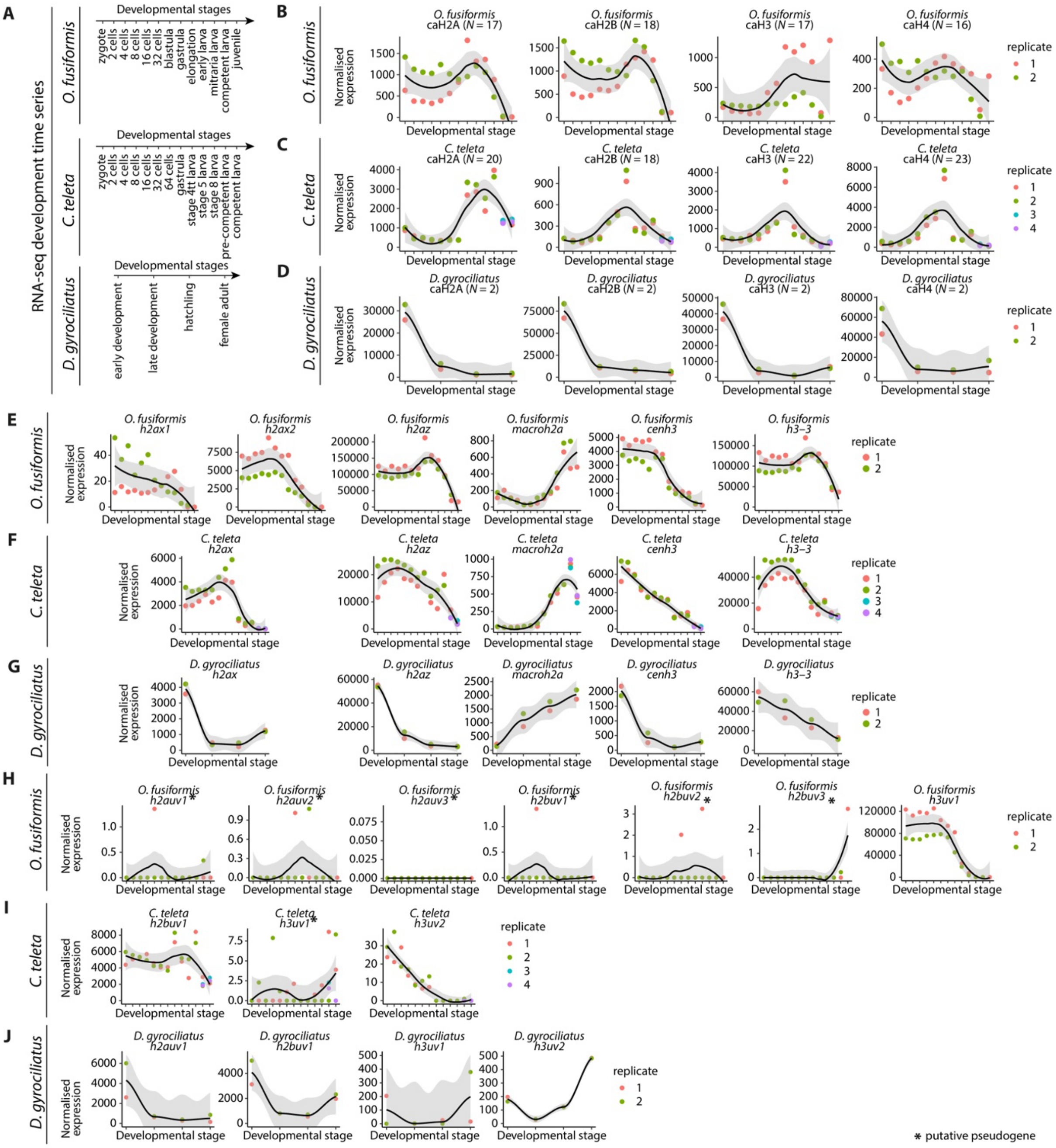


**Supplementary Figure 7 | Expression levels of canonical histones, variant histones, and unknown histone variants.**

(**A**) Time points corresponding to the RNA-seq development time series for *O. fusiformis* (top), *C. teleta* (centre), and *D. gyrociliatus* (bottom). (**B–D**) Total normalised expression levels (summed) of all canonical histones, classified into caH2A, caH2B, caH3, and caH4 families, during the development of *O. fusiformis* (**B**), *C. teleta* (**C**), and *D. gyrociliatus* (**D**). (**E–G**) Normalised expression levels of *h2ax*, *h2az*, *macroh2a*, *cenh3*, and *h3–3* variant histone genes, for *O. fusiformis* (**E**), *C. teleta* (**F**), and *D. gyrociliatus* (**G**). (**H–J**) Normalised expression levels of unknown variants during the development of *O. fusiformis* (**H**), *C. teleta* (**I**), and *D. gyrociliatus* (**J**). Genes with expression levels below 10 (in DESeq2 normalised units) all throughout the animals' development were deemed putative pseudogenes and are shown here flagged with asterisk (*). Curves in **B–J** are locally estimated scatterplot smoothings, coloured shaded areas represent standard error of the mean.

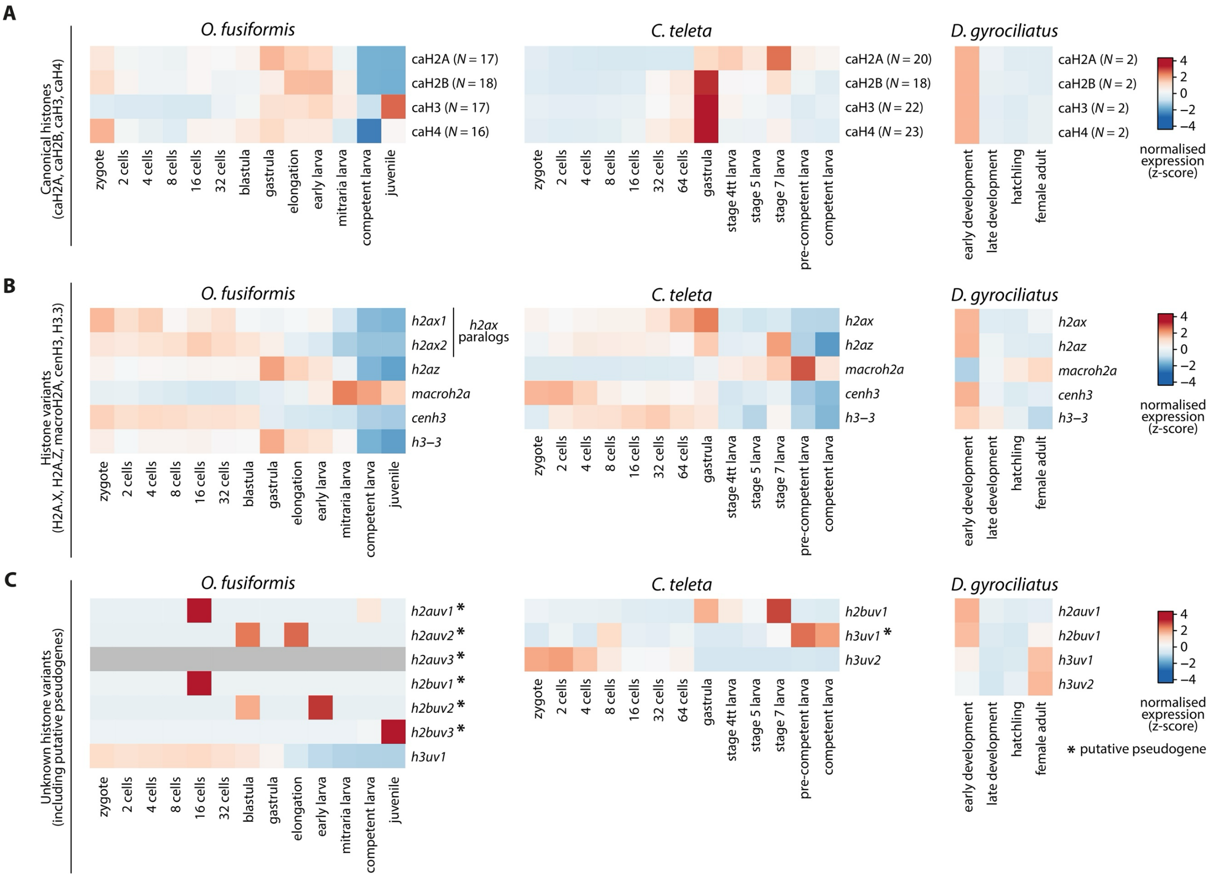


**Supplementary Figure 8 | Histone expression dynamics in the development of Annelida.**
(**A–C**) Expression dynamics of canonical histones (**A**), histone variants with inferable orthology (**B**), and unknown variants (**C**), across the development of *O. fusiformis* (left), *C. teleta* (centre), and *D. gyrociliatus* (right). Dynamics of canonical histones in **a** were derived from total levels obtained from adding all canonical histones' expression levels together. Colour scale denotes normalised gene expression, in a z-score scale. Genes in **c** flagged with an asterisk (*) represent putative pseudogenes.

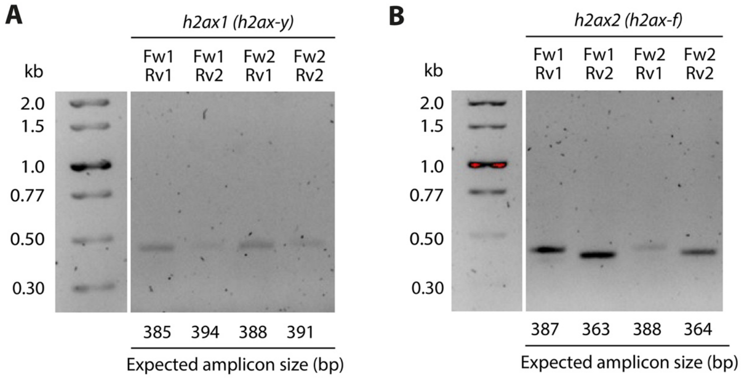


**Supplementary Figure 9 | Amplification of H2A.X variants in *O. fusiformis*.**
(**A**, **B**) Agarose electrophoresis of close-to-full-length amplified fragments of *h2ax1* (*h2ax-y*, **A**) and *h2ax2* (*h2ax-f*, **B**) variants of *O. fusiformis*. cDNA was a pool of cDNAs from multiple time points of the development of *O. fusiformis*. Given the high sequence identity between both orthologs, a strategy involving four different combinations of two gene-specific primers for each gene were used to confirm the specific amplification of the genes of interest. Note that the bands sizes correspond to the expected amplicon sizes for each primer combination. Primers are listed in Supplementary Table 3.16.

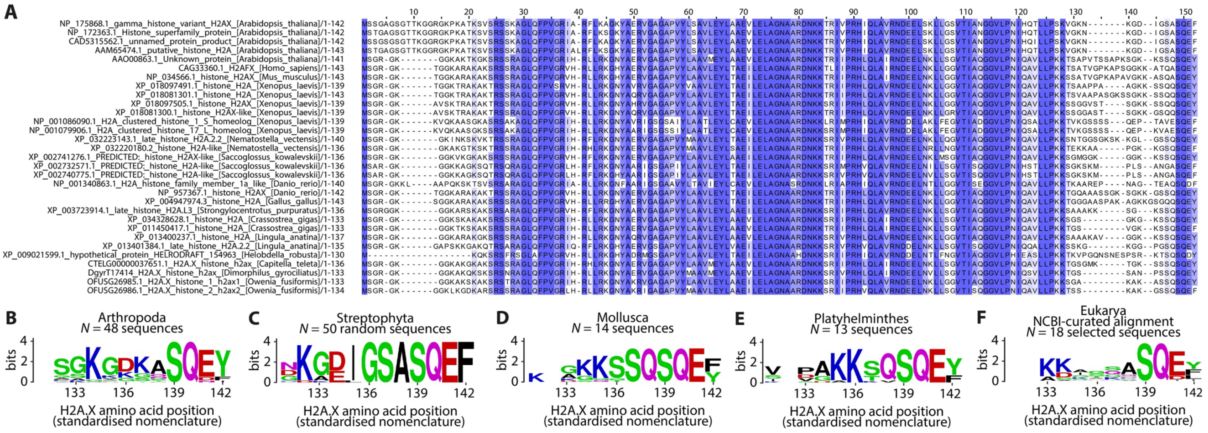


**Supplementary Figure 10 | The C-terminal region of the H2A.X variant across Metazoa.**
(**A**) Full-length MSA of selected H2A.X proteins shown in Fig. 4C. Residue of interest Y142 is in position 152 in the MSA. Blue gradient represents the sequence identity for each position in the alignment. (**B–F**) Sequence logos of the C-terminal region (positions 131–142) of the 48 curated arthropod H2A.X sequences (**B**), 50 random streptophytes H2A.X sequences (**C**), the 14 curated molluscan H2A.X sequences (**D**), the 13 curated platyhelminth H2A.X sequences (**E**), and an NCBI-curated alignment from 18 representative eukaryote sequences (**F**), obtained from the HistoneDB 2.0 database (Draizen et al. 2016).

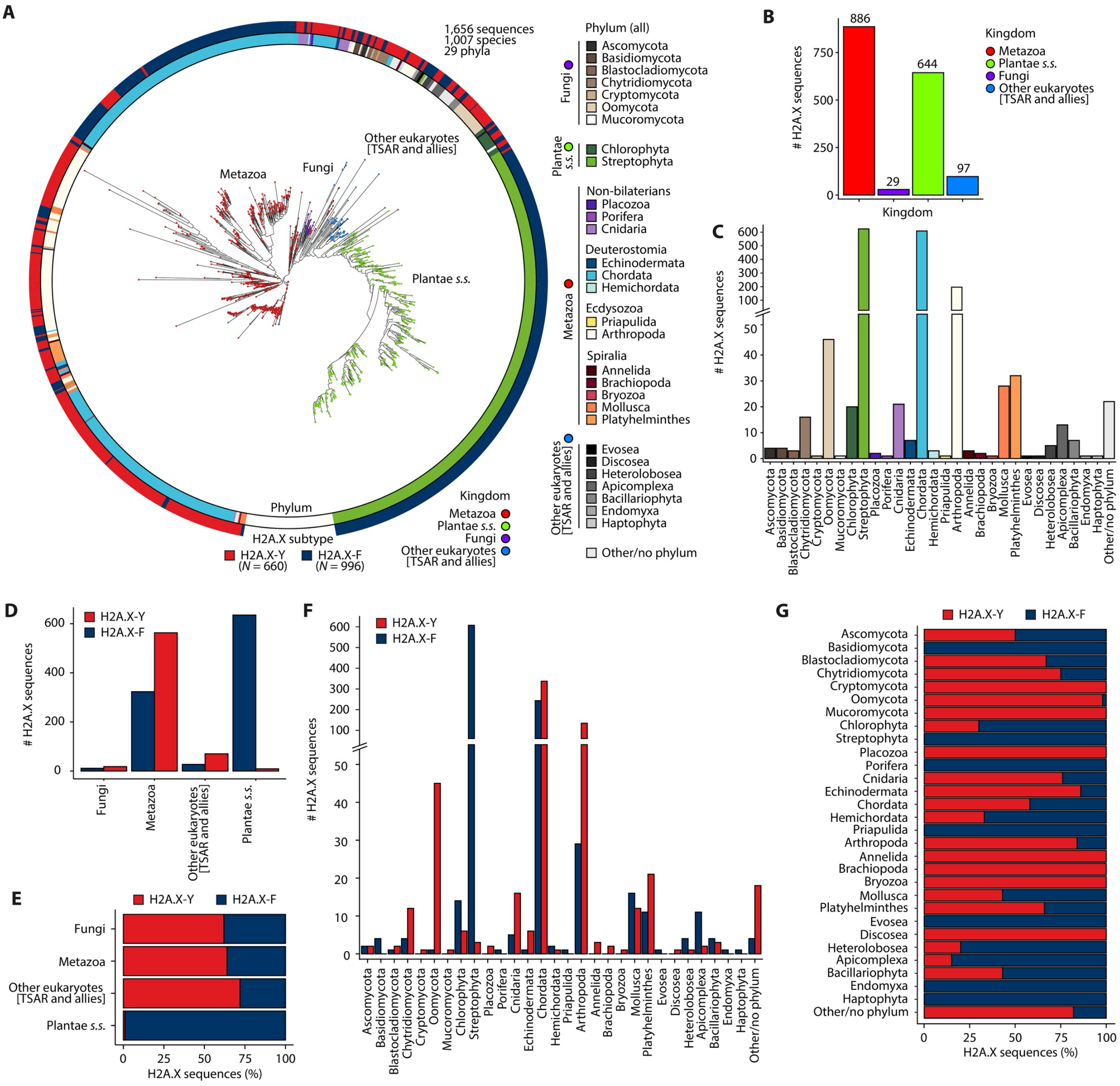


**Supplementary Figure 11 | Evolutionary analysis of H2A.X variants across Eukarya.**
(**A**) Maximum likelihood evolutionary reconstruction of PHI-BLAST-retrieved H2A.X-Y and H2A.X-F variants. Phylum of sequences is shown in the inner circle, H2A.X subtype/variant is shown in the outer circle, in colour coded scales. *s.s.*: *sensu stricto*; TSAR: Telonemia, Stramenopiles, Alveolata, and Rhizaria. (**B**, **C**) Number of retrieved H2A.X sequences per kingdom (**B**) and per phylum (**C**). (**D**, **E**) Number (**D**) and percentage (**E**) of H2A.X sequences per kingdom, classified by subtype (H2A.X-Y and H2A.X-F). (**F**, **G**) Number (**F**) and percentage (**G**) of H2A.X sequences per phylum, classified by subtype (H2A.X-Y and H2A.X-F).

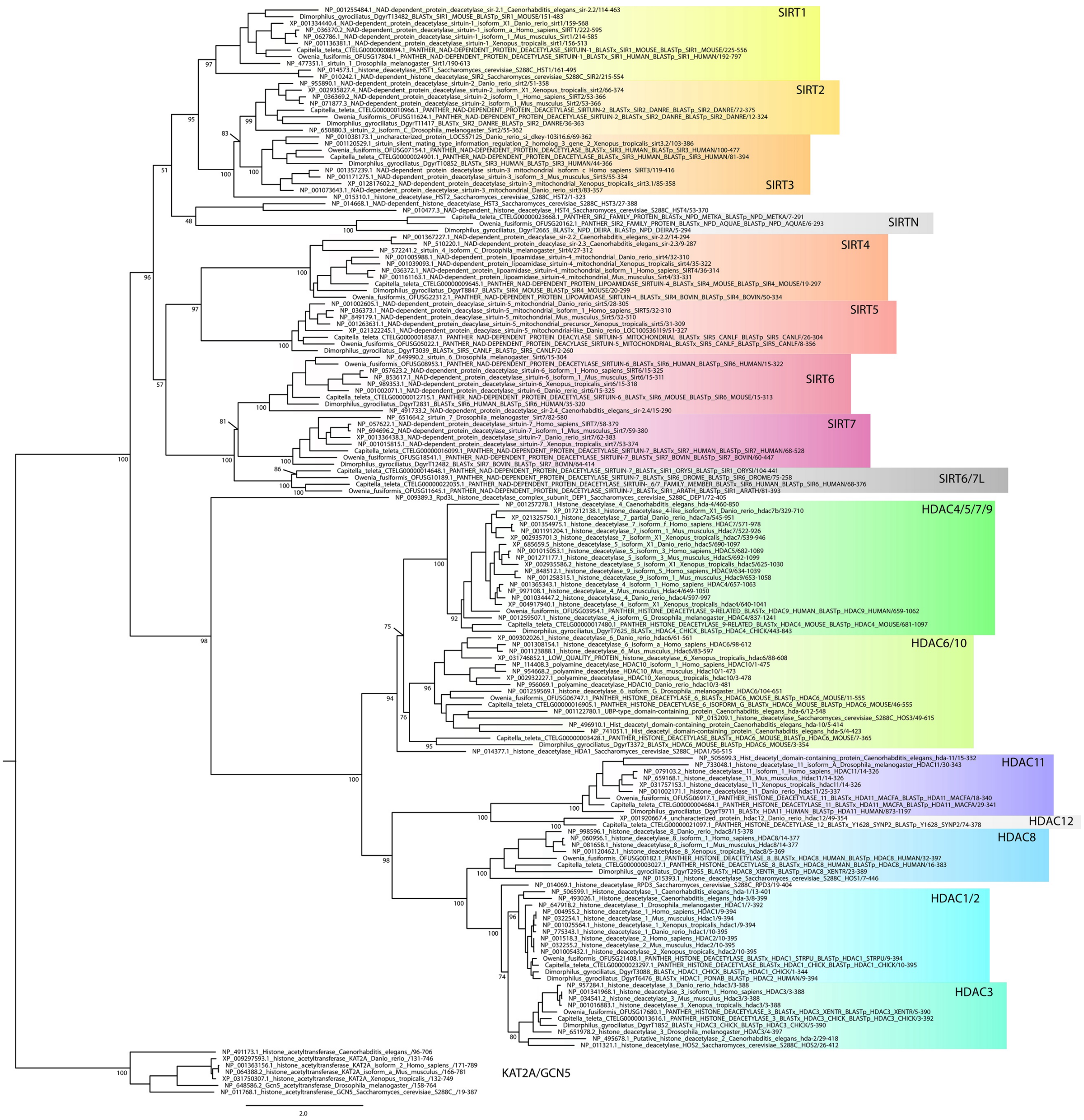


**Supplementary Figure 12 | Maximum likelihood phylogeny of histone deacetylases.**
Maximum likelihood phylogeny for gene orthology assignment of HDAC genes in *O. fusiformis*, *C. teleta*, and *D. gyrociliatus*. Branch support values represent bootstrap values (0–100 values) at key nodes. Coloured boxes highlight the extent of each HDAC clade. Some protein symbols are custom for annelid or lineage-specific clades, as described in text (e.g., SIRT6/7L). Orthologs to more than 1 gene in mammals are assigned as a single one, separated by strokes (e.g., HDAC1/2). Scale bar depicts the number of amino acid changes per site along the branches.

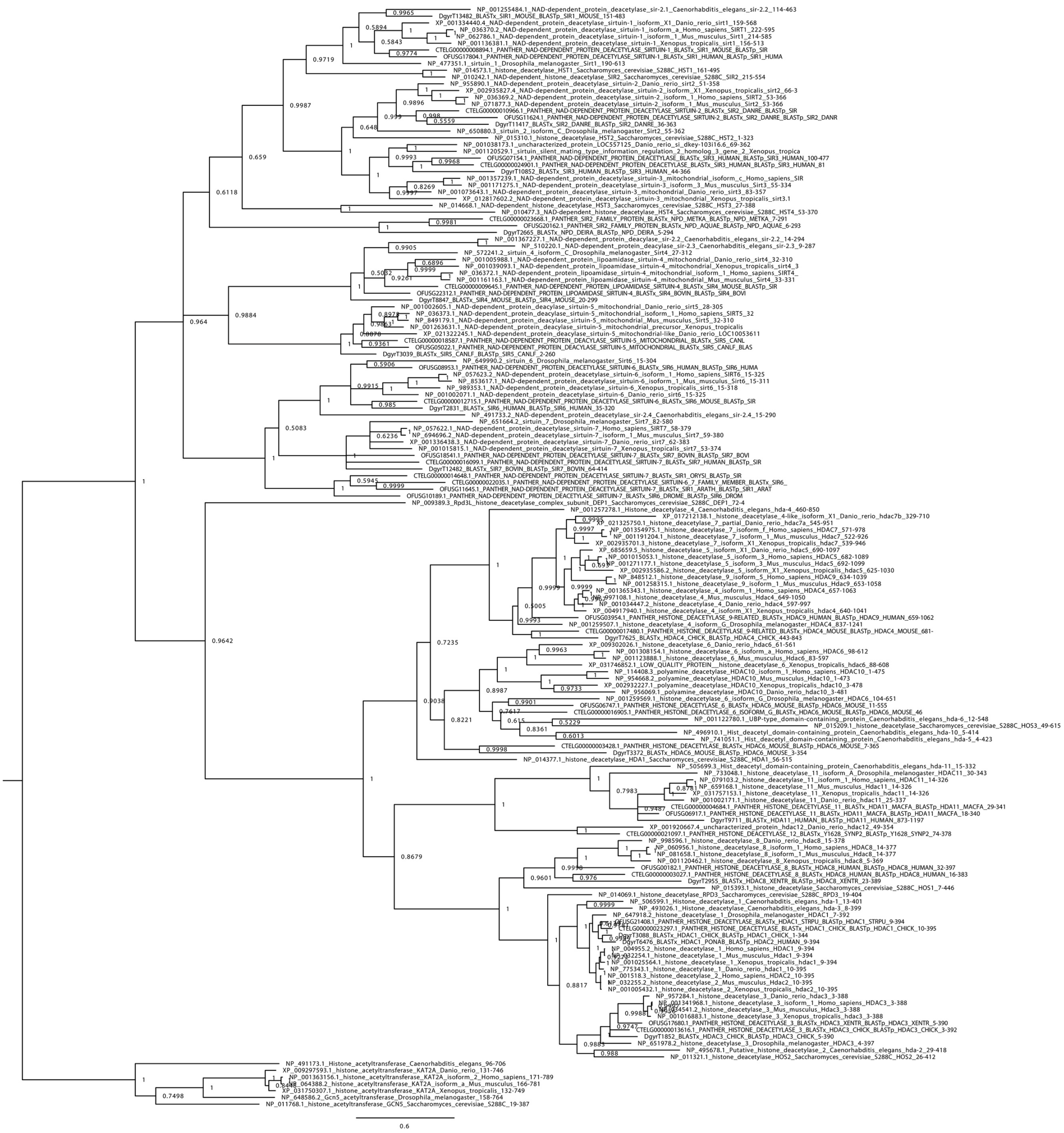

**Supplementary Figure 13 | Bayesian phylogeny of histone deacetylases.**

Bayesian phylogeny for gene orthology assignment of HDAC genes in *O. fusiformis*, *C. teleta*, and *D. gyrociliatus*. Branch support values represent posterior probabilities (0–1 values) at each node. Scale bar depicts the number of amino acid changes per site along the branches.

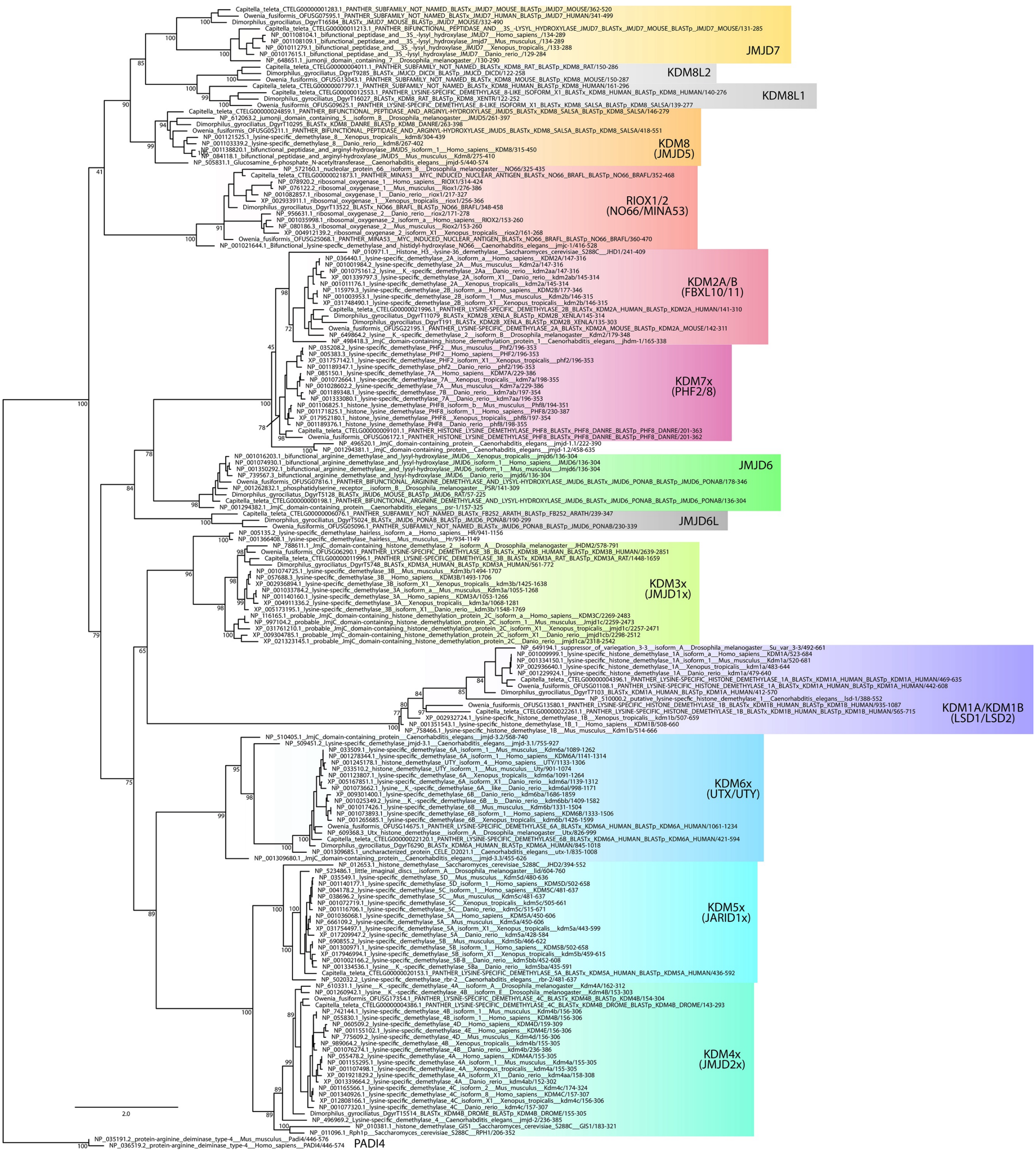


**Supplementary Figure 14 | Maximum likelihood phylogeny of histone demethylases.** Maximum likelihood phylogeny for gene orthology assignment of HDM genes in *O. fusiformis*, *C. teleta*, and *D. gyrociliatus*. Branch support values represent bootstrap values (0–100 values) at key nodes. Coloured boxes highlight the extent of each HDM clade. Some protein symbols are custom for annelid or lineage-specific clades, as described in text (e.g., KDM8L1). Orthologs to more than 1 gene in mammals are assigned as a single one, separated by strokes (e.g., RIOX1/2) or replaced by an x where the numbers would normally be (e.g., KDM3x). Scale bar depicts the number of amino acid changes per site along the branches.

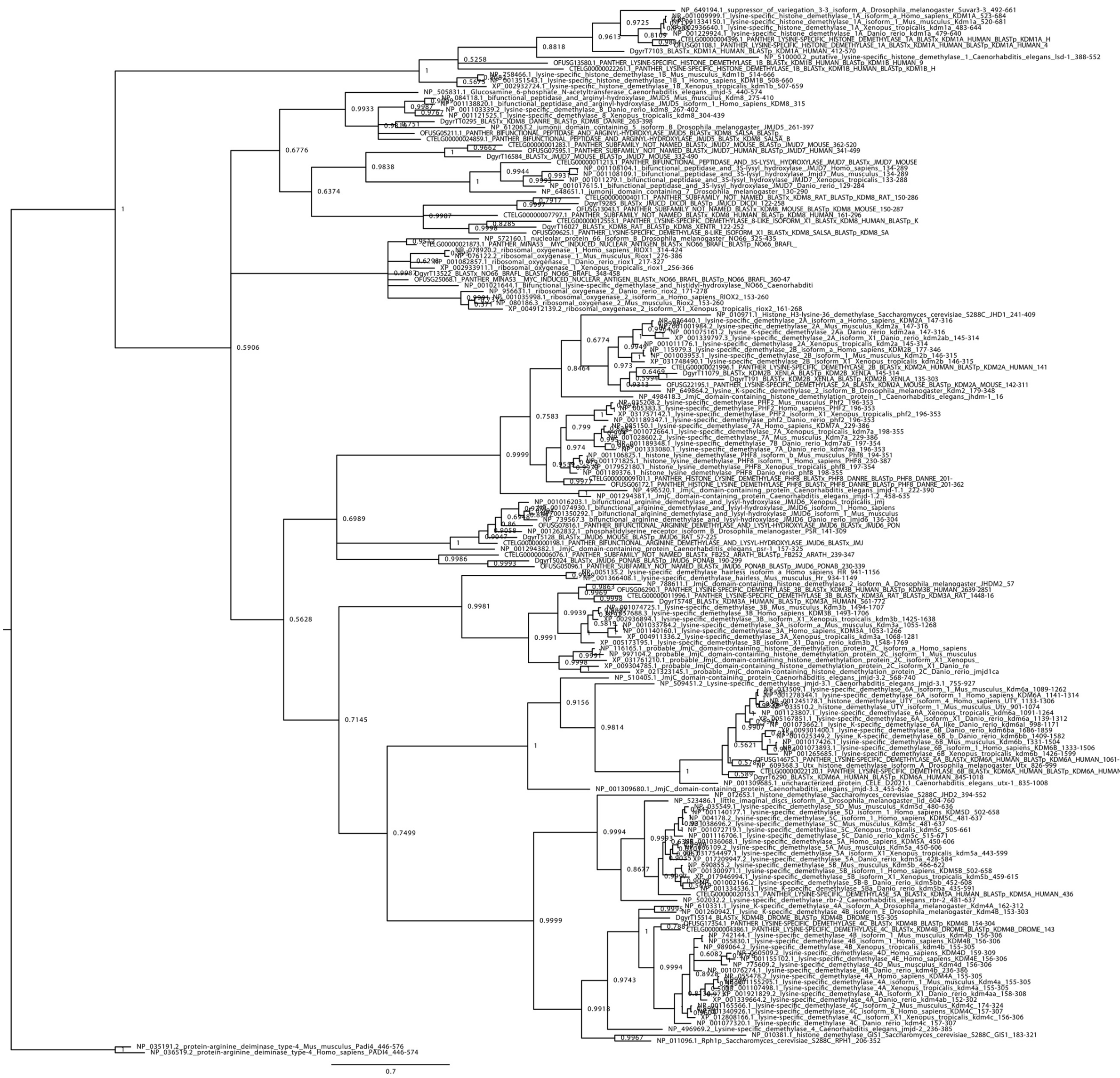

**Supplementary Figure 15 | Bayesian phylogeny of histone demethylases.**
Bayesian phylogeny for gene orthology assignment of HDM genes in *O. fusiformis*, *C. teleta*, and *D. gyrociliatus*. Branch support values represent posterior probabilities (0–1 values) at each node. Scale bar depicts the number of amino acid changes per site along the branches.

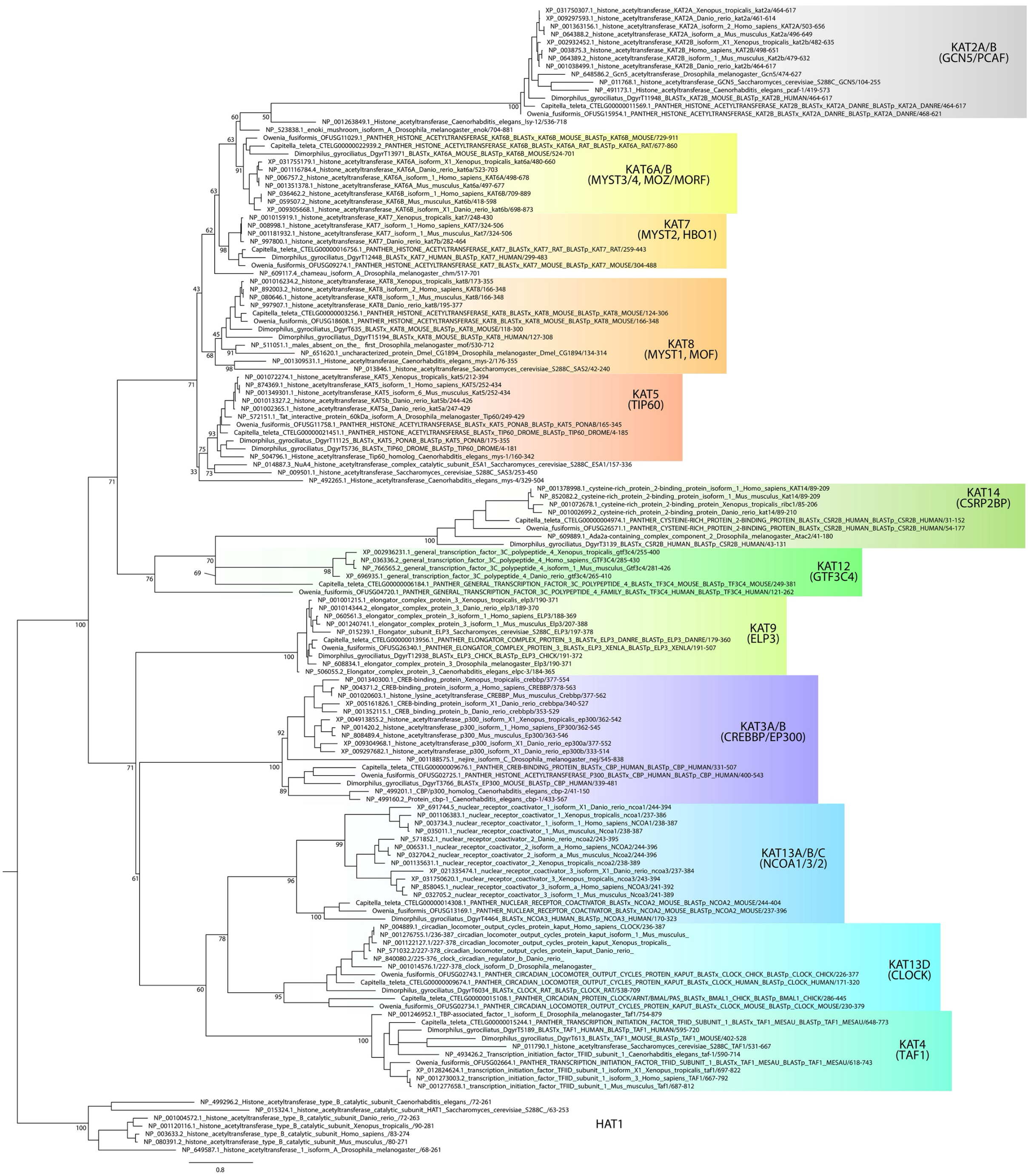


**Supplementary Figure 16 | Maximum likelihood phylogeny of type A histone acetyltransferases.**
Maximum likelihood phylogeny for gene orthology assignment of type A HAT genes in *O. fusiformis*, *C. teleta*, and *D. gyrociliatus*. Branch support values represent bootstrap values (0–100 values) at key nodes. Coloured boxes highlight the extent of each type A HAT clade. Orthologs to more than 1 gene in mammals are assigned as a single one, separated by strokes (e.g., KAT2A/B). Scale bar depicts the number of amino acid changes per site along the branches.

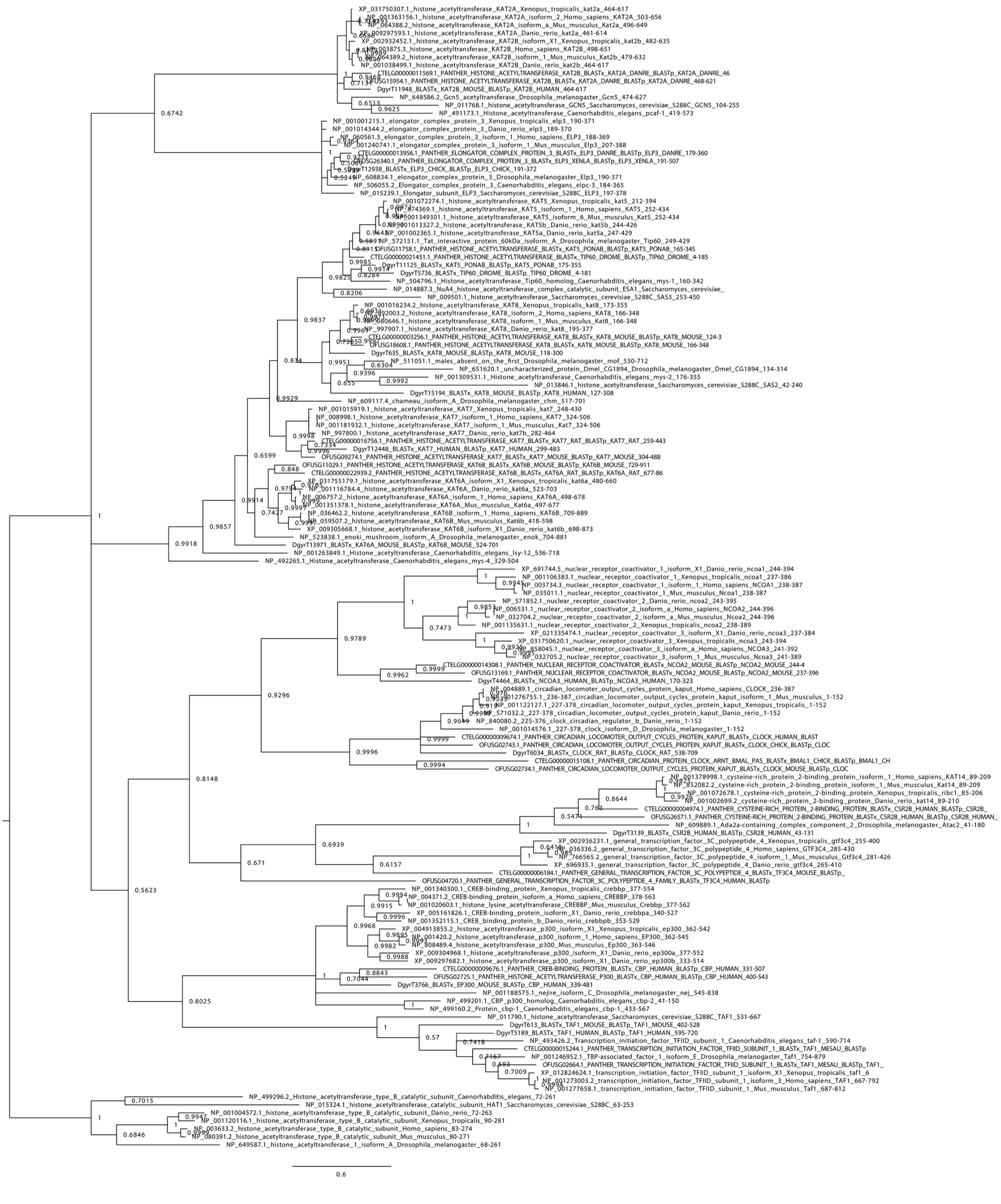

**Supplementary Figure 17 | Bayesian phylogeny of type A histone acetyltransferases.**
Bayesian phylogeny for gene orthology assignment of type A HAT genes in *O. fusiformis*, *C. teleta*, and *D. gyrociliatus*. Branch support values represent posterior probabilities (0–1 values) at each node. Scale bar depicts the number of amino acid changes per site along the branches.

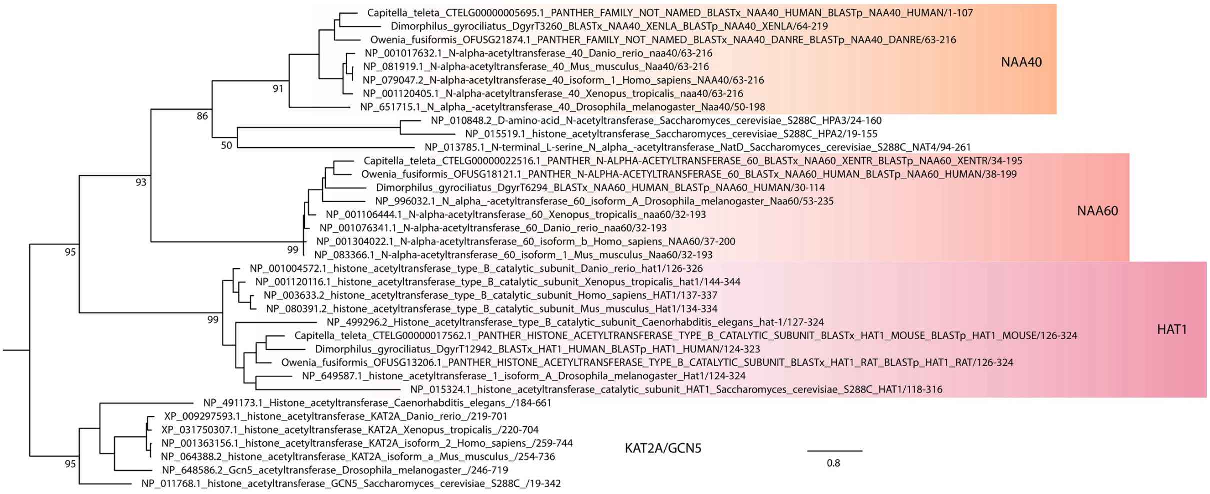


**Supplementary Figure 18 | Maximum likelihood phylogeny of type B histone acetyltransferases.**
Maximum likelihood phylogeny for gene orthology assignment of type B HAT genes in *O. fusiformis*, *C. teleta*, and *D. gyrociliatus*. Branch support values represent bootstrap values (0–100 values) at key nodes. Coloured boxes highlight the extent of each type B HAT clade. Scale bar depicts the number of amino acid changes per site along the branches.

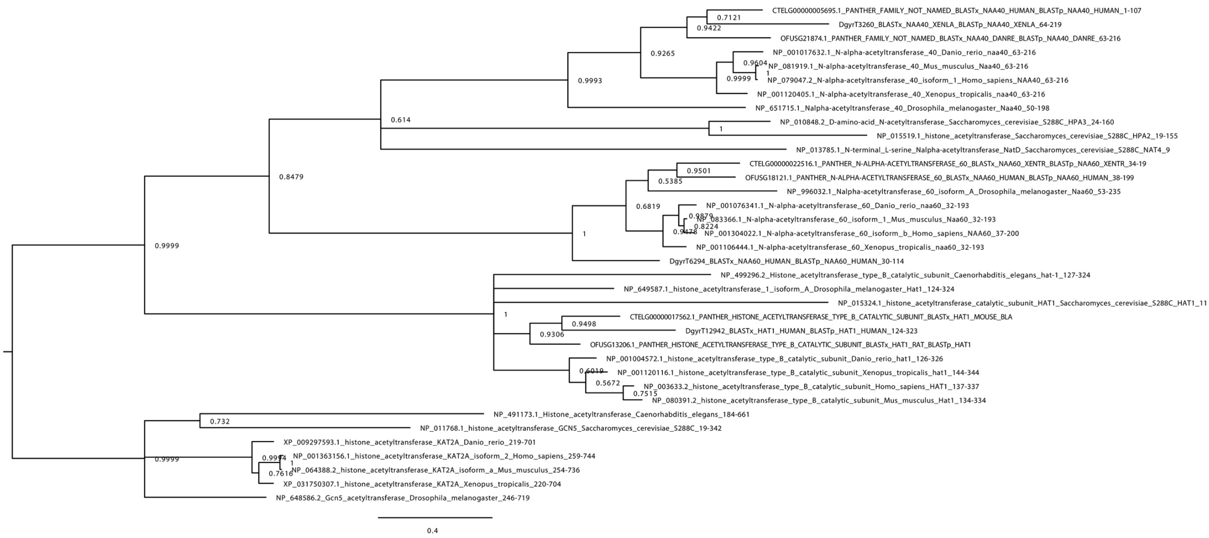


**Supplementary Figure 19 | Bayesian phylogeny of type B histone acetyltransferases.**
Bayesian phylogeny for gene orthology assignment of type B HAT genes in *O. fusiformis*, *C. teleta*, and *D. gyrociliatus*. Branch support values represent posterior probabilities (0–1 values) at each node. Scale bar depicts the number of amino acid changes per site along the branches.

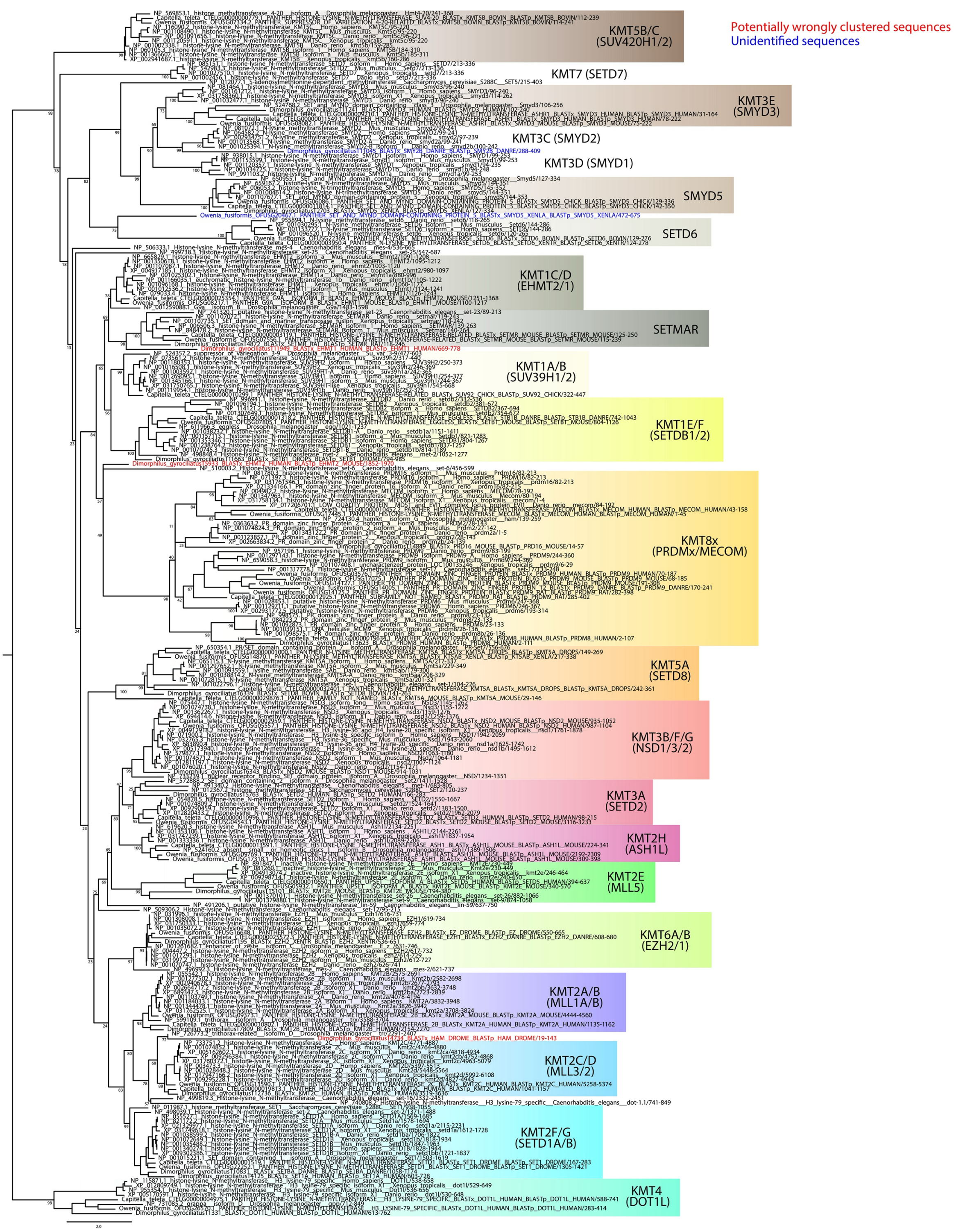


**Supplementary Figure 20 | Maximum likelihood phylogeny of lysine-specific histone methyltransferases.** Maximum likelihood phylogeny for gene orthology assignment of KMT genes in *O. fusiformis*, *C. teleta*, and *D. gyrociliatus*. Branch support values represent bootstrap values (0–100 values) at key nodes. Coloured boxes highlight the extent of each KMT clade. Scale bar depicts the number of amino acid changes per site along the branches. Orthologs to more than 1 gene in mammals are assigned as a single one, separated by strokes (e.g., KMT3B/F/G) or replaced by an x where the numbers would normally be (e.g., KMT8x). Potentially wrongly clustered sequences and unidentified proteins are shown in red and blue font, respectively. Scale bar depicts the number of amino acid changes per site along the branches.

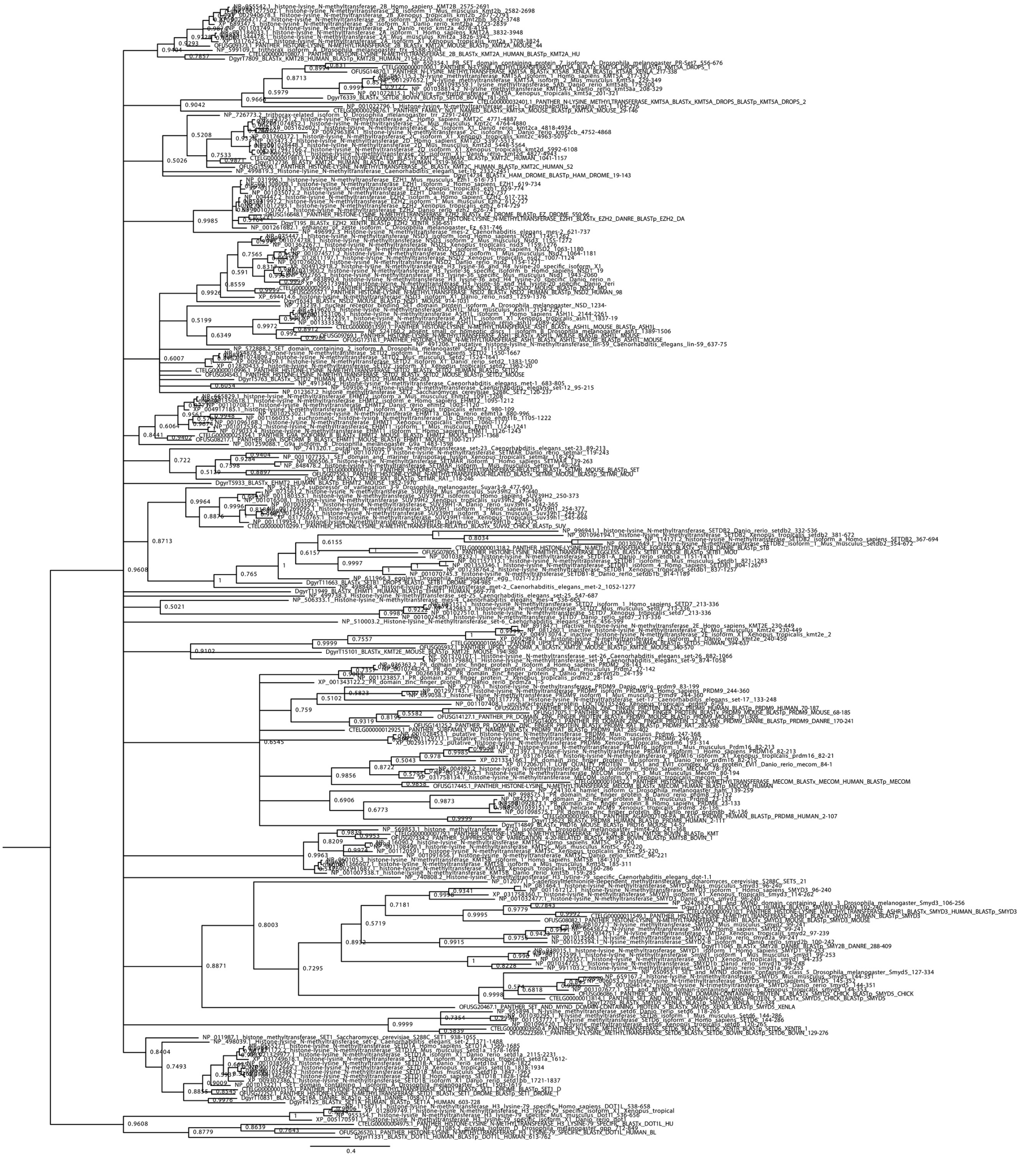

**Supplementary Figure 21 | Bayesian phylogeny of lysine-specific histone methyltransferases.** Bayesian phylogeny for gene orthology assignment of KMT genes in *O. fusiformis*, *C. teleta*, and *D. gyrociliatus*. Branch support values represent posterior probabilities (0–1 values) at each node. Scale bar depicts the number of amino acid changes per site along the branches.

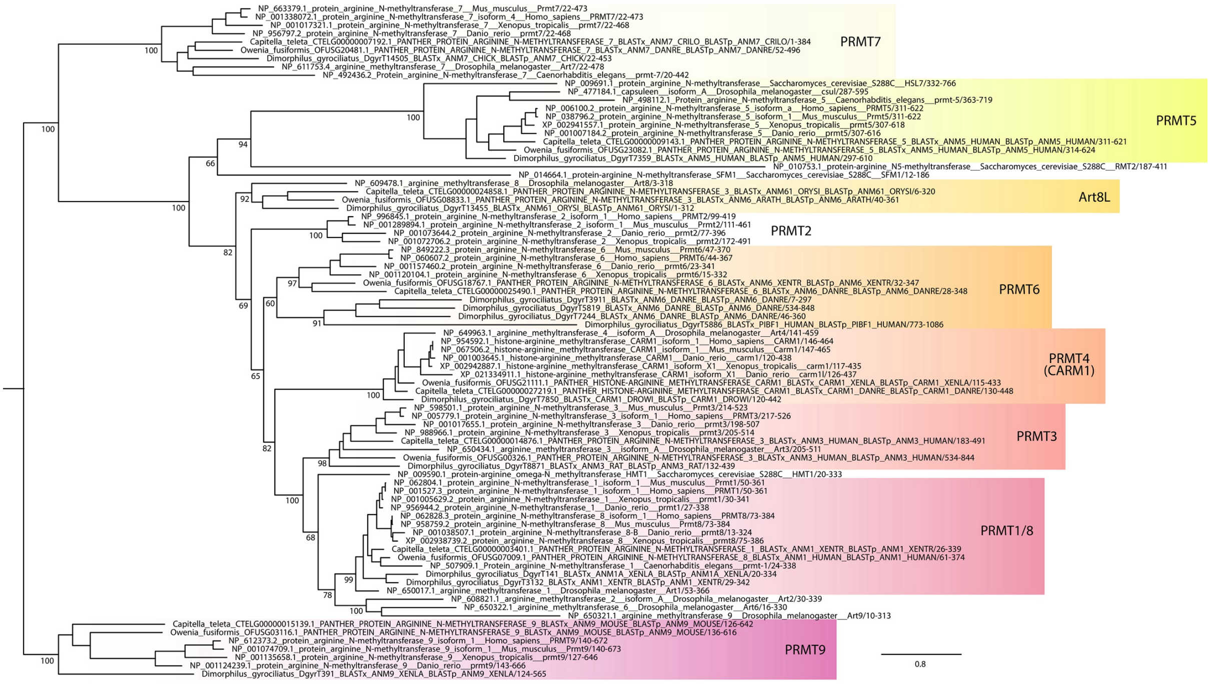


**Supplementary Figure 22 | Maximum likelihood phylogeny of arginine-specific methyltransferases.** Maximum likelihood phylogeny for gene orthology assignment of PRMT genes in *O. fusiformis*, *C. teleta*, and *D. gyrociliatus*. Branch support values represent bootstrap values (0–100 values) at key nodes. Coloured boxes highlight the extent of each PRMT clade. Some protein symbols are custom for annelid or lineage-specific clades, as described in text (e.g., Art8L). Orthologs to more than 1 gene in mammals are assigned as a single one, separated by strokes (e.g., PRMT1/8). Scale bar depicts the number of amino acid changes per site along the branches.

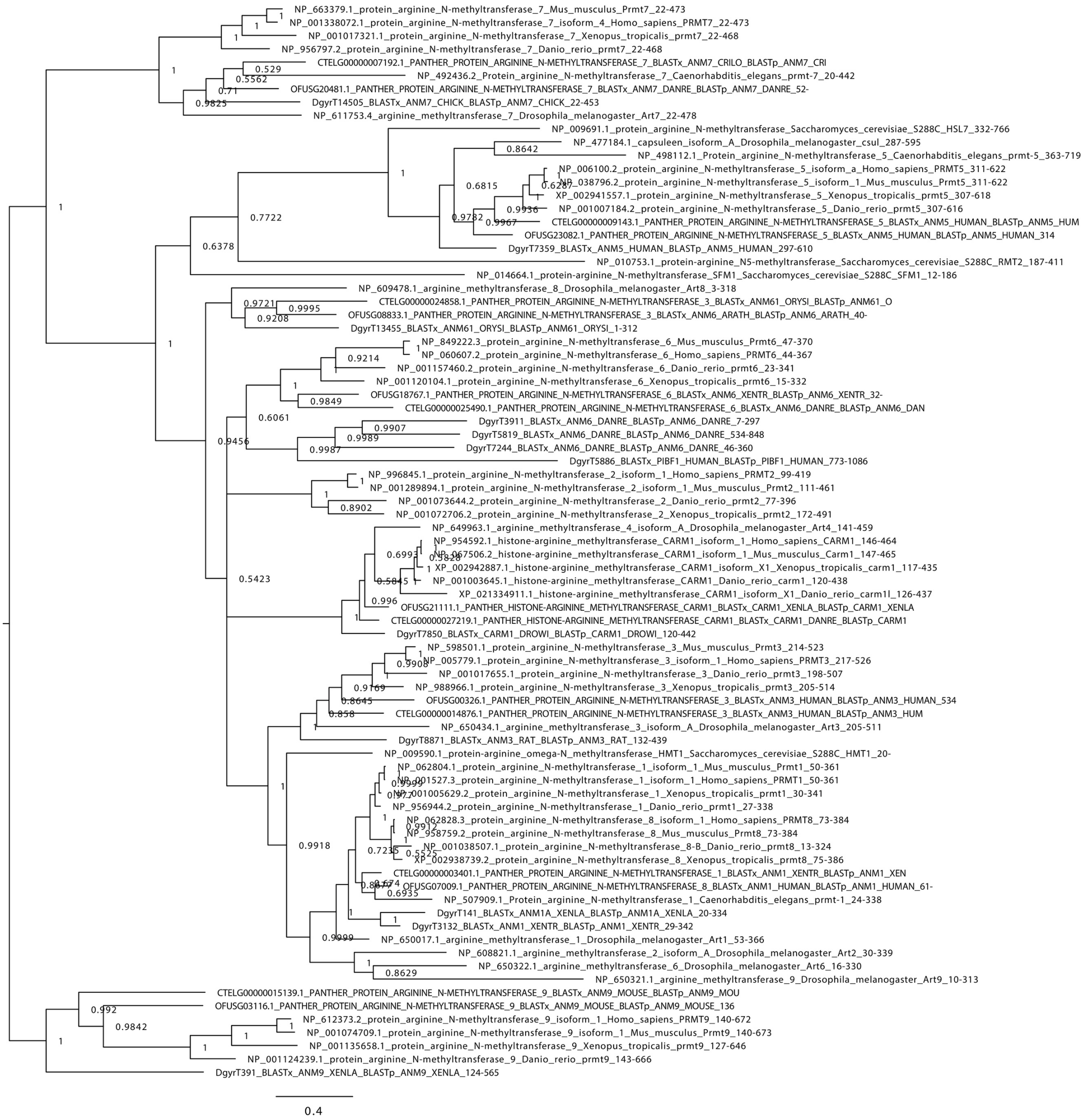


**Supplementary Figure 23 | Bayesian phylogeny of arginine-specific methyltransferases.**

Bayesian phylogeny for gene orthology assignment of PRMT genes in *O. fusiformis*, *C. teleta*, and *D. gyrociliatus*. Branch support values represent posterior probabilities (0–1 values) at each node. Scale bar depicts the number of amino acid changes per site along the branches.

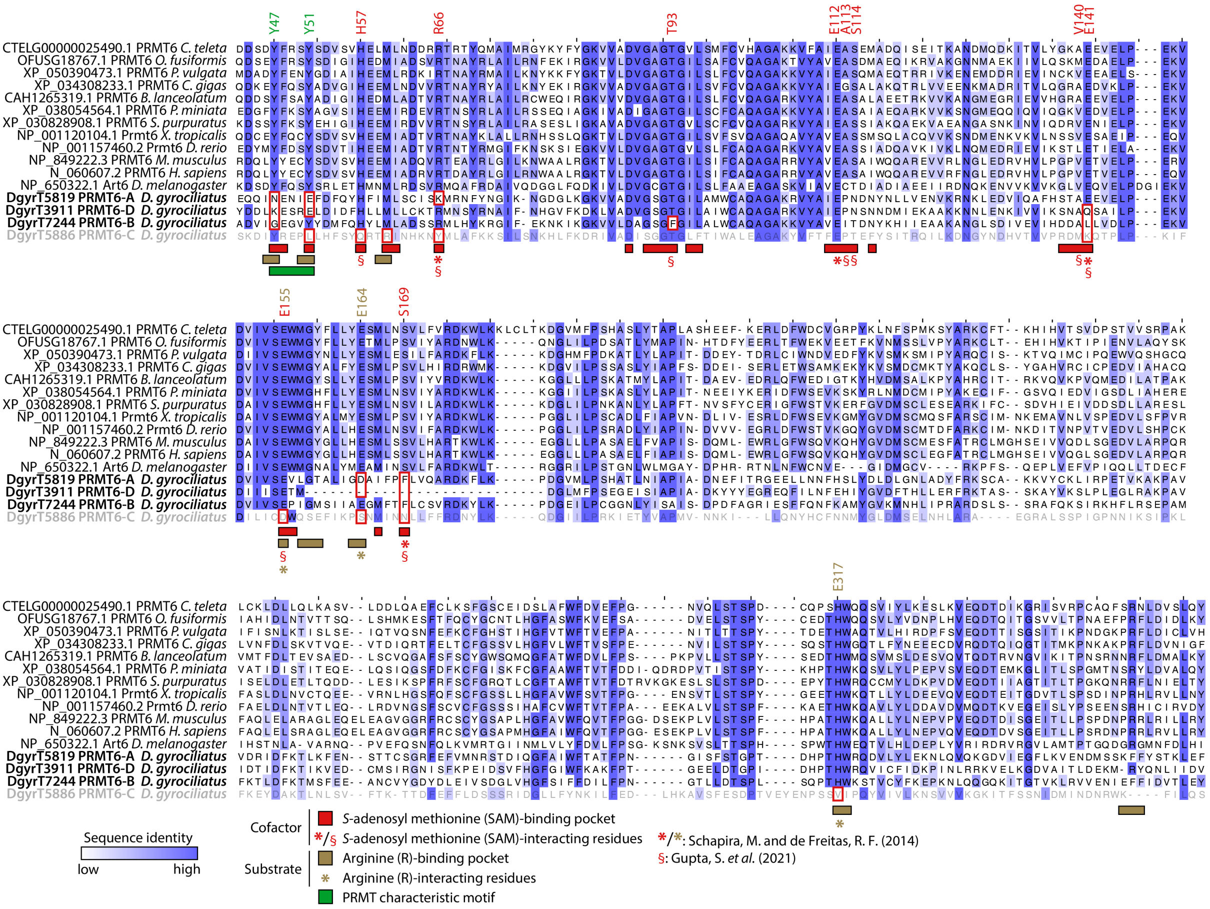


**Supplementary Figure 24 | Sequence diversity in the PRMT6 expansions of *D. gyrociliatus*.**
Full-length MSA of representative PRMT6 sequences, trimmed to the R43–Y359 positions (as per the human PRMT6 nomenclature). At the bottom of the MSA, all four putative PRMT6 orthologs from *D. gyrociliatus* are highlighted in bold. PRMT6-C is greyed out here as well to show its likelihood as an annotation artefact. Key protein regions and residues are highlighted under the MSA. Amino acids with a specified position (as per the human PRMT6 nomenclature) are SAM and arginine-interacting residues. Residues inside red boxes denote conserved positions in key regions or key interacting residues with no conservation in one or more of the orthologs of *D. gyrociliatus*. *X. tropicalis*: *Xenopus tropicalis*. *: SAM-interacting and arginine-interacting residues determined via homology to PRMT4 (CARM1), as in (Schapira and Freitas 2014); §: SAM-interacting residues determined directly in PRMT6, as in (Gupta et al. 2021).

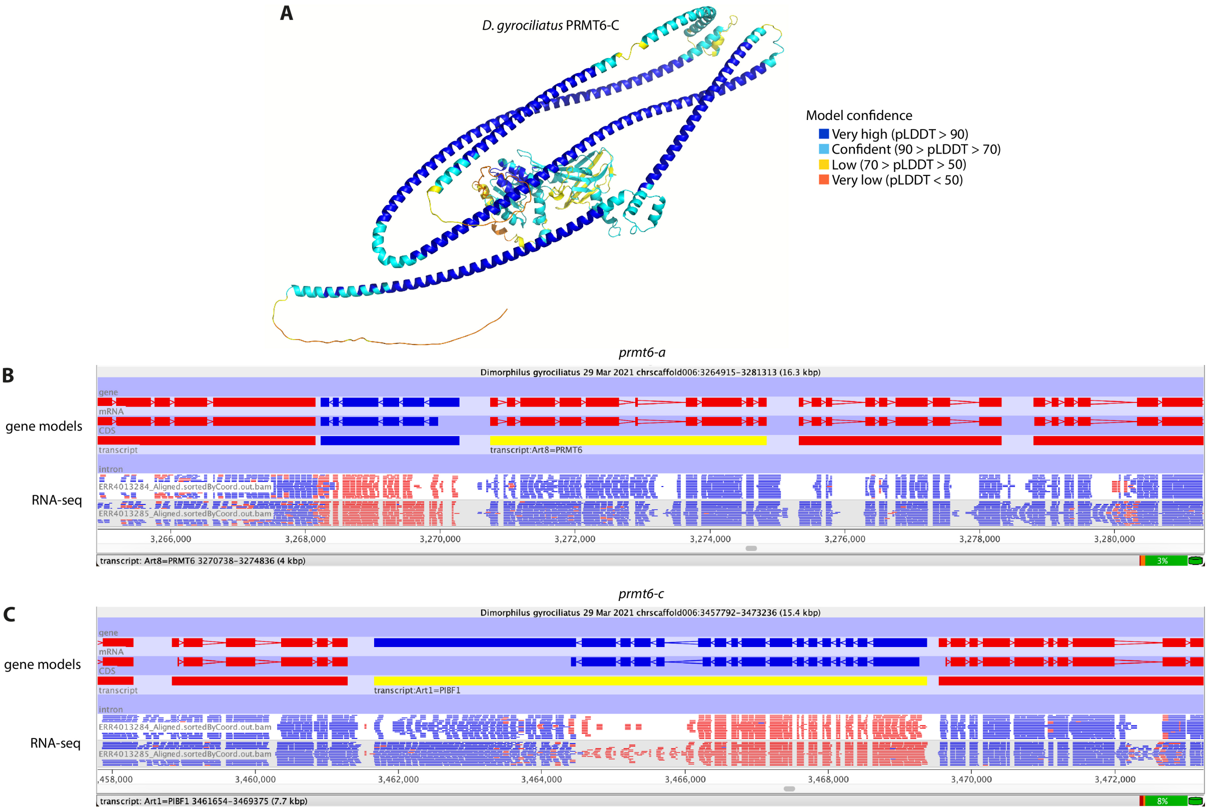


**Supplementary Figure 25 | Domain fusions in the *D. gyrociliatus prmt6-c* gene are likely an artefact.** (**A**) Render of the AlphaFold3 structural predictions of the *D. gyrociliatus* PRMT6-C ortholog. Render colour depicts the model's confidence in the prediction. (**B**, **C**) Seqmonk screenshots of the RNA-seq read density in the late development time point over the *prmt6-a* (**B**) and *prmt6-c* (**C**) gene models. RNA-seq library is opposing strand-specific, meaning that canonical transcription will have the opposite colour as shown above in the gene models track. Antisense reads display the same colour as the gene model. *prmt6-a* and *prmt6-c* gene models are highlighted in yellow with their previously published names. Transcription in the *prmt6-a* gene includes only reads in the expected orientation, homogenously distributed along the gene body and across both fused parts of the gene. This indicates that the gene is most likely indeed a novel fused gene. In the case of *prmt6-c*, the downstream region of the gene, which appears to be non-coding, contains a large chunk of unexpected reads in the opposite orientation. Furthermore, the middle section of the gene, which corresponds with the PRMT6 fraction of the gene, has radically different expression levels than the upstream region of the gene, which corresponds with the progesterone-induced blocking factor 1 family domain, and is much more highly expressed. The lack of continuous transcription suggests these are two different genes (potentially even three when considering the non-coding fraction) that have been misannotated as a single one. Regardless of whether the fusion is artefactual, the PRMT6 fraction is expressed and its sequence is largely divergent from that of other annelids and model organisms.

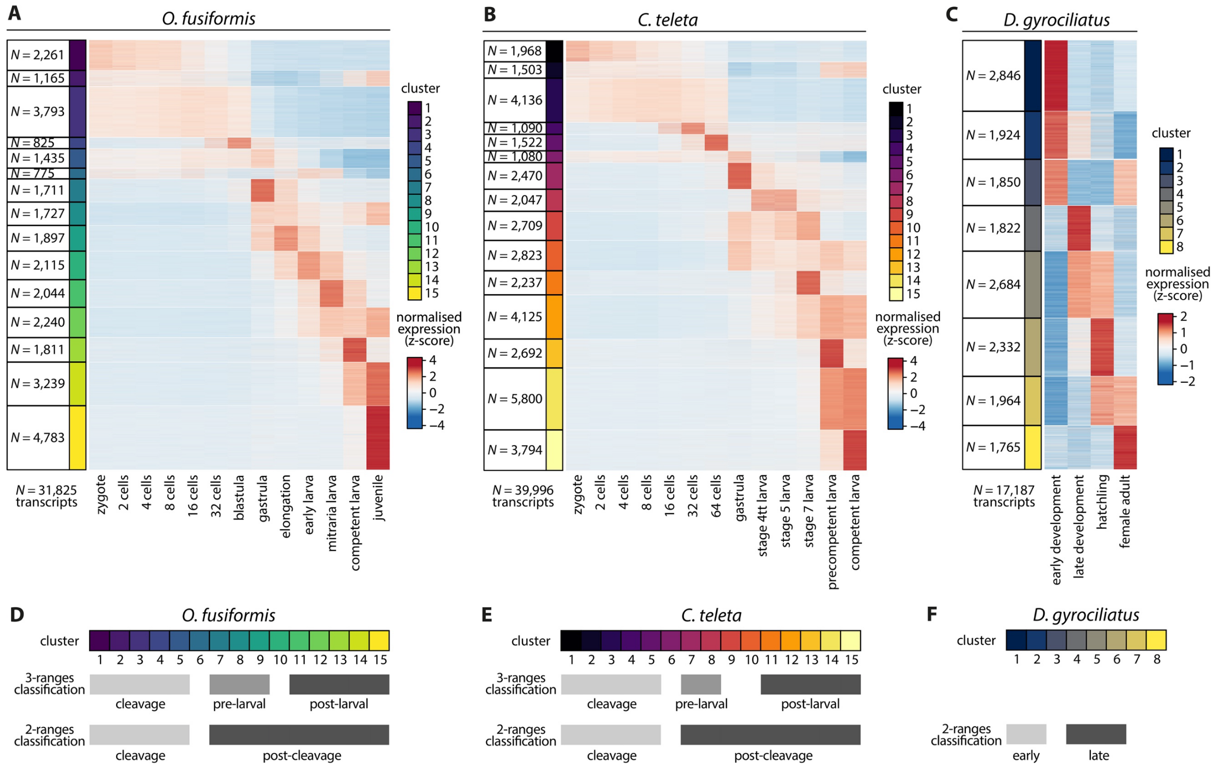


**Supplementary Figure 26 | Transcripts clustering according to full RNA-seq time series.**
(**A–C**) Soft *k*-means clustered heatmap of temporally co-regulated transcripts with a non-null expression in at least one time point into an optimal number of 15 clusters (*O. fusiformis*, **A**; and *C. teleta*, **B**), and 8 clusters (*D. gyrociliatus*, **C**). Next to each colour-coded cluster, *N* denotes the number of transcripts within the cluster. Colour scale denotes normalised gene expression, in a z-score scale. For each species, largest *N* = number of transcripts expressed in at least one developmental stage. (**D–F**) Clusters were classified into a 3- and 2 -ranges classification to perform comparative gene expression analyses, as shown here for *O. fusiformis* (**D**), *C. teleta* (**E**), and *D. gyrociliatus* (**F**).

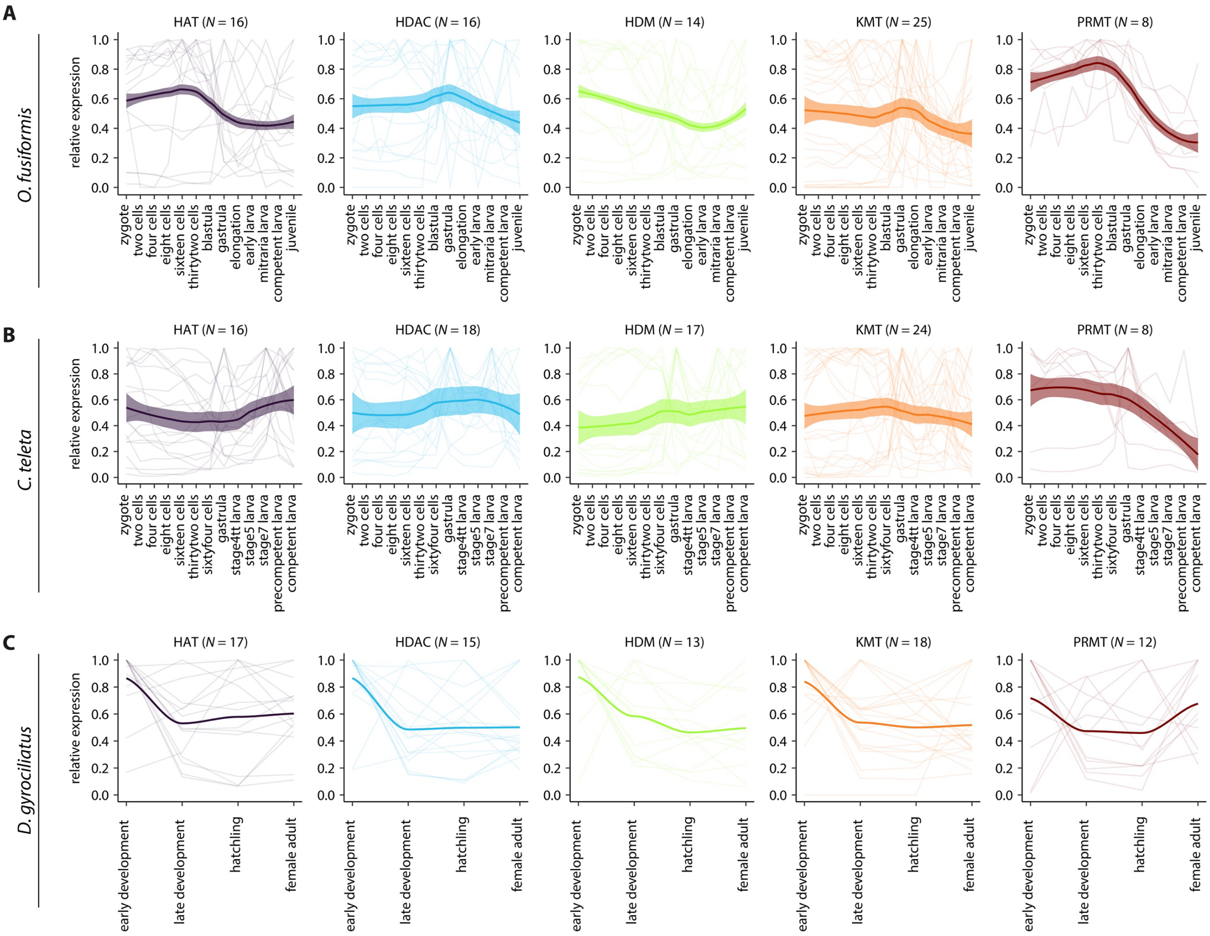


**Supplementary Figure 27 | Family-wise histone modifier expression dynamics in Annelida.**
(**A–C**) Gene-wise relative expression levels (thin background lines) and locally estimated scatterplot smoothings (solid thick lines) for each family of histone modifiers, for *O. fusiformis* (**A**), *C. teleta* (**B**), and *D. gyrociliatus* (**C**). Coloured shaded areas represent standard error of the mean.

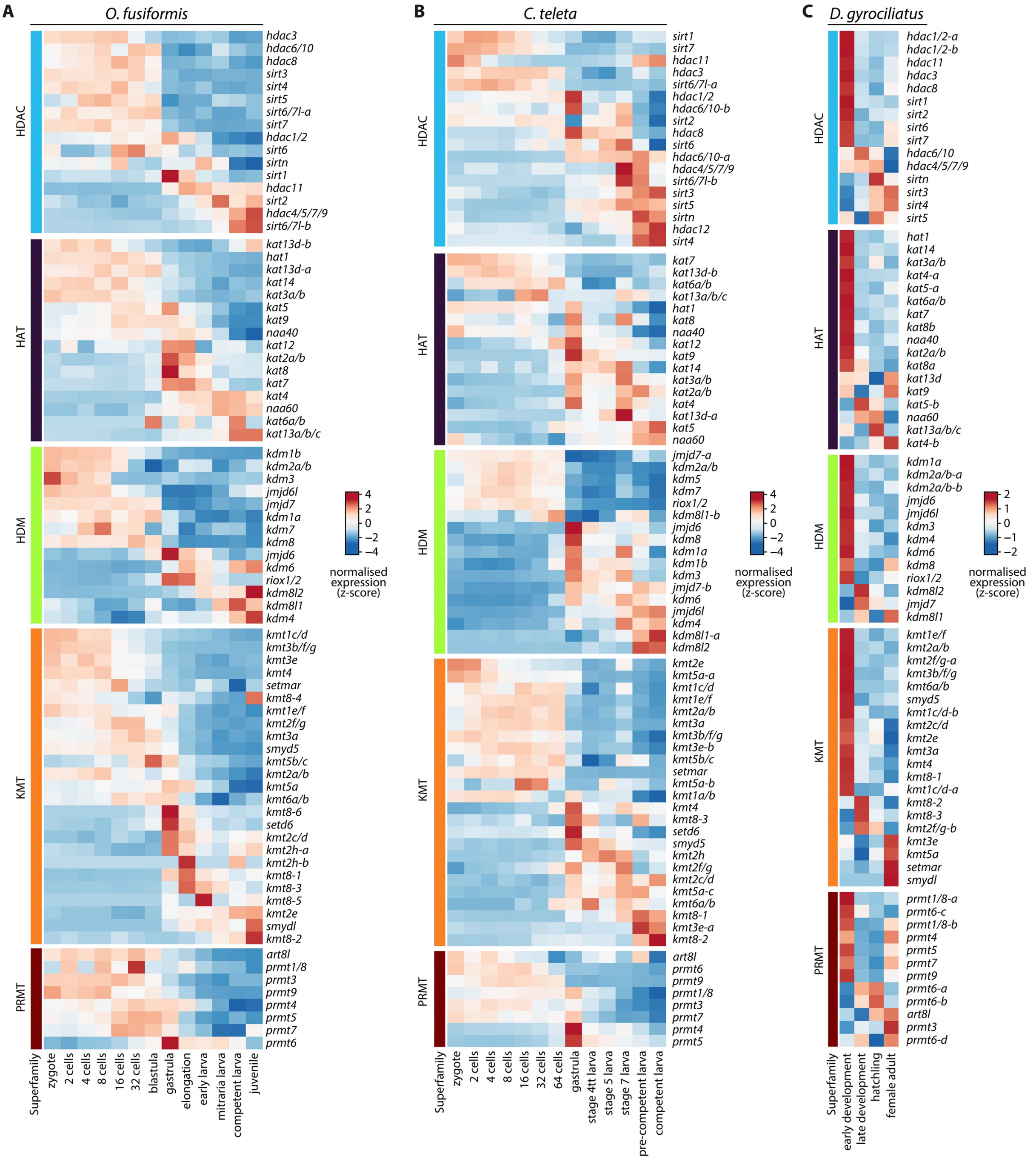


**Supplementary Figure 28 | Histone modifiers expression dynamics in Annelida.**
(**A–C**) Soft *k*-means clustered heatmaps (as in Supplementary Fig. 26) of gene expression dynamics of histone modifier genes across the development of *O. fusiformis* (**A**), *C. teleta* (**B**), and *D. gyrociliatus* (**C**), classified by family of histone modifiers. Colour scale denotes normalised gene expression, in a z-score scale.

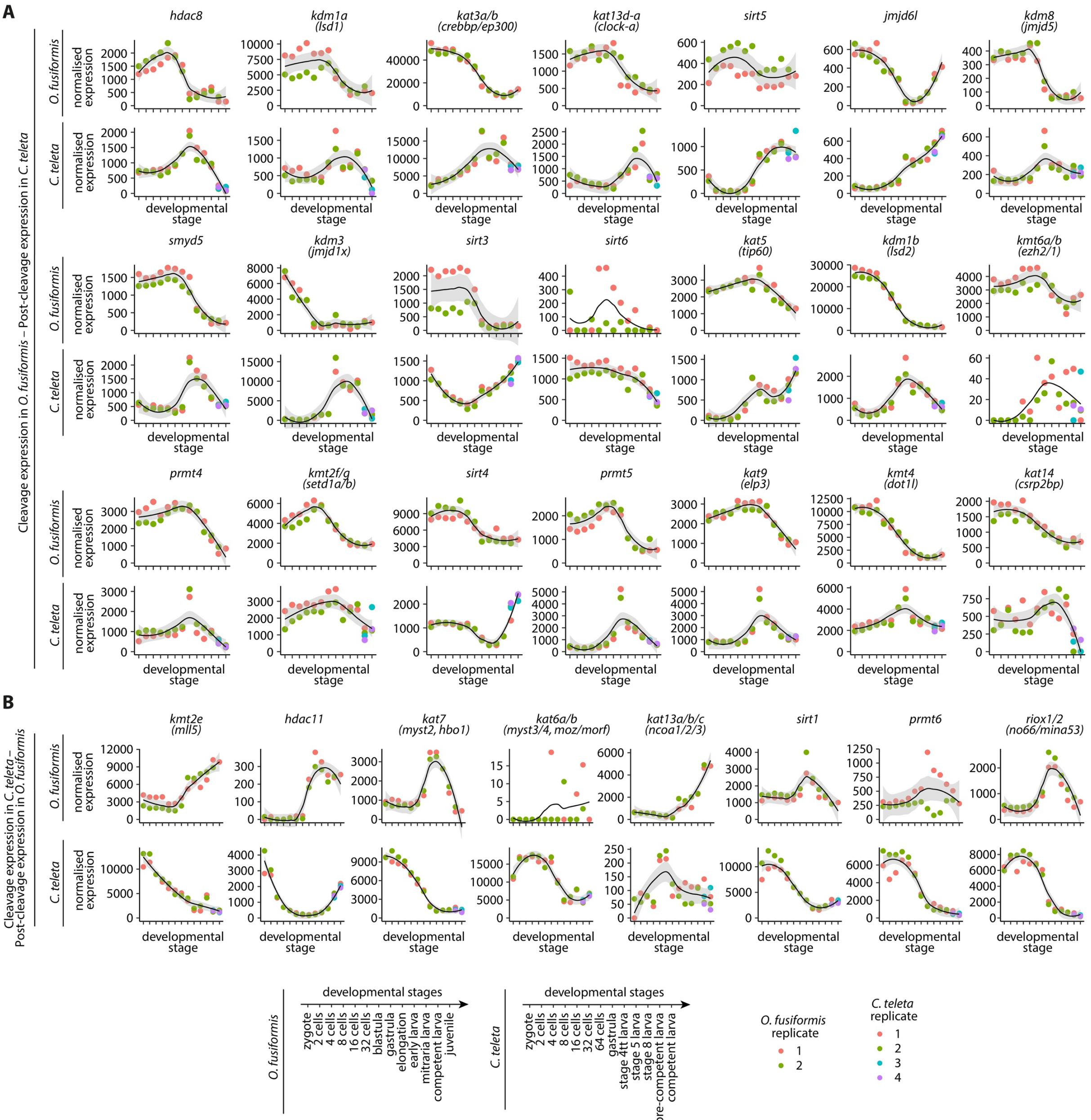

**Supplementary Figure 29 | Gene expression levels of heterochronic histone modifiers correlated with larval type.**

(**A**) Normalised expression levels of histone modifier genes under heterochronic shift between cleavage expression in *O. fusiformis* and post-cleavage expression in *C. teleta*, during the development of *O. fusiformis* (top) and *C. teleta* (bottom). (**B**) Normalised expression levels of histone modifier genes under heterochronic shift between cleavage expression in *C. teleta* and post-cleavage expression in *O. fusiformis*, during the development of *O. fusiformis* (top) and *C. teleta* (bottom). Curves in **A** and **B** are locally estimated scatterplot smoothings, coloured shaded areas represent standard error of the mean. Time points are summarised on the bottom for both RNA-seq time series.

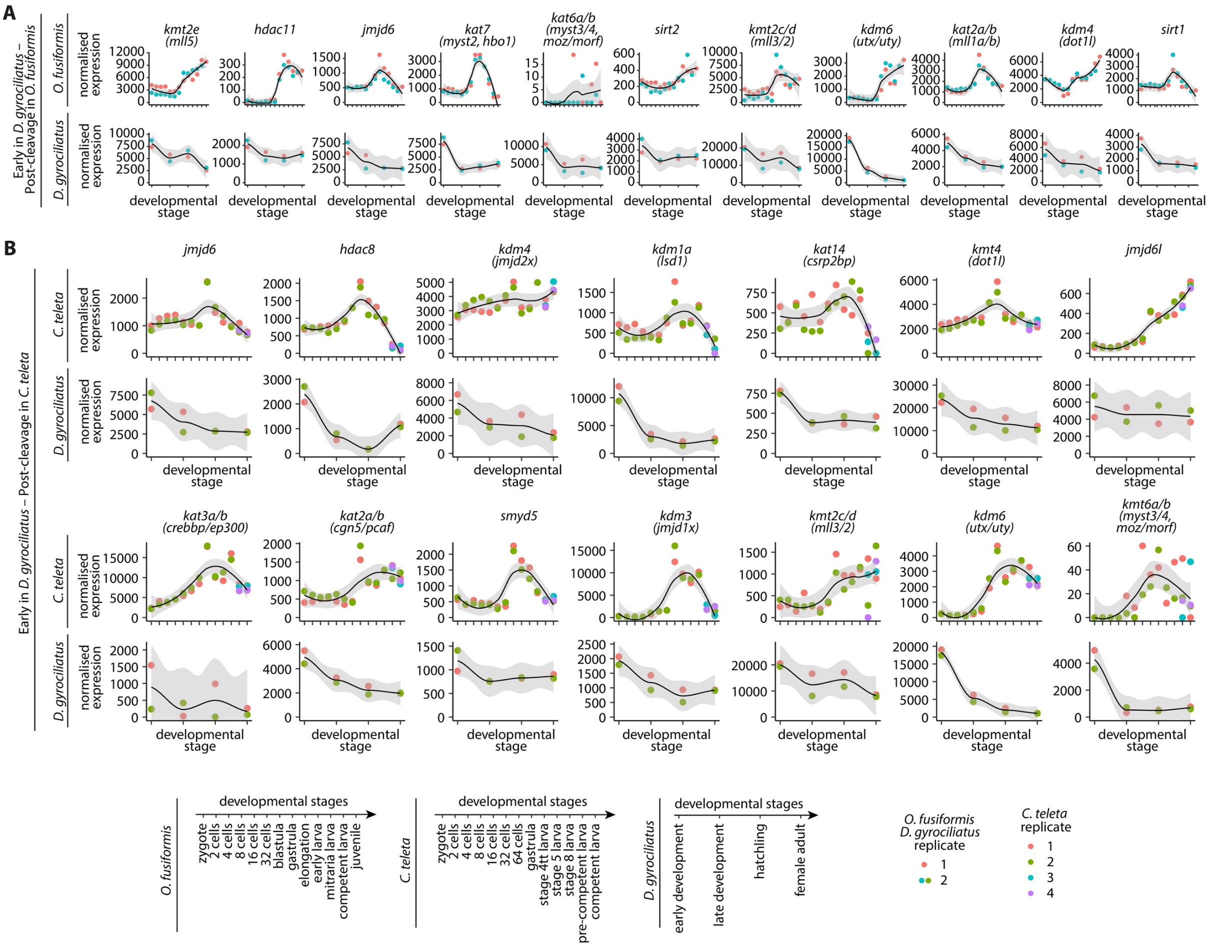


**Supplementary Figure 30 | Gene expression levels of heterochronic histone modifiers correlated with life cycle.**

(**A**) Normalised expression levels of histone modifier genes under heterochronic shift between early expression in *D. gyrociliatus* and post-cleavage expression in *O. fusiformis*, during the development of *O. fusiformis* (top) and *D. gyrociliatus* (bottom). (**B**) Normalised expression levels of histone modifier genes under heterochronic shift between early expression in *D. gyrociliatus* and post-cleavage expression in *C. teleta*, during the development of *C. teleta* (top) and *D. gyrociliatus* (bottom). Curves in **A** and **B** are locally estimated scatterplot smoothings, coloured shaded areas represent standard error of the mean. Time points are summarised at the bottom for all three RNA-seq time series.

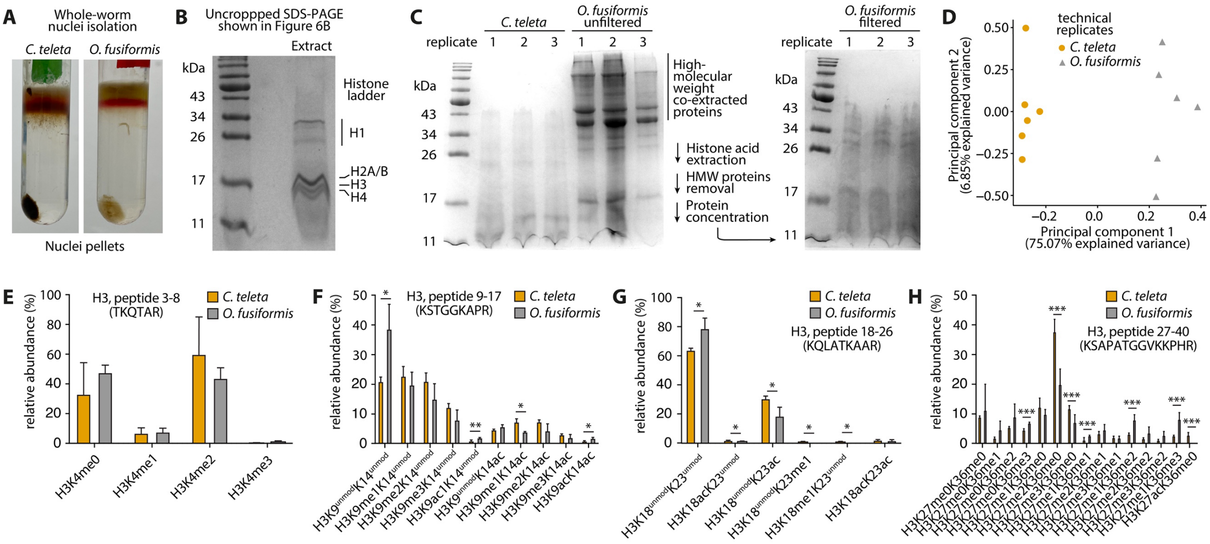


**Supplementary Figure 31 | LC-MS/MS hPTM quantification in acid-extracted histones from adult annelids.**

(**A**) Nuclei isolated from *C. teleta* (left) and *O. fusiformis* (right) through rate-zonal centrifugation in a sucrose solution of the raw whole-worm lysates, coming from 15 and 3 specimens, for *C. teleta* and *O. fusiformis*, respectively. (**B**) Uncropped SDS-PAGE shown in Fig. 8B, depicting the traditional histone ladder observed in acid-extracted histone samples. (**C**) SDS-PAGE analysis comparing acid-extracted histone samples from *C. teleta* and *O. fusiformis* (left gel). *O. fusiformis* histones co-purify with unidentified high-molecular weight (HMW) proteins. We therefore included an additional cleaning step with a 30 kDa NMWL ultrafiltration device to remove these HMW proteins. Resulting samples (right gel) show a successful filtration step. (**D**) Principal component analysis of the histone H3 and H4 hPTM profiles derived from LC-MS/MS experiments, by technical replicate, for all analysed samples of *O. fusiformis* and *C. teleta*. (**E–H**) Relative abundance bar plots of the H3 3–8 peptide (TKQTAR) based on H3K4 methylation status (**E**), the H3 9–17 peptide (KSTGGKAPR) based on H3K9 methylation or acetylation and H3K14 acetylation status (**F**), the H3 18–26 peptide (KQLATKAAR) based on H3K18 methylation or acetylation and H3K23 methylation or acetylation status (**G**), and the H3 27–40 peptide (KSAPATGGVKKPHR) based on H3K26 methylation or acetylation and H3K36 methylation status, in *O. fusiformis* and *C. teleta*. Error bars in **E–H** represent standard deviation. *P* values were derived from two-tailed Student's *t*-tests. *: *P* value < 0.05; **: *P* value < 0.01; ***: *P* value < 0.001; otherwise not significant.
